# Planning and Conducting an Online Conference at the time of COVID-19: Lessons Learned from EGREPA 2021

**DOI:** 10.1101/2021.09.19.460976

**Authors:** Ellen Bentlage, Luca Roekens, Lennart Bußmann, Michael Brach

**Affiliations:** Institute of Sport and Exercise Sciences, University of Münster, Horstmarer Landweg 62b, 48149 Münster, Germany; University of Münster (Students), 48149 Münster, Germany

**Keywords:** Online Conference, Indico, YouTube, OBS, Zoom, Gather, COVID-19, Pandemic

## Abstract

**Background:** This paper describes the realisation and evaluation of the EGREPA 2021 conference, in online format. As an example, and support for future conferences, the decision process to switch during the COVID-19 pandemic from onsite to online, the conception and tools, the experiences of the organisers and the feedback of the participants are described.

**Methods:** The CERN tool “Indico” served as the central conference management system including public homepage, submission and payment system, abstract review and decision platform, and conference material repository. Zoom, YouTube Live and Open Broadcaster Software (OBS) were utilised for keynotes and other presentation sessions, including livestreaming of pre-recorded presentations. Gather was implemented for poster sessions and individual meetings. An anonymous survey included Likert-type questions on acceptance and quality of the tools and instructions, open questions related to positive and negative issues, and ideas for the next online-conference.

**Results:** The event was evaluated well, with mostly fluently working technical solutions. A few participants indicated problems switching from YouTube to Zoom between presentations and discussions. Opinions on pre-recordings of talks were divided. Participants required more time for socialising. Guidelines and materials were mostly rated helpful.

**Lessons Learned:** Simplifying the livestreaming, presenter’s choice between pre-recording or presenting live, more time for socialising and breaks are discussed. Tools for conducting an online conference should be introduced to the participants in advance. Indico is rated as a very helpful tool for planning and conducting an online-conference.

## 1. Introduction

The European Group for Research into Elderly and Physical Activity (EGREPA) [1] has organised seven conferences over the last 15 years, each one of them usually attracts between 100-300 participants. The latest conference which took place in May 2021, was jointly organised by the University of Münster (WWU), the University of Physical Education in Krakow, and European Advisors’ Association PlinEU under the topic “Active Aging – New Challenges, New Opportunities – Bridging Research & Practice”. Beyond the typical components such as keynote lectures, invited symposia, short talks and posters sessions, the conference programme included physical activity choices and project demonstrations. Due to the fluctuating pandemic situation in the European countries, the seventh EGREPA conference was held virtually.

The purpose of this paper is to describe the decision making in face of the fluctuating pandemic situation in European countries (section 2), the conception and tools for the online format of this conference (section 3), the experiences of the organisers and the feedback of the participants (section 4), in order to present a useful and practical example and to discuss lessons learnt for future conferences, with or without pandemic situations (section 5 and 6).

## 2. Online conference: Decision making and examples

Initially, the conference had been planned to take place in Krakow, Poland, between 19-21 May 2021 (Plan A). Due to the COVID-19 pandemic situation, and the associated national and international measures that restrict the in-person group meetings, several online meetings with the conference committee took place between October 2020 and March 2021, in order to discuss the format and the tools of the conference. Four options were considered: Postponing the conference (Plan B), “hybrid” with on-site and online participants (Plan C) and online only (Plan D). In case of postponing, we would risk losing most of the organisational work, as presenters, symposia organisers and keynote lecturers would have to be asked for availability again. In addition, there would be no guarantee for a significant decrease of the pandemic situation in the host and participants countries. While the different local and national traveling restrictions had brought up the hybrid format, it could not be realised due to the increased COVID-19 incidences two months before the start of the conference in Krakow. Consequently, the University of Physical Education in Krakow was not able to welcome external participants. Therefore, on the 18^th^ of March 2021 it was decided to switch the conference from onsite to online (Plan D).

In order to review experiences with tools for livestreaming and video-conferencing, different actions were made:

- A literature review of best practices was carefully screened,
- Informal request between academic peers were exchanged,
- Reports and blogs discussing online conferences, that had already been held, were checked.

The results of the literature review [2] showed that different tools are specified by the number of maximum users, costs, media support, recording and replay functions. For live-streaming pre-recorded presentations, Twitch [3], Adobe Connect [4], Zoom Webinar [5], Crowdcas [6], Intrado [7], YouTube Live [8] and Vimeo live events [9] were mentioned. For video-conferencing, the platforms Zoom [10], Webex [11], GoToMeeting [12], Dialpad Meetings [13], Blujeans Meetings [14], GlobalMeet [15], Google meet [16], Jitsi Meet [17], Skype [18], and Microsoft Teams [19] were reported.

Informal requests showed that an online conference with local attendees was held via Zoom. In addition, good experiences with YouTube Live were shared.

Results from reports and blogs gave technical insights about three already conducted conferences: The Photonics Online Meetup (POM) 2020, with 3 live sessions of 1,5 hours each via the video platform WebEx [20]. Questions from the audience were raised via the WebEx chat, session chairs forwarded the questions to the speakers. Talks were recorded and made available for two weeks. As primary communication platform, Slack was integrated for participants and the organisers. Twitter was used for the poster session. The organisers generated a poster template, which was optimised for the twitter format. Presenters were asked to create a personal or group account in order to insert their posters with a short description. Comments were publicity visible below each poster. A website served as a central information repository. In addition, the submission of abstracts and the registration for the event was performed with Qualtrics and Google Forms. For the event was free, no payment process was considered [20]. The feedback gathered on the POM 2020 was mainly regarding the interaction during the sessions, and how it would be better if the audience can ask questions directly via audio- or video-connection. Additionally, Twitter was not accessible in some regions. One year later, the poster session of the POM was held via the virtual conference space Gather, which allowed participants to walk around with an own avatar, view a poster and interact with the presenter [21]. The Virtual Remote Future Summit in 2019 was a two-day event. Around 80 percent held their presentations live (keynotes and those with experience in holding presentations). The other 20 percent (speaker who never presented on camera) recorded their presentations in advance. The tool Teachable was used for uploading pre-recorded video talks and other resources. For delivering live presentations, YouTube Live was used and Slido acted as the real-time communication platform for collecting questions during the presentations in the chat. All attendees were able to vote for the ones they liked. Then, speakers answered them in real time. Organisers indicated, that it is important, to introduce the tools in advance to the audience. [22].

## 2. Methods

In this section we describe the technical and practical conception of the seventh EGREPA conference. In order to avoid extra costs for the attendees, the conference committee refused to hire a video-conference company. Instead, WWU academics were mainly responsible for the organisation and digital implementation., benefiting from their online teaching experience, developed in the pandemic time. Furthermore, the conference was supported by the WWU technical staff and tools.

For recently, the WWU had established an own installation of Indico, a well-known conference management system, which operated as the central homepage for participants and main tool for organisers (2.1). In order to allow a high technical standard, independent from presenters’ local internet connections, all presentations (except the keynotes) were pre-recorded and streamed via YouTube Live. The sequence of the live played videos was prepared in advance with the encoder Open Broadcaster Software (OBS) (2.2). For video-conferencing, the platform Zoom was utilised, including live presentations and discussions of the two keynote lectures, discussions after each pre-recorded presentation, and all moderation activities (2.3). The poster sessions took place in the virtual conference space in Gather, which was mainly integrated for socialising (2.4). For the internal team communication during the conference, WhatsApp was utilised (2.5).

While the technical tools are sketched in this section, the procedures during the conference sessions are described in the additional file 1.

### 2.1. Conference management and website with Indico

Indico is an open-source tool for organising lectures, meetings, and conferences [23], that started as a European project in 2002. Since 2004, CERN financed its continuous development and use as own event-management solution [24]. The integrated manual on the official website supports an easy usage of the Indico platform, because it combines e-learning videos and a written user-guide [25]. The WWU has established an own instance of this system, enabling WWU researchers to create and manage events, sharing different access levels for organisers, managers, reviewers, and active and passive participants. By adding data (e.g. dates, topics, fees), promotional material (logo, image, call texts) and instructions (for participants, speakers, reviewers), the “external” conference website and “internal” management area emerge at the same time. Updates are directly visible without need for webpage expert knowledge. Information flow to the public and to specific user groups (roles) were easily retrieved. All typical management processes like abstract submission, review and decision, setting up the programme, building the book of abstracts, registration and payments are integrated in the system. For the abstract review process, three user roles (organiser, reviewer, judge), four criteria (originality, quality of writing, scientific rigor, topic fit) and three outputs (accept, resubmit, change of contribution type, reject) were established. Decisions were automatically sent to the authors. The official call for abstracts ended on 20^th^ of January 2021. Due to popular demand, the committee decided to open up the opportunity for ‘late breaking’ poster presentations from 12^th^ until 25^th^ of April 2021. For registering to the conference, participants could choose between payment per credit card or per invoice. Indico had been internally connected to the university ERP system, this helped to conduct any financial transaction according to the university procedures. Alternatively, an external bank account could be used. A free registration was offered for EGREPA members, as well as for the EGREPA Board, the Scientific Committee and Invited symposia speakers. All participants who had registered were integrated into the official conference programme. The book of abstracts, different links for attending the conference and pre-recorded presentations were uploaded on the Indico website. The protected mode was enabled, in order to limit the access only to the registered participants. As Indico doesn’t allow for data processing permission during the registration process, we created a Google Form [26] and sent it to all presenters. In addition, the evaluation questionnaire was conducted via Indico (section 4).

### 2.2. YouTube Live/ Open Broadcaster Software (OBS)

The video platform YouTube, founded in 2005, offers users to upload or live-stream video and make them available with or without restrictions for free [27]. Complex livestreams, like running large conferences, can be planned in advance with encoder software, which converts the media into a suitable digital format for livestreaming to the audience [28]. For the EGREPA online conference, YouTube Live was combined with Open Broadcaster Software (OBS), a free, easy to handle, open-source software for encoding. Based upon the official programme schedule, the pre-recorded talks and other audio and video content were integrated into OBS as different scenes. During the conference, the different scenes were selected and streamed live on YouTube, with unlimited number of viewers. Reminders to switch back to Zoom or break announcement were placed at the end of each scene. After the event, YouTube generated an overview of the number of participants who accessed the livestream over time. In addition, the maximum number of participants who simultaneously watched the presentations could be accessed.

### 2.3. Zoom

Zoom, founded in 2011, is a pioneer in the field of modern cloud platform for video and audio conferencing [29]. In order to guarantee a high quality of the video and audio, a license to upgrade the number of participants to 500 had been issued via WWU IT staff. Keynote live presentations, as well as discussions after each presentation took place in Zoom. Support from one assistant was continuously necessary, e.g. muting participants while watching presentations on YouTube, allowing keynote lecturers to share their screen, supporting the session chairs during announcements and discussions (who raised the virtual hand or wrote into the chat). Reminder to switch to YouTube in order to attend the next presentation and other organisational information was shared via the screen. After the event, a list of conference attendees could be downloaded from Zoom, where the total number of participants, the start- and end-time of each user, and the total attendance duration were visible.

### 2.4. Gather

The platform Gather, founded in 2020, combines video-calling with a 2D map, in which the user can walk around as an avatar and interact with others [30]. Like in reality, users can see and hear each other (by video-call), when their avatars are in the near. The administrator can select from default 2D maps and/or built own environments. The conference space in Gather was built two months before the event started. Three main areas were created: The main conference hall with open spaces and private rooms, two poster session rooms for poster discussions, and a lounge with a warm atmosphere for private conversations. The virtual conference Gather space can be still accessed [31]. Gather was mainly used for poster discussions in smaller groups, which took place after all posters of each session were presented in YouTube Live. Like a usual poster session, the posters were visible in the Gather room, and presenters were available for discussions near the poster. Participants walked around and could discuss in front of the poster in a private-space in a video-call. Gather offers a limited free feature (about 25 users). For extended features and stable video-conferencing for more users, a fee based upon the event time was charged [32].

### 2.5. WhatsApp

WhatsApp (founded in 2009), guaranteed a fast and safe communication among the conference team, independently from local LAN / WLAN [33]. It was chosen, because the pandemic regulation required different rooms and not all staff was present in the institute building. The messenger also worked for spontaneous requests or technical problems of the participants.

## 3. Conference Materials

The term “conference materials” refers to the section in the Indico system, where files can be shared with any user.

### 3.1. Topical materials

The official conference programme, including authors and titles of each presentation was shared with all participants two months before the event started (additional file 2). The digital book of abstract was uploaded one day in advance (additional file 3). The pre-recorded videos were available for all participant at the end of each conference day, and for one week after the end of the conference.

### 3.2. Training materials

Included instructions for presenters on how to prepare the pre-recording of the presentation, a checklist and practical tips (additional file 4), a user-guide how to record with power-point (additional file 5) and how to upload the presentation into Indico (additional file 6) were generated. In addition, a video to introduce Gather and a handout (additional file 7) were produced for participants. These materials were shared per email, and additionally uploaded to the central homepage.

## 4. Evaluation of the Online Conference

In order to evaluate the online conference, an anonymous survey was created and launched in Indico system. Participants were asked to complete the survey up to three weeks after the conference ended.

### 4.1. Survey Items

An overall rating (Q1) and questions related to the technical components (Q2-Q13) were structured as a five-point Likert Scale. Open-ended questions asked about the positive and negative aspects (Q14, Q15) and about improvement suggestions (Q16). Figure 1, figure 2, and the additional file 8 provide details.

**Fig. 1.**
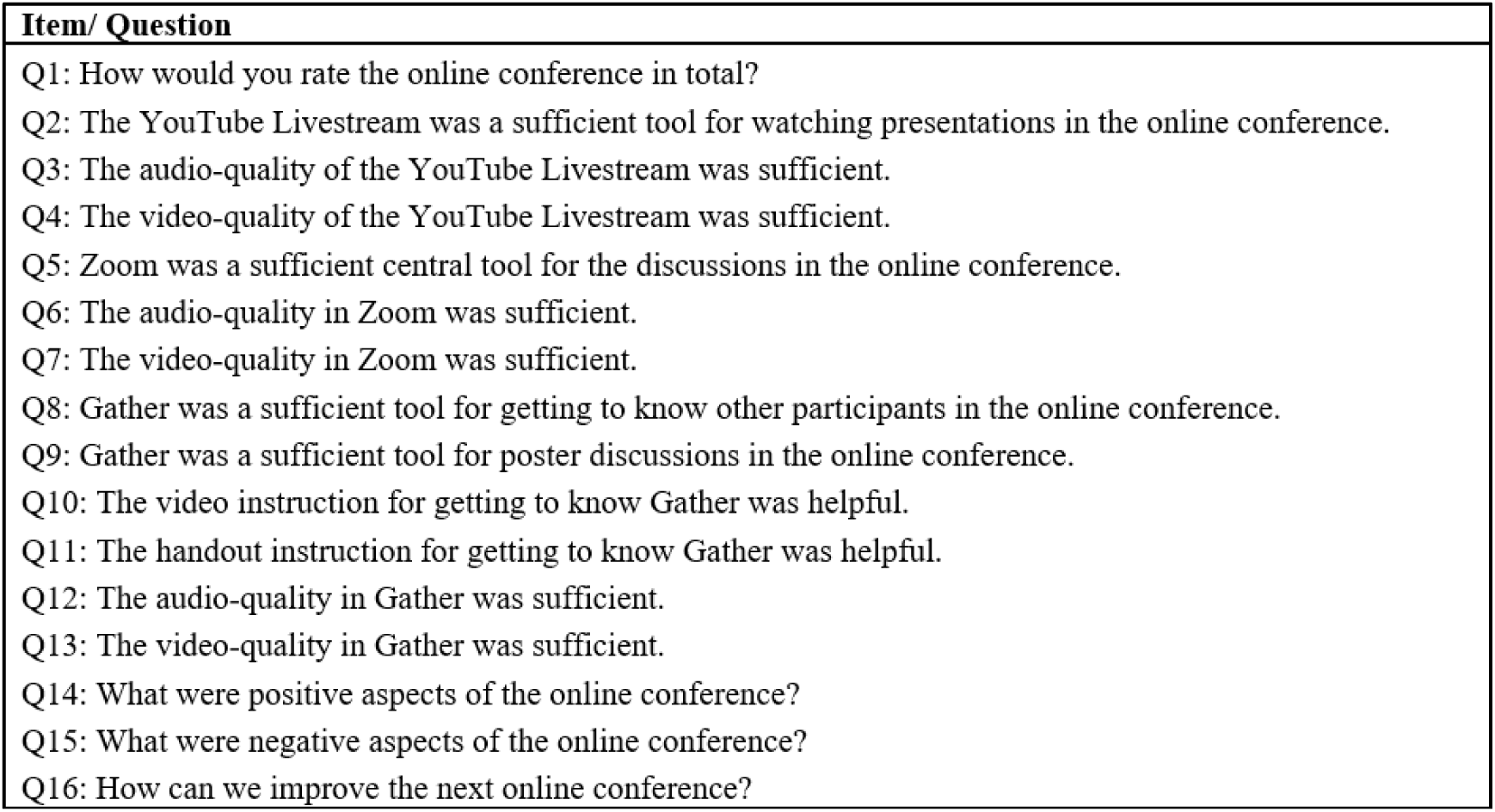
Questions of the survey

**Fig. 2.**
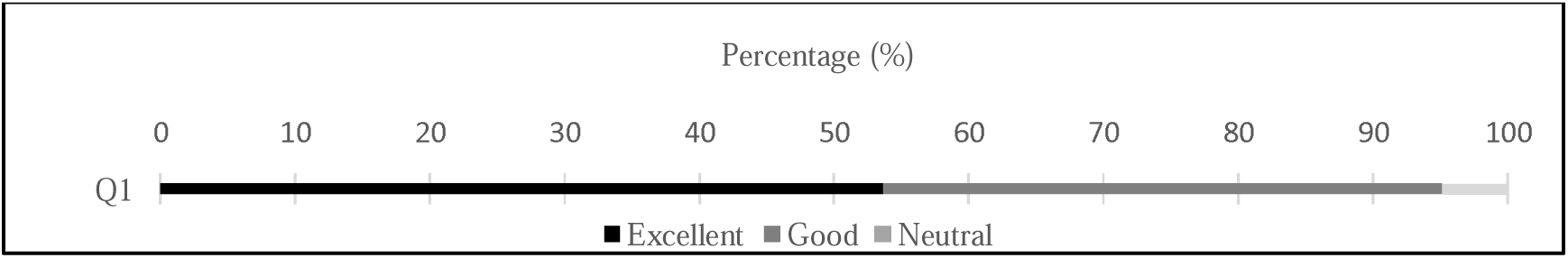
Results - Q1

**Fig. 3.**
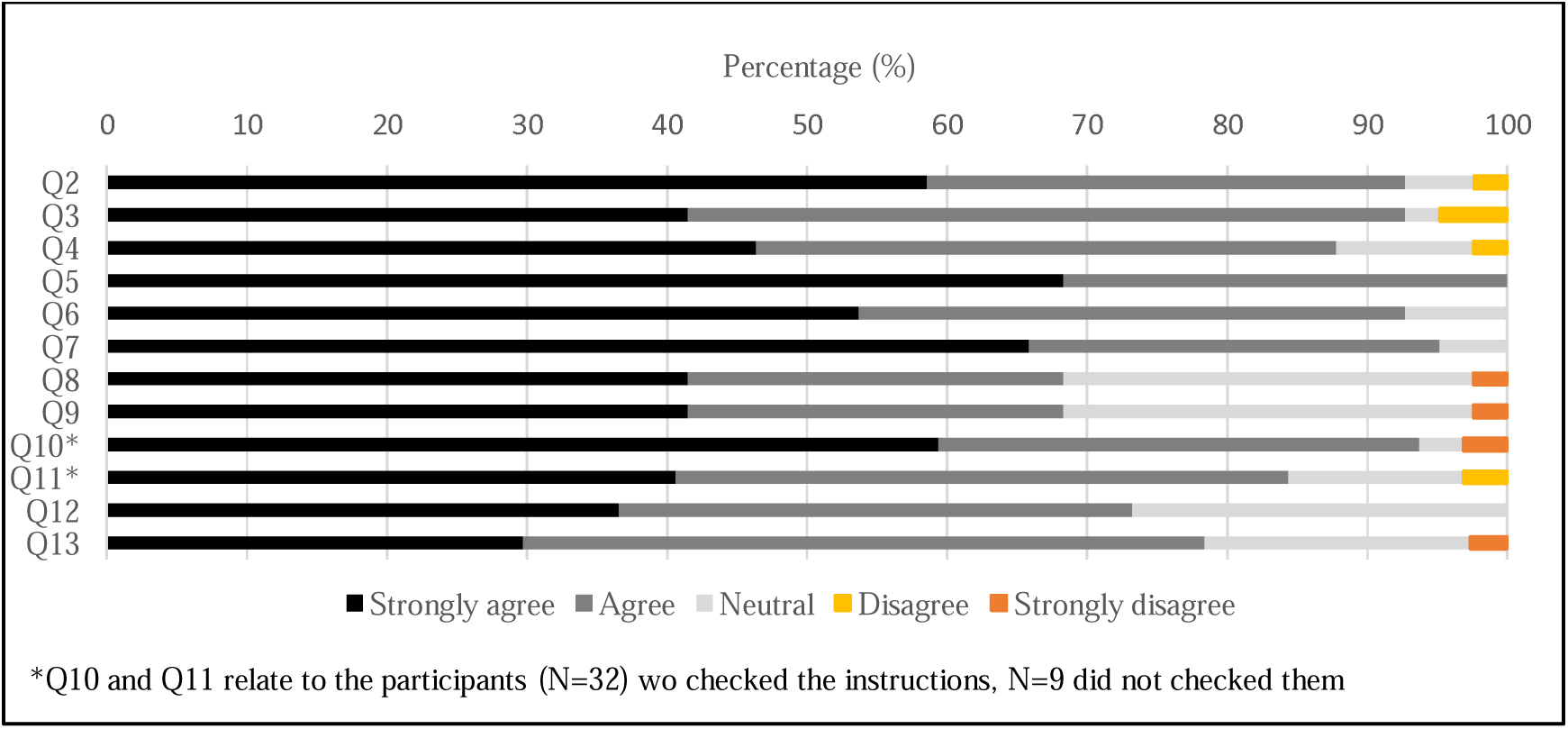
Results – Q2 until Q13

### 4.2. Results

During the three-day online conference, about 55 out of 132 registered participants from 19 countries simultaneously attended the presentations. Overall, 41 participants (response rate 31%) participated in the survey.

More than 95% rated the online conference overall “excellent” or “good”. Nearly 5% evaluated “neutral”, none rated “poor” or “very poor” (Q1). The YouTube-Livestream was a sufficient tool for watching presentations (Q2), with sufficient audio and video quality for more than 85% (Q3, Q4). Zoom as conference tool for the discussions reached even higher rates in general (Q5), and regarding a sufficient audio quality more than 90% agreed or strongly agreed; only 7% rated it as “neutral” (Q6). Regarding the video quality around 95% agreed or strongly agreed, 5% were neutral (Q7). As Gather was considered new tool to most of the organisers and participants, we asked more specific questions on it. The purposes of the tool, getting to know each other (Q8) and conducting the poster session (Q9), were rated identically, with majority agreeing/ strongly agreeing (almost 70%). 29% of the participants rated it as “neutral” and (2,4%) strongly disagreed.

More than 80% found the Video (Q10) and written instruction (Q11) helpful. For the video explanation, 3,1% evaluates it as “neutral” and 3,1% as not helpful. Few participants rated the usefulness of the written instruction “neutral” (12,5%), or rejected it (3,1%). Gather’s video and audio quality (Q12, Q13) was sufficient for more than 74% (agree/ strongly agree), with only 2,7% strongly disagreed.

Answers to the open-ended question regarding the positive aspects (N=29, Q14), negative aspects (N=29, Q15) and improvements of the online conference (N=19, Q16) are visible in the additional file 8. The main findings can be summarised into scientific, technical, social and organisational aspects:

- **Scientific aspects** The high quality of research, diverse projects and constructive discussions were highly appreciated. In summary, the conference was evaluated as interesting and enlightening. None of the participants referred to negative aspects or suggestions for improvements regarding scientific aspects.
- **Technical aspects** With regard to technical aspects the extremely well-run program was highlighted, especially the transition between the different tools (YouTube, Zoom and Gather), which worked well in most of the cases, after a short familiarisation period. However, some participants recommended to reduce the number of the tools used. There were divided opinions regarding the pre-recordings. While some participants preferred live-presentations, others valued the stress limitation with doing the pre-recordings. In future, participants could be motivated to turn on their video, when interacting with others.
- **Social aspects** Participants seemed to be happy that the conference could be held under safe circumstances. Some even reported the feeling of a shared conference visit, despite it was being virtual, generated by alternative technical solutions. Overall, some participants still favour the meeting in person, because it allows a face-to-face interaction and deeper discussion. One person indicated that it was a pity that Gather was not used more often. The idea to allow exchanges in smaller groups of interest just after each presentation in Gather was raised.
- **Organisational aspects** In summary, the digital organisation was classified as great, because helpful instructions and information were given in advance, the time schedule was adhered to, the physical activity-breaks, and the well-structured programme. But at the same time, the need for more and longer breaks, because of sitting in front of the screen all day, was often mentioned. Because of that, attendees suggested to extend the breaks for the next conference.

## 5. Discussion

The purpose of this paper was, to give insight and recommendations to the different software and tools, that were used, about the materials, that were generated, for an online-conference, as a benefit for future conference organisers. Through the evaluation of the participants, it was possible to identify strengths and weaknesses, especially from the technical point of view. With these inputs and the open questions about possible improvements for the next online conference, different ideas for scaling-up could be extracted.

Without very detailed technical expertise, but with user manuals of the different systems, the support of the local IT-Service and volunteering colleagues, it was possible to organise a scientifically high appreciated and technical mostly fluent international online conference. Indico, with its functionalities, offers great features for building either an onsite- or online-conference. This conference management system was not evaluated through the survey. However, the organisers experienced it as a flexible system, with various features, which is easy to handle. We received only a few questions and problems from user suede. We strongly recommend it for planning future conferences.

The tools for realising the conference also mainly fulfilled their purpose. Only few technical issues with regard to using multiple platforms appeared. Which is an argument to use a single platform for next conferences. Knowing that some attendees did not check the video and handout instructions, before the conference started, we believe that if they did, a more success to use multiple platforms could have been approached. For instance, the instructions recommended, to keep both YouTube and Zoom continuously open in two browser tabs, for a smooth transition between the tools. Regarding the audio- and video-quality in Gather, which was mostly rated as good, only some participants reported connection problems. This may be due to the fact, that the Gather link, which expired after one day, was activated first. Therefore, a new link had to be forwarded for each conference day. Later, the functionality to activate one link for all conference days, was identified. We assume that the other connection problems are based upon a local problem with the internet, because in most of the cases, the presentations could be followed easily. For the case that the participants had problems, we offered them re-access to the pre-recorded presentations in Indico, because after each conference day, all presentations of that day were uploaded.

For simplifying the livestreaming, Zoom could be connected to OBS. YouTube Live can be omitted. Then, the switching back and forth between the two tabs, would disappear. This would serve as a flexible solution, which would also give presenters the opportunity to decide, if they would like to present live or use the pre-recording feature.

In addition, the focus should be moved in future online conferences to give more time for socialising, for instance via Gather. Despite the fact, that Gather was criticised by a few participants, it offered an appropriate platform to meet in smaller groups and discuss the poster and other presentations in a deeper manner. For a stable internet-connection, the conference space in Gather should be updated, because the tool can only be used free of charge by a dedicated number of participates at a time. But, if sufficient budget is available, more time should be given to Gather in future online conferences. These additional costs could be covered by sponsors filling advertising space in Gather. In the official conference programme, breaks should not be missed. These could be a mix of active physical activity breaks, which were mostly recommended, and normal breaks.

For the future, we expect increasing demand to combine the benefits of on-site and online formats, independent of pandemic situations. Different mixed formats with in person and virtual meetings could be tried. However, hybrid meetings with full individual flexibility for presenters and audience would need additional technical solutions.

## 6. Conclusion

Concluding, the following sentences summarize the lessons learned:

- Indico covers almost all functionalities for planning and conducting an online-conference.
- All tools for realising an online conference have to be explained to attendees in advance.
- Training materials should be shared with participants beforehand, in case of pre-recorded presentations.
- As backup, presentations should be recorded with presenter’s permission.
- A mix of active physical activity breaks and normal breaks should be considered.
- Enough time for breaks and for socialising in smaller groups should be part of the conference programme.

## List of abbreviations

EGREPA: European Group for Research into Elderly and Physical Activity
OBS: Open Broadcaster Software
POM: Photonics Online Meetup
WWU: University of Münster

---

**Additional file 1**

File name: Programme Procedures

Procedures during the conference sessions

**Additional file 2**

File name: Official Programme

Official conference programme including authors and titles of each presentation

**Additional file 3**

File name: Book of Abstracts

Digital book of abstracts with information to the keynote lecturers, organisational remarks, the final programme and abstracts of keynotes, symposia, oral presentations, poster presentations and physical activity projects for older adults

**Additional file 4**

File name: Checklist and practical tips for pre-recording

Instructions on how to prepare the pre-recording of the presentation

**Additional file 5**

File name: Checklist and practical tips for pre-recording

Instructions on how to prepare the pre-recording of the presentation

**Additional file 6**

File name: User-Guide: How to upload the presentation into Indico

A user-guide on how to technically upload a presentation into the conference-management-system Indico.

**Additional file 7**

File name: Gather instruction video and handout

A link to the video to explain Gather and a written text explaining the main functions of Gather

**Additional file 8**

File name: Answers to the open questions Q14, Q15 and Q16

Participants answers to the open question of the survey

## Declaration

### Ethics approval and consent to participate

Not applicable.

### Consent for publication

Not applicable.

### Availability of data and materials

All data generated or analysed during this study are included in this published article and its additional information files.

### Competing interests

The authors declare that they have no competing interests.

### Funding

This research received no external funding.

## Authors’ contributions

EB was mainly responsible for the organisation of the EGREPA 2021 conference, therefore she was responsible for leading the author team and prepare the original draft. EB was mainly responsible for writing the introduction and MB helped in reviewing the different tools. EB wrote the methods section. LR and LB were responsible for the results section. LR and LB were major contributors for the task of visualisation. EB was responsible for the discussion, LR and LB also made some improvements there. EB and MB were the major contributors for the conclusions. MB supervised and reviewed the manuscript. All authors have read and agree to the published version of the manuscript.

## Acknowledgements

We especially acknowledge the work carried out by Mona Ahmed, who revised the language of the manuscript. We also thank the IT-Service from the University of Münster WWU for helping us with its internally supported system Indico. For the YouTube Livestreaming with OBS, we thank Markus Jürgens for his great support. Without the great work from the Scientific Board and the EGREAPA board, it would not have been possible to plan and conduct this conference. Also, we thank all participants and speakers of the EGREPA 2021 conference.

## Authors information

EB works at the Institute of Sport and Exercise Sciences at the University of Münster as a research assistant and PhD Candidate. She acted as the responsible person for organising and conducting the EGREPA 2021 conference. LR and LB are students at the University of Münster and helped the organisers during the EGREPA 2021 conference with assistive work as part of an internship. MB is a professor at the Institute of Sport and Exercise Sciences at the University of Münster. For the EGREPA 2021 conference, he was a member of the scientific committee.

## Notes

### Competing Interest Statement

The authors have declared no competing interest.

