## Additional file 1 for "Planning and Conducting an Online Conference at the time of COVID-19: Lessons Learned from EGREPA 2021"

**PROGRAMME PROCEDURES**

As the event started, the moderator of the conference welcomed all participants in Zoom. Then the opening session started on YouTube. The opening session started with a tension rising GEMA-free 6-minutes soundtrack. For the duration of the soundtrack, a central slide with the title of the conference and the date was visible on YouTube. After that, several sequential pre-recorded welcome speeches from representatives of the different institutions and universities which jointly organised the conference were played. At the end of the opening session, a video with a traditional polish folk dance group was shown.

Thereafter, the first keynote speaker was introduced from a representative person of the conference in YouTube. Then, the first keynote lecture and the discussion took place via Zoom. After a short break, the first chair briefly introduced the symposium in Zoom and referred to YouTube, for attending the presentations. After each presentation, the discussion took place in YouTube.

Through various information slides on both platforms, conference attendees were constantly updated where the showplace was. For example, a slide with information that the opening session starts in a dedicated time in YouTube, was visible at the beginning in Zoom. After it, a slide with the information that the first keynote lecture starts soon in Zoom was visible on YouTube. For watching presentations, a reference to YouTube was provided. For the discussion, information to transfer back to Zoom was visible.

In the morning of the second and third conference day, participants were offered for a duration of 30 minutes to enter Gather. A reference to the training materials for getting to know Gather was again provided. Participants could try out all functionalities in the conference space, for example accessing posters and physical activity project movies for older adults, or talking to other participants.

Onwards, oral presentations to a specific topic were streamed via YouTube. Two representatives of the scientific committee led the discussion after these presentations in Zoom. Afterwards, the next symposium started. This process continued in such a way, it alternated between symposia and orals. Sometimes, within a symposium, a three-minute active break was visible on YouTube, which invited the participants to join exercising.

Before lunch-time of the second and third conference day, poster presentations with a duration of maximum three minutes were shown one after the other on YouTube. After it, it was possible to join small groups for poster discussion in Gather. After that, the conference proceeded with individual oral presentations and symposia.

The final keynote lecture was held live via Zoom. After it, Participants were invited to rate the conference through a survey, which could be accessed through a visible QR-Code in Zoom. In addition, a link to the survey was integrated into the chat. In the closing session, all participants were thanked for being part and attending the conference and the winner for the scientific research award was announced.
