## Additional file 2 for "Planning and Conducting an Online Conference at the time of COVID-19: Lessons Learned from EGREPA 2021"

| Day | Time (CET) | Programme |
| --- | --- | --- |
| Wednesday May, 19^th^ | 16:00-16:30 | **Opening** |
|  | 16:30-17:15 | **Keynote: Prof. Dr. Eling de Bruin**  **Design considerations for transforming video games into serious games**  Chair: Prof. Yael Netz |
|  | 17:15-17:30 | **Break** |
|  | 17:30-18:45 | **Symposium  Training of cognition and motor performance in very old age and nursing home residents: Feasibility and effects of tailored approaches** Chairs: Prof. Dr. Claudia Voelcker-Rehage & Ms. Madeleine Fricke   - **Uroš Marušič:** Cognitive approaches to enhance simple and complex locomotion in community-dwelling older adults - **Oliver Vogel:** Multimodal exercise effects in older adults depend on sleep, movement biography and habitual physical activity: A randomised controlled trial - **Thomas Cordes:** A multicomponent exercise intervention to improve motor functioning, cognition and psychosocial well-being for nursing home residents who are unable to walk - **Madeleine Fricke:** Feasibility of a multicomponent exercise intervention to enhance spatial navigation skills in nursing home residents |
| Thursday, May 20^th^ | 08:30-09:00 | **Gather is open**  Opportunity to view posters, videos and talk to other participants  Implemented accessible videos of physical activity projects for older adults:   - **Sylwia Talach-Kubas:** Using the Actimentia training platform to build active and healthy habits among elderly at a risk of dementia – good practice during pandemia - **Ismaël Brunot:** The Actimentia Motion Capture System - **Joanna Gradek:** Active and Healthy Senior, a project tailored to the times |
|  | 09:00-10:30 | **Oral Presentations**  **Physical Activity and Cognitive Functioning**  Chairs: Prof. Dr. Soledad Ballesteros & Dr. Oron Levin   - **Wouter Vints:** Resistance training and cognitive aging: Muscle-brain crosstalk - **Bohumila Krčmárová:** The influence of dance on the quality of life of older females - **Oron Levin:** Motor and cognitive inhibition in high and low fit old and young adults - **Soledad Ballesteros:** The effects of combined physical and cognitive training on cognition in healthy older adults: A systematic review and three-level meta-analysis - **Karolina Talar:** Augmenting the effects of aerobic training by transcranial direct current stimulation on cognition in older adults with and without cognitive impairment: A systematic review of randomised controlled trials |
|  | 10:30-10:45 | **Break** |
|  | 10:45-12:30 | **Symposium  Exercise and health of community-welling older adults (exercise before and during COVID-19 pandemic)** Chair: Prof. Dr. Vania Loureiro  Co-Chair: Dr. Anna Bukowska   - **Bebiana Sabino:** Postural balance as predictor of fall in community-dwelling older adults - **Miguel Peralta:** Grip strength and depressive symptoms among European middle-aged and older adults - **Tiago Rosa:** Effectiveness of a combined exercise program to improve functional fitness in community-dwelling older adults: A randomised controlled trial - **Miguel Peralta:** The association of grip strength with depressive symptoms among middle-aged older adults with different chronic diseases - **Adilson Marques:** Physical activity is negatively associated with depression symptoms independently of socioeconomic status - **Vânia Loureiro:** Home-based exercise program and health literacy intervention during COVID-19 pandemic: Up again senior pilot study |
|  | 12:30-13:00 | **Poster Session 1**  Chairs: Prof. Dr. Yael Netz & Prof. Dr. Wiebren Zijlstra   - **Ivan Serbetar:** The influence of fitness training program on resistance, flexibility and body fat mass of middle-aged women - **Gintarė Katkutė:** Neurochemical correlates of balance stability and dual-task effects in older adults with intact cognitive functioning and mild cognitive impaired patients - **Rodrigo Gallardo:** Sedentarism, mental health and presence of pathologies on Chilean older adults during confinement due to COVID-19 pandemic - **Gabrielle McKee:** Why older adults engage in a physical activity app: A qualitative analysis - **Tal Lifshits:** Physical activity level among adults – How does it change across life span and differ between men and women? - **Charlotte Sylvie Le Mouel:** Postural adjustments in anticipation of predictable perturbations allow elderly fallers to achieve a balance recovery performance equivalent to elderly non-fallers - **Kristīne Šneidere:** Relationship between cognitive reserve, physical activity, hippocampal volume and working memory in older adults - **Saar Frank Herschkovitz:** The effect of a single bout of aerobic exercise on limb motor inhibition as compared to cognitive inhibition in middle age adults - **Laimute Samsoniene:** The diversity of successful aging: The empowerment of sports veterans - **Kathleen Kang:** Plasticity of sequential decision-making in old age |
|  | 13:00-13:45 | **Gather**  **Individual Poster Discussions**  **Poster Session Room 1** |
|  | 13:45-15:00 | **Symposium**  **Relationship of physical and cognitive performance in community-dwelling older adults and nursing home residents**  Chairs: Prof. Dr. Bettina Wollesen & Dr. Katrin Müller   - **Stephanie Fröhlich:** Trajectories of cognitive performance in community-dwelling adults older than 80 years over a 16-month period - **Katrin Müller:** Relationship of physical activity and cognitive performance in community-dwelling older adults and nursing home residents - **Tina Auerswald:** Application of activity trackers among nursing home residents - Daily physical activity and sedentary behaviour and feasibility - **Katharina Zwingmann:** Comparison of physical and cognitive status of nursing home residents with and without the ability to walk |
|  | 15:00-15:15 | **Break** |
|  | 15:15-16:30 | **Oral Presentations**  **Physical Activity in Clinical Settings**  Chairs: Dr. Sylwia Mętel & Dr. Katarzyna Kucia   - **Tim Stuckenschneider:** Disease-inclusive – Time to rethink exercise classes?! - **Mona Ahmed:** Users' and other stakeholders' needs in the development of a personalised integrated care platform (PROCare4life) for older people with dementia or Parkinson disease - **Michael Brach:** Exercise groups for long-term cardiac rehabilitation in Germany: Feasability and acceptance survey three month after changes in medical care - **Sylwia Mętel:** Chest mobility of older adults participating in speleotherapy combined with pulmonary rehabilitation |
|  | 16:30-16:45 | **Break** |
|  | 16:45-17:45 | **Symposium**  **ADYMA: Psycho-social and behavioural impact of an adapted physical activity program for seniors living in residential-based communities**  Chair: Prof. Dr. Sebastien Chastin  Co-Chair: Prof. Dr. Michael Brach   - **Sebastien Chastin:** Interventions to reduce sedentary behavior in older adults: A Cochrane systematic review of the literature - **Amélie Baghdiguian & Yannick Manet:** ADYMA: A program to encourage mobility and promote healthy aging among seniors living in residences - **Ariane Gautier & Gonzalo Marchant:** Embedding psycho-social approach in a program designed to promote mobility in seniors living in residence |
| Friday, May 21^th^ | 08:30-09:00 | **Gather is open**  Opportunity to view posters and videos and talk to other participants (see page 1) |
|  | 09:00-10:45 | **Symposium**  **Healthy aging: From the brain to the muscles**  Chair: Prof. Dr. Anita Hökelmann   - **Anita Hökelmann & Achraf Ammar:** The effects of physical activity - dance and sport intervention- on brain structure and cognition in elderly - **Rado Pišot:** Early neuromuscular alterations in older compared to younger men following prolonged bed rest and subsequent recovery - **Boštjan Šimunič:** Age-related slowing of skeletal muscles in non-athletes, power and endurance master athletes - **Uroš Marušič:** The role of enhanced cognition for mobility improvements: A neurophysiological perspective - **Bernhard Grässler:** Neurophysiological correlates of cognitive performance in elderly: Heart rate variability as part of multimodal measurement approach - **Nicole Halfpaap:** Neuromuscular functions of MCI patients in comparison to healthy seniors - muscle contractile properties measured with Tensiomyography |
|  | 10:45-11:00 | **Break** |
|  | 11:00-12:30 | **Oral Presentations**  **Motor Performance and Functional Fitness**  Chairs: Dr. Magdalena Majer & Prof. Dr. Yael Netz   - **Kaisa Koivunen:** Birth cohort differences in maximal physical performance in 75- and 80-year-old men and women: A comparison of two cohorts over 28 years - **Magdalena Majer:** Changes in functional fitness and somatic indicators of women participating in the “Active and Healthy Senior” project - **Nikola Stračárová:** Influence of physical activity on reaction rate in a selected group of seniors - **Agnieszka Kowalska:** Evaluation of the impact of an exercise program implemented in an outdoor gym on the physical fitness people over 60 years old - **Yael Netz:** Prescribing individualised exercise programs based upon remote assessments of motor fitness: A pilot study among healthy people aged 65 and over |
|  | 12:30-13:00 | **Poster Session 2**  Chair: PD Dr. med Timo Hinrichs   - **Małgorzata Bagińska:** Nutritional behaviour and selected lifestyle aspects in patients of a physiotherapy clinic in Ustrzyki Dolne, Poland, in the context of osteoporosis prevention - **Zbigniew Ossowski:** Association between sarcopenia related parameters and cognitive functions in postmenopausal women - **Jessica Koschate:** Covid-19 pandemic – Time course of physical activity and functional abilities in active older people associated with the lockdown periods in Germany - **Katarzyna Kucia:** The effects of 3-month aqua-fitness program on the functional fitness in elderly women - **Aileen Lynch:** A pilot study to assess the effectiveness of a brief intervention on community-dwelling older adults’ physical activity - **Gina Krause:** Physical Activity and exercising benefits in dementia care - **Ana Maria Rizescu:** Means of injury prevention and the influence of the correlation between the type of temperament and the severity of injuries in the game of football-tennis      - **Daniele Magistro:** Effectiveness of a lifestyle intervention in promoting the mobility functioning in institutionalised older adults - **Monika Wilk-Głodzik:** Perception of physical self-attractiveness in terms of pro-health behaviour including physical activity and the use of beauty treatments in women over 60 |
|  | 13:00-13:45 | **Gather**  **Individual Poster Discussions**  **Poster Session Room 2** |
|  | 13:45-15:00 | **Symposium**  **Life-Space mobility in old age**  Chairs: Dr. Eleftheria Giannouli, PD Dr. med. Timo Hinrichs   - **Eleftheria Giannouli:** Life-space mobility in patients with neurological diseases: A scoping review - **Julia Seinsche:** Mobility in frail older adults: First insights into its relationship with physical activity and life-space mobility - **Carl-Philipp Jansen:** Intervention effects on life-space mobility: Current evidence and future perspectives - **Erja Portegijs:** Older adults´ activities in the life-space before and during COVID-19 restrictions |
|  | 15:00-15:15 | **Break** |
|  | 15:15-16:45 | **Oral Presentations**  **Movement, Activities and Lifestyles**  Chairs**:** Dr. Rafał Stemplewski & Prof. Dr. Michael Brach   - **Veronique Wolter:** Care providers and sports clubs? Chances and barriers of joint activities for older people in stationary and ambulant settings - **Manca Peskar:** Age-related slowing of hand and food reaction times - **Lenka Svobodova:** Variables interfering into the quality of standing balance in older adults - **Stephanie Schmidle:** Insights from monitoring activities of daily living in elderlies with a smartwatch - **Ellen Bentlage:** Practical recommendations for maintaining active lifestyle during the COVID-19 pandemic: A systematic literature review |
|  | 16:45-17:00 | **Break** |
|  | 17:00-17:45 | **Keynote: Prof. Dr. Taina Rantanen**  **Is old age changing? Views on muscle strength, mobility, activity and survival**  Chair: PD Dr. med. Timo Hinrichs |
|  | 17:45-18:15 | **Closing** |
