## Additional file 3 for "Planning and Conducting an Online Conference at the time of COVID-19: Lessons Learned from EGREPA 2021"

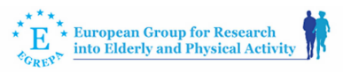

**EGREPA Conference 2021**

***“Active aging - new challenges and new opportunities”***

BOOK OF ABSTRACTS

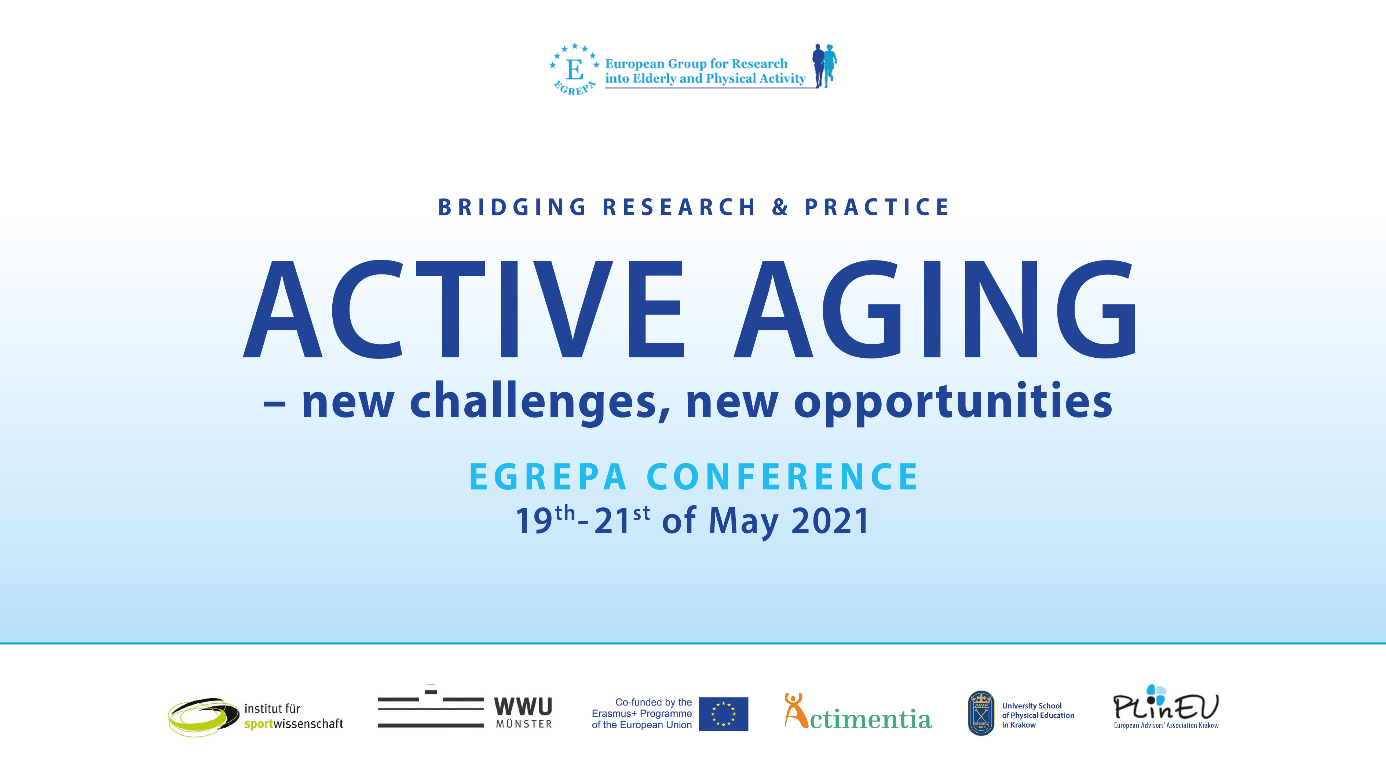

**The Conference is jointly organised by:**

European Group for Research into Elderly and Physical Activity (EGREPA)
University of Münster WWU
University of Physical Education in Krakow
European Advisors’ Association PlinEU.

This conference introduces results from the Erasmus+ project „Actimentia – Physical activity and exercising benefits in dementia care“ and is co-funded by the European Commission.

**Organising Committee:**

- Ellen Bentlage – *University of Münster, Germany*
- Dr. Joanna Gradek – *University of Physical Education in Krakow, Poland*
- Sylwia Tałach-Kubas – *European Advisors’ Association PLinEU, Poland*

**Scientific Committee:**

Head: Prof. Dr. Yael Netz – *Academic College at Wingate, Israel*

Members:

- Prof. Dr. Michael Brach – *University of Münster, Germany*
- Prof. Dr. Heinz Mechling, Emeritus – *German Sport University Cologne, Germany*
- PD. Dr. Timo Hinrichs – *University of Basel, Switzerland*
- Prof. Dr. Rafał Stemplewski –*University School of Physical Education, Poznan, Poland*
- Prof. Dr. Oron Levin – *Katholic University Leuven, Belgium*
- Prof. Dr. Wiebren Zijlstra – *German Sport University Cologne, Germany*
- Prof. Dr. Soledad Ballesteros – *Universidad Nacional de Educación a Distancia (UNED), Madrid, Spain*
- Prof. Dr. Anita Hökelmann – *Otto-von-Guericke-University Magdeburg, Germany*
- Dr. Magdalena Majer – *University of Physical Education in Krakow, Poland*
- Dr. Anna Bukowska – *University of Physical Education in Krakow, Poland*
- Dr. Sylwia Mętel – *University of Physical Education in Krakow, Poland*

© University of Münster, 2021

Editors: Ellen Bentlage, Michael Brach, Yael Netz, Timo Hinrichs, Heinz Mechling & Sylwia Tałach-Kubas

ISBN: 978-3-00-068824-9

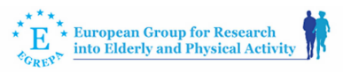

The European Group for Research into Elderly and Physical Activity aims to promote physical activity and health in older adults through the carrying out and promotion of research and the collection and diffusion of information related to this field of interest.
[EGREPA Website](http://www.egrepa.org)

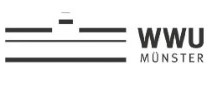

The University of Münster (WWU) is one of Germany's largest institutions of higher education (~46,000 students, 600 professors and 4500 research staff). The Institute of Sport and Exercise Sciences includes six departments from natural science, social science and humanities. Research and teaching addresses target groups from preschool children to the oldest old, and includes physical education as well as high performance sport, preventive exercise as well as everyday activity. [University of Münster Website](https://www.uni-muenster.de/en/) + [Movement Science Website](https://www.uni-muenster.de/Sportwissenschaft/en/Bewegungswissenschaft/index.html)

[WWU Münster - Website](https://www.uni-muenster.de/en/)

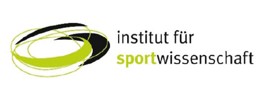

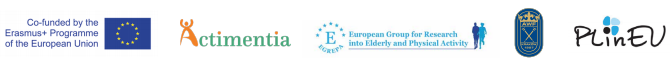

Started more than 30 yeas ago, Erasmus+ is the EU's programme to support education, training, youth and sport. Beneath mobility, strategic partnerships is a major strand. In addition to offering grants, Erasmus+ also supports teaching, research, networking and policy debate on EU topics. The sports chapter promotes grassroots activities in sports.

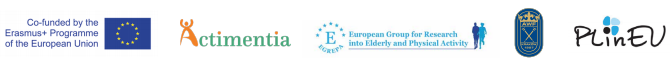

Actimentia is an Erasmus+ project whose purpose is to foster physical activity in the everyday life of people with mild cognitive impairment or dementia and their caregivers. The developed on-line platform offers a great choice on exercises that can be used as a tool for preserving their quality of life and wellbeing of all 60+ persons. <https://actimentia.org/>. Project Reference: 2018-1-DE02-KA204-005231

The Bronisław Czech University of Physical Education in Cracow is one of the best physical education universities in Poland. Education takes place within three faculties: the Faculty of Physical Education and Sports, where, among others, studies in the field of Physical Culture for the Elderly are offered, the Faculty of Movement Rehabilitation, the Faculty of Tourism and Recreation. Almost 4000 young people study at the AWF. Teaching and research activities are carried out by a staff of approximately 270 academic teachers. Many graduates of the University of Physical Education are outstanding athletes, such as Kamil Stoch and Agnieszka Radwańska. University website: <https://www.awf.krakow.pl/>

European Advisors’ Association PLinEU is a non-governmental organization from Krakow (Poland), which elaborates and implements innovative methods of lifelong learning, among others - elderly adults education projects. We are partner of ACTIMENTIA project and we have tested with great success training programme on 2 groups of 60+ persons at risk of dementia or suffering from MCI.

<https://www.plineu.org/en/>

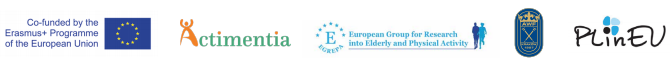

**Welcome Greetings:**

The European Group for Research into Elderly and Physical Activity (EGREPA) is delighted to welcome all of you to this on-line Conference on: Active Aging – New Challenges, New Opportunities – Bridging Research & Practice. Our scientific program includes two scholars who will deliver a central keynote speech, as well as symposiums, oral sessions and posters. We wish you all a pleasant, fruitful conference and hope to meet you on-site in future EGREPA conferences.

**Keynote Lecturers:**

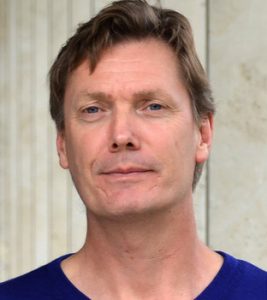
**Prof. Dr. Eling de Bruin** is a full professor in Karolinska Institute, Sweden & ETH Zurich, Switzerland. His research focuses on the relation between the use of exergames and their influence on physical & cognitive functioning of elderly. His research specifically addresses the theoretical relevance of novel Exergame training approaches and helps establishing research data that underpin the relevance of novel diagnostic systems and game-based brain exercise training that specifically focus on aspects of neuromuscular functioning in (frail) elderly aimed at ameliorating cognitive & physical function. Prof. de Bruin also focuses on the intimately linked motor and cognitive aspects of human movement and age-related risk of falling. He extends his findings ‘from bench to bedside’ using efficient combinations of basic and advanced technologies to capture the plasticity of central and peripheral motor control systems as well as of cognition in response to physical activity and training interventions. Prof. de Bruin is the author of over 280 publications in top journals. His skills and expertise include a long list of fields. He is a member of a number of Journals' Editorial Boards, the most important of which is the EURAPA – European Review of Aging and Physical Activity – the EGREPA Journal.

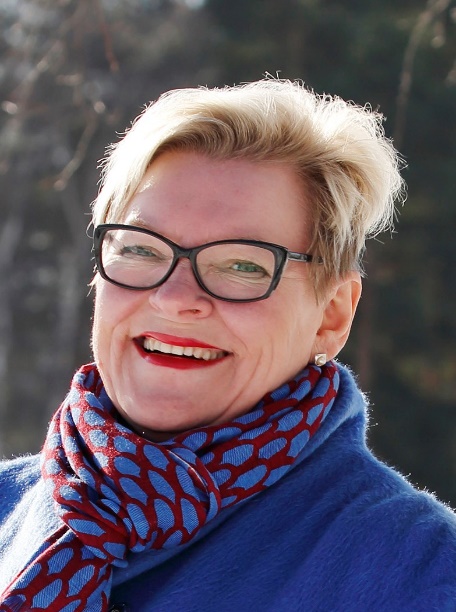
**Prof. Dr. Taina Rantanen**, Professor of Gerontology and Public Health at the Faculty of Sport and Health Sciences, University of Jyväskylä, Finland is one of the world leaders in research on old age, functioning and active ageing. She has led numerous studies funded by national and international science organizations; in 2016 she received a highly competitive ERC Advanced Grant for her study ‘Active Ageing – Resilience and External Support as Modifiers of the Disablement Process’ (AGNES). Prof. Rantanen has participated in national (Academy of Finland, Ministry of Health and Social Affairs, Ministry of Education and Culture) and European (Joint Programming Initiative) strategy and evaluation panels and has been employed by the U.S. National Institutes of Health and the World Health Organization. Prof. Rantanen is a member of editorial boards of ‘Journals of Gerontology: Medical Sciences’, ‘Journal of Aging and Frailty’, ‘Journal of Aging and Health’ and ‘European Review of Aging and Physical Activity (EURAPA)’. She has received several awards for scientific achievements as well as for promoting public and policy awareness of ageing issues. Honors include the prestigious Sohlberg Nordic Gerontology Prize awarded by the Nordic Gerontology Federation, the Schildt Prize awarded by the Finnish Cultural Foundation and the Knight (1st class) of the Order of the White Rose awarded by the Finnish Government. Besides authoring numerous top ranked and highly cited international journal publications, she is also editor of the Gerontology textbook ‘Gerontologia’ (in Finnish) and author of several textbook chapters (nationally and internationally). Prof. Rantanen’s Keynote Lecture at the EGREPA Conference 2021 is entitled ‘Is old age changing? Views on muscle strength, mobility, activity and survival’.

**Organisational remarks:**

On the next pages (6-11) you can access the detailed programme of our conference. The left column displays the conference day, beside it you find the time of the respective session and in the right column the programme items are listed. In the conference programme, only the presenters with title of their presentation are listed. Information about the affiliations, authors and abstracts can be retrieved starting from page 18. All oral presentations are assigned to a special topic. Posters are divided into two sessions. All pre-recorded presentations will be streamed live via our YouTube-Stream, in order to ensure the best technical conditions. Zoom will be used for the live presentations of the keynote lecturers and for short discussions after each livestreamed presentation, with the exception of poster discussions. These will take place via the 2D space Gather, where participants of the conference can walk virtually with an avatar though our conference lobby, poster session rooms and a lounge with a warm atmosphere. All videos of the presentations will be additionally uploaded to our conference system Indico. Participants of the conference will have free access to all pre-recorded presentations from 19th until 27th of May, 2021.

**Final Programme:**

| Day | Time (CET) | Programme |
| --- | --- | --- |
| Wednesday May, 19^th^ | 16:00-16:30 | **Opening** |
|  | 16:30-17:15 | **Keynote: Prof. Dr. Eling de Bruin**  **Design considerations for transforming video games into serious games**  Chair: Prof. Yael Netz |
|  | 17:15-17:30 | **Break** |
|  | 17:30-18:45 | **Symposium  Training of cognition and motor performance in very old age and nursing home residents: Feasibility and effects of tailored approaches** Chairs: Prof. Dr. Claudia Voelcker-Rehage & Ms. Madeleine Fricke   - **Uroš Marušič:** Cognitive approaches to enhance simple and complex locomotion in community-dwelling older adults - **Oliver Vogel****:** Multimodal exercise effects in older adults depend on sleep, movement biography and habitual physical activity: A randomised controlled trial - **Thomas Cordes****:** A multicomponent exercise intervention to improve motor functioning, cognition and psychosocial well-being for nursing home residents who are unable to walk - **Madeleine Fricke:** Feasibility of a multicomponent exercise intervention to enhance spatial navigation skills in nursing home residents |
| Thursday, May 20^th^ | 08:30-09:00 | **Gather is open**  Opportunity to view posters, videos and talk to other participants  Implemented accessible videos of physical activity projects for older adults:   - **Sylwia Ta**ł**ach-Kubas:** Using the Actimentia training platform to build active and healthy habits among elderly at a risk of dementia – good practice during pandemia - **Ismaël Brunot:** The Actimentia Motion Capture System - **Joanna Gradek:** Active and Healthy Senior, a project tailored to the times |
|  | 09:00-10:30 | **Oral Presentations**  **Physical Activity and Cognitive Functioning**  Chairs: Prof. Dr. Soledad Ballesteros & Dr. Oron Levin   - **Wouter Vints:** Resistance training and cognitive aging: Muscle-brain crosstalk - **Bohumila Krčmárová:** The influence of dance on the quality of life of older females - **Oron Levin:** Motor and cognitive inhibition in high and low fit old and young adults - **Soledad Ballesteros:** The effects of combined physical and cognitive training on cognition in healthy older adults: A systematic review and three-level meta-analysis - **Karolina Talar:** Augmenting the effects of aerobic training by transcranial direct current stimulation on cognition in older adults with and without cognitive impairment: A systematic review of randomised controlled trials |
|  | 10:30-10:45 | **Break** |
|  | 10:45-12:30 | **Symposium  Exercise and health of community-welling older adults (exercise before and during COVID-19 pandemic)** Chair: Prof. Dr. Vânia Loureiro  Co-Chair: Dr. Anna Bukowska   - **Nuno Loureiro:** Effectiveness of a combined exercise program to improve functional fitness in community-dwelling older adults: A randomised controlled trial - **Miguel Peralta:** Grip strength and depressive symptoms among European middle-aged and older adults - **Bebiana Sabino:** Postural balance as predictor of fall in community-dwelling older adults - **Priscilla Marconcin:** The association of grip strength with depressive symptoms among middle-aged older adults with different chronic diseases - **Adilson Marques:** Physical activity is negatively associated with depression symptoms independently of socioeconomic status - **Vânia Loureiro:** Home-based exercise program and health literacy intervention during COVID-19 pandemic: Up again senior pilot study |
|  | 12:30-13:00 | **Poster Session 1**  Chairs: Prof. Dr. Yael Netz & Prof. Dr. Wiebren Zijlstra   - **Ivan Serbetar:** The influence of fitness training program on resistance, flexibility and body fat mass of middle-aged women - **Gintarė Katkutė:** Neurochemical correlates of balance stability and dual-task effects in older adults with intact cognitive functioning and mild cognitive impaired patients - **Rodrigo Gallardo:** Sedentarism, mental health and presence of pathologies on Chilean older adults during confinement due to COVID-19 pandemic - **Gabrielle McKee:** Why older adults engage in a physical activity app: A qualitative analysis - **Tal Lifshits:** Physical activity level among adults – How does it change across life span and differ between men and women? - **Kristīne Šneidere:** Relationship between cognitive reserve, physical activity, hippocampal volume and working memory in older adults - **Saar Frank Herschkovitz:** The effect of a single bout of aerobic exercise on limb motor inhibition as compared to cognitive inhibition in middle age adults - **Laimute Samsoniene:** The diversity of successful aging: The empowerment of sports veterans - **Kathleen Kang:** Plasticity of sequential decision-making in old age |
|  | 13:00-13:45 | **Gather**  **Individual Poster Discussions**  **Poster Session Room 1** |
|  | 13:45-15:00 | **Symposium**  **Relationship of physical and cognitive performance in community-dwelling older adults and nursing home residents**  Chairs: Prof. Dr. Bettina Wollesen & Dr. Katrin Müller   - **Katrin Müller:** Relationship of physical activity and cognitive performance in community-dwelling older adults and nursing home residents - **Stephanie Fröhlich:** Trajectories of cognitive performance in community-dwelling adults older than 80 years over a 16-month period - **Tina Auerswald:** Application of activity trackers among nursing home residents - Daily physical activity and sedentary behaviour and feasibility - **Katharina Zwingmann:** Comparison of physical and cognitive status of nursing home residents with and without the ability to walk |
|  | 15:00-15:15 | **Break** |
|  | 15:15-16:30 | **Oral Presentations**  **Physical Activity in Clinical Settings**  Chairs: Dr. Sylwia Mętel & Dr. Katarzyna Kucia   - **Tim Stuckenschneider:** Disease-inclusive – Time to rethink exercise classes?! - **Mona Ahmed:** Users' and other stakeholders' needs in the development of a personalised integrated care platform (PROCare4life) for older people with dementia or Parkinson disease - **Michael Brach:** Exercise groups for long-term cardiac rehabilitation in Germany: Feasability and acceptance survey three month after changes in medical care - **Sylwia Mętel:** Chest mobility of older adults participating in speleotherapy combined with pulmonary rehabilitation |
|  | 16:30-16:45 | **Break** |
|  | 16:45-17:45 | **Symposium**  **ADYMA: Psycho-social and behavioural impact of an adapted physical activity program for seniors living in residential-based communities**  Chair: Prof. Dr. Sebastien Chastin  Co-Chair: Prof. Dr. Michael Brach   - **Sebastien Chastin:** Interventions to reduce sedentary behavior in older adults: A Cochrane systematic review of the literature - **Amélie Baghdiguian & Yannick Manet:** ADYMA: A program to encourage mobility and promote healthy aging among seniors living in residences - **Ariane Gautier & Gonzalo Marchant:** Embedding psycho-social approach in a program designed to promote mobility in seniors living in residence |
| Friday, May 21^th^ | 08:30-09:00 | **Gather is open**  Opportunity to view posters and videos and talk to other participants (see page 6) |
|  | 09:00-10:45 | **Symposium**  **Healthy aging: From the brain to the muscles**  Chair: Prof. Dr. Anita Hökelmann   - **Anita Hökelmann & Achraf Ammar:** The effects of physical activity - dance and sport intervention- on brain structure and cognition in elderly - **Rado Pišot:** Early neuromuscular alterations in older compared to younger men following prolonged bed rest and subsequent recovery - **Boštjan Šimunič:** Age-related slowing of skeletal muscles in non-athletes, power and endurance master athletes - **Uroš Marušič:** The role of enhanced cognition for mobility improvements: A neurophysiological perspective - **Bernhard Grässler:** Neurophysiological correlates of cognitive performance in elderly: Heart rate variability as part of multimodal measurement approach - **Nicole Halfpaap:** Neuromuscular functions of MCI patients in comparison to healthy seniors - muscle contractile properties measured with Tensiomyography |
|  | 10:45-11:00 | **Break** |
|  | 11:00-12:30 | **Oral Presentations**  **Motor Performance and Functional Fitness**  Chairs: Dr. Magdalena Majer & Prof. Dr. Yael Netz   - **Kaisa Koivunen:** Birth cohort differences in maximal physical performance in 75- and 80-year-old men and women: A comparison of two cohorts over 28 years - **Magdalena Majer:** Changes in functional fitness and somatic indicators of women participating in the “Active and Healthy Senior” project - **Nikola Stračárová:** Influence of physical activity on reaction rate in a selected group of seniors - **Agnieszka Kowalska:** Evaluation of the impact of an exercise program implemented in an outdoor gym on the physical fitness people over 60 years old - **Yael Netz:** Prescribing individualised exercise programs based upon remote assessments of motor fitness: A pilot study among healthy people aged 65 and over |
|  | 12:30-13:00 | **Poster Session 2**  Chair: PD Dr. Timo Hinrichs   - **Małgorzata Bagińska:** Nutritional behaviour and selected lifestyle aspects in patients of a physiotherapy clinic in Ustrzyki Dolne, Poland, in the context of osteoporosis prevention - **Zbigniew Ossowski:** Association between sarcopenia related parameters and cognitive functions in postmenopausal women - **Jessica Koschate:** Covid-19 pandemic – Time course of physical activity and functional abilities in active older people associated with the lockdown periods in Germany - **Katarzyna Kucia:** The effects of 3-month aqua-fitness program on the functional fitness in elderly women - **Aileen Lynch:** A pilot study to assess the effectiveness of a brief intervention on community-dwelling older adults’ physical activity - **Gina Krause:** Physical Activity and exercising benefits in dementia care - **Ana Maria Rizescu:** Means of injury prevention and the influence of the correlation between the type of temperament and the severity of injuries in the game of football-tennis      - **Daniele Magistro:** Effectiveness of a lifestyle intervention in promoting the mobility functioning in institutionalised older adults - **Paulina Łukaszek:** Perception of physical self-attractiveness in terms of pro-health behaviour including physical activity and the use of beauty treatments in women over 60 - **Charlotte Sylvie Le Mouel:** Postural adjustments in anticipation of predictable perturbations allow elderly fallers to achieve a balance recovery performance equivalent to elderly non-fallers |
|  | 13:00-13:45 | **Gather**  **Individual Poster Discussions**  **Poster Session Room 2** |
|  | 13:45-15:00 | **Symposium**  **Life-Space mobility in old age**  Chairs: Dr. Eleftheria Giannouli, PD Dr. Timo Hinrichs   - **Eleftheria Giannouli:** Life-space mobility in patients with neurological diseases: A scoping review - **Julia Seinsche:** Motility in frail older adults: First insights into its relationship with physical activity and life-space mobility - **Carl-Philipp Jansen:** Intervention effects on life-space mobility: Current evidence and future perspectives - **Erja Portegijs:** Older adults´ activities in the life-space before and during COVID-19 restrictions |
|  | 15:00-15:15 | **Break** |
|  | 15:15-16:45 | **Oral Presentations**  **Movement, Activities and Lifestyles**  Chairs**:** Dr. Rafał Stemplewski & Prof. Dr. Michael Brach   - **Veronique Wolter:** Care providers and sports clubs? Chances and barriers of joint activities for older people in stationary and ambulant settings - **Manca Peskar:** Age-related slowing of hand and food reaction times - **Lenka Svobodová:** Variables interfering into the quality of standing balance in older adults - **Stephanie Schmidle:** Insights from monitoring activities of daily living in elderlies with a smartwatch - **Ellen Bentlage:** Practical recommendations for maintaining active lifestyle during the COVID-19 pandemic: A systematic literature review |
|  | 16:45-17:00 | **Break** |
|  | 17:00-17:45 | **Keynote: Prof. Dr. Taina Rantanen**  **Is old age changing? Views on muscle strength, mobility, activity and survival**  Chair: PD Dr. Timo Hinrichs |
|  | 17:45-18:15 | **Closing** |

### KEYNOTE LECTURES

Prof. Dr. Eling De Bruin,

*ETH Zürich, Switzerland and Karolinska Institute, Sweden*

###### Design considerations for transforming video games into serious games

An increase of muscle strength and improved balance is achievable with traditional training and this, hence, positively influences some measures of gait. However, these exercise components often do not impact on spatial and temporal characteristics of gait that are associated with distinct brain networks. Because these gait characteristics are associated with distinct brain networks, it can be hypothesised that addressing neuronal loss in these networks may be an important strategy to prevent mobility disability in older adults. A way to bring in a cognitive element into an exercise program is the use of virtual reality techniques. There are reports on the use and effects of virtual reality exergaming-training in various populations. Methods using immersive computer technologies resulted in improved motor functions of upper extremities and a cortical activation after virtual reality intervention in patients with chronic stroke. Older adults benefited from training in terms of improved functional abilities, postural control and simple auditory reaction times under dual task conditions. This talk will focus on the relation between the use of Exergames and their influence on physical functioning of elderly. As people age, a self-reinforcing, downwards spiral of reduced interaction with challenging environments and reduced brain health significantly contribute to cognitive decline. Furthermore, brain activity needs to be able to adapt to challenges posed from the environment. Novel training paradigms; e.g. virtual reality interaction exergaming, indicate they might be able to effect on brain functioning in elderly. This talk specifically discusses [1] the theoretical relevance of novel Exergame training approaches and [2] presents research data suggesting that game-based brain exercise training with a focus on aspects of neuromuscular functioning in (frail) elderly are effective for ameliorating cognitive & physical gait function.

Prof. Dr. Taina Rantanen,
*University of Jyväskylä, Finland*

###### Is old age changing? Views on muscle strength, mobility, activity and survival

Research on active ageing has been hindered, because the well-known WHO policy definition of active ageing is not feasible for surveys, but rather can be used to rate countries. To remedy this, we created an operational definition applicable to individuals: a person’s striving for activity as per one’s goals, capacities and available opportunities, and developed the University of Jyväskylä active ageing scale (UJACAS) that includes 17 items each covering goals, capacities, opportunities and the actual activity. The score is the sum total of all responses (range 0-272), with higher scores indicating a more active approach to life. Mobility and psychological resources underlie active ageing scores, which are highest among those with intact mobility, somewhat lower among those with slightly limited mobility and lowest among those with more severe mobility limitation. Better psychological coping correlates positively with activity among those with intact or slightly limited mobility but not among those with more severe mobility limitation. This relates to an interesting new approach to positive ageing, i.e. resilience. Resilience refers to better than expected tolerance of or recovery from adversity, while robustness refers to a low risk of adversity. For example, higher muscle strength predicts lower risk of mortality and disability, i.e. indicates robustness. We recently reported that pre-fracture muscle strength predicts better post-fracture survival, and that its predictive power is higher during post-fracture time than during non-fracture time, which indicates resilience. COVID-19 pandemic provided an opportunity to study social distancing as an adversity. An important predictor of resilience of quality of life was walking speed assessed before the pandemic, while psychological coping and not being lonely predicted robustness of quality of life. In all, positive changes have taken place in ageing. In the current 75 to 85-year-old cohorts, the cognitive and physical functioning is 5-10 years ‘younger’ than among those of the same age were 30 years ago, suggesting that old age has moved to older ages and midlife has grown longer. The longer and healthier lives underline the importance of raising awareness about active ageing and the potential of older people to contribute to the society.

### SYMPOSIA PRESENTATIONS

###### Training of cognition and motor performance in very old age and nursing home residents: Feasibility and effects of tailored approaches

Symposium Chair:
Prof. Dr. Claudia Voelcker-Rehage, *University of Muenster, Department Neuromotor Behavior and Exercise, Germany* & Ms. Madeleine Fricke, *Berlin University of Technology, Germany*

For a large number of older people, an independent lifestyle is an essential ability, which affects the quality of life. This is opposed by aging-associated declining cognitive abilities and decreasing mobility (Seidler et al., 2010). Especially the simultaneous execution of cognitive and motor processes prerequisites the efficient performance of most daily activities (Lacour et al., 2008). Since research has repeatedly shown that brain and body remain plastic in old age (Erickson & Kramer, 2009; Park & Bischof, 2013), numerous studies examined different training approaches to slow down or even reverse the decline of cognitive and motor function. This symposium introduces studies that both evaluate the impact of cognitive and multi-component exercise interventions in community-dwelling older adults and nursing home residents. The first two talks will address the main outcomes (simple and complex locomotion) and covariates (e.g. fitness, emotion and cognition) of training approaches in community-dwelling older adults. Talk three and four will concern nursing home residents. First, the effects of a multicomponent exercise in non-walkers will be presented, focusing on the improvement of motor functioning, cognition, and psychosocial wellbeing. The last talk will demonstrate the development and modification of a multicomponent exercise intervention, which aims to improve the spatial orientation ability and life-space mobility of walkable nursing home residents. The concluding discussion will analyse how systematic piloting of training approaches improve the feasibility for the respective target group and which effects can be achieved by different training approaches for older persons, irrespective of whether they live independently or in a nursing home.

###### Uroš Marušič: Cognitive approaches to enhance simple complex locomotion in community-dwelling older adults

Authors: Uroš Marušič^1,3^, Jeannette Mahoney^2^, Joe Verghese^2^

^1^ Science and Research Center Koper, Slovenia

^2^ Albert Einstein College of Medicine, USA

^3^ Alma Mater Europaea – ECM, Slovenia

A close inter-relationship between mobility and cognition is reported in older adults, with improvements in gait performance noticeable after cognitive remediation in frail individuals. The aim of this study was to evaluate the effect of computerized cognitive training (CCT) in healthy and independently living older adults, and determine whether CCT was associated with neuroplasticity. Using a randomized single-blind control design, sixty-three adults age (age >60 years; mean Montreal Cognitive Assessment; MoCA score = 27) were randomly assigned to either a 2-month CCT (8 weeks, 3x/week, 40 min/session) or wait-list. Primary outcomes included gait speed during normal and dual-task walking conditions. Secondary outcomes included executive control and sensorimotor integration processes assessed via electroencephalography/event-related potentials (EEG/ERP). Results from a linear mixed effect model, adjusted for baseline MoCA score, age and gender revealed that CCT improved executive functions (p < 0.05), as well as dual-task walking speed (p < 0.05). These improvements were accompanied with shorter behavioural responses (p < 0.05), as evidenced by shorter P2 latencies on ERP (occipital electrodes, p < 0.05) and enhanced motoric processes (central electrodes, p < 0.05). CCT improved mobility in active independently living older adults. Electrophysiological outcomes support possible neuroplasticity effects that might explain the mobility gains of CCT.

###### Oliver Vogel: Multimodal exercise effects in older adults depend on sleep, movement biography and habitual physical activity: A randomised controlled trial

Authors: Oliver Vogel^1^, Daniel Niederer^2^, Lutz Vogt^2^

^1^ University of Hamburg, Germany

^2^ Goethe University Frankfurt, Germany

Background: Promotion of healthy ageing is one of the major challenges for health care systems in current times. The present study investigates the effects of a standardised physical activity intervention for older adults with regard to their movement biography, sleep parameters and current activity behavior. Methods: This single-blinded, randomised controlled trial included 49 older adults (36 female; 82.9 ± 4.5 (M ± SD)). Movement biography, sleep parameters, cognitive capacity, subjective health status and fear of falls were assessed by means of questionnaires. Functional status, gait and current activity levels were captured using accelerometers, a pressure plate and physiological tests. The multicomponent intervention took place 45 minutes twice a week for 16 weeks. Intervention effects were evaluated in univariate anovas between subcohorts of different sleep duration in both groups, investigating changes in primary objectives while using the respective baseline values, movement biography and current activity levels as covariates. Results: We found differences in cognitive capacity change scores according to group allocation (F = 4.483; p = .008). Effects of group allocation on fear of falls (F = 12.260; p = .001) and balance change scores (F = 7.697; p = .008) were modified by current activity. Effects of group allocation on gait cadence were modified by the lifetime physical activity level (F = 4.124; p = .049). Conclusion: Multicomponent interventions as well as adequate sleep durations appear to provide combinable beneficial effects for cognitive capacity. Gait, fear of falls and flexibility seem to be affected more by movement biography and current physical activity levels than participation in generalised interventions.

###### Thomas Cordes: A multicomponent exercise intervention to improve motor functioning, cognition and psychosocial well-being for nursing home residents who are unable to walk

Authors: Thomas Cordes^1^, Julian Rudisch^2^, Katharina Zwingmann^3^, Claudia Voelcker-Rehage^2^, Bettina Wollesen^1^

^1^ University of Hamburg, Germany, Department of Human Movement and Training Science

^2^ University of Münster, Department Neuromotor Behavior and Exercise, Germany

^3^ Chemnitz University of Technology, Department of Sport Science, Germany

Nursing home residents are often characterized by multimorbidity and dependency. Tailored multicomponent exercise programs have been shown to induce positive effects on mobility, cognitive and psychosocial resources. Most exercise studies, however, aimed at residents who are able to walk and did not consider those unable to walk. The aim of this study was to determine the effects of a multicomponent chair-based exercise (CBE) intervention on motor functioning, cognition and wellbeing for nursing home residents who are unable to walk. A two-arm single-blinded randomised controlled trial integrated N=52 nursing home residents (81 ±11 years, 63% female) assigned to a training or a wait list control group. Training duration was 16 weeks (twice a week; 60 minutes) Primary outcomes were hand grip strength, fine motor skills, cognitive performance (MoCA) and well-being (i.e. SF12, satisfaction with life, depression (CESD)). Statistics was performed using ANOVA for repeated measures. The CBE intervention improved hand grip strength, fine motor skills, cognition, and depression in the intervention group while the CG revealed decline (interaction term always p < .03; ES > 15). Satisfaction with life, physical well-being, and mental well-being remained stable while the CG, again, revealed decline (no significant time x group interaction). In addition to previous research, our study addressed an intervention that uses multicomponent CBE for residents who are unable to walk. The results support the hypothesis that despite their lower mobility and their high level of frailty nursing home residents can improve or maintain motor and cognitive resources as well as psychosocial well-being.

###### Madeleine Fricke: Feasibility of a multicomponent exercise intervention to enhance spatial navigation skills in nursing home residents

Authors: Madeleine Fricke^1^, Adele Kruse^2^, Bettina Wollesen^1^

^1^ Berlin University of Technology

^2^ University of Hamburg

Spatial abilities, including orientation in and navigation through the environment, are known to deteriorate with age (Burns, 1999; Moffat and Resnick, 2002; Iaria et al., 2009). While spatial navigation skills in familiar environments maintain relatively safe, navigation in unfamiliar environments decreases notably with increasing age (Colombo et al., 2017). Due to multimorbidity, some older persons need to be hospitalised in nursing homes and therefore experience an unfamiliar environment. Studies showed that their range of action is limited, leading to low physical activity and social interaction (Jansen et al., 2017; den Ouden et al., 2015), and, as a consequence a lack of physical and cognitive stimulation (Schrempft et al., 2019). To counteract this degradation, training elements, which intend to promote spatial navigation skills have been developed and tested for their feasibility. Until now a total of 31 persons (age range 60 to 97 years, 30 female) participated in six regular training groups, two further groups are planned. All training sessions are observed and systematically documented by an independent person, which focuses on the clarity of the instructions, the classification of the participants’ performance and motivation, and the length of execution time. The analysis of the observation protocols has led to three modification levels and will be successively adjusted, resulting in a final set of orientation training elements tailored to the target group of nursing home residents. These will be applied to a randomised, controlled intervention study in a total of 18 nursing homes to evaluate their effectiveness on spatial navigation skills.

###### Exercise and health of community-dwelling older adults (exercise before and during covid-19 pandemic)

Symposium Chair:
Prof. Dr. Vânia Loureiro, *Polytechnic Institute of Beja, Portugal*

The health benefits of physical activity are well-known, including improvements in longevity, bone mineral density, cardiovascular risk factors, aerobic fitness, muscular strength and endurance, diabetes, obesity, some cancers, osteoporosis, and mental health. Among older adults, physical activity interventions can be considered as a non-pharmacological alternative. Additionally, it´s associated with functional fitness and cognitive functions, which lead to a more independent life and better quality of life. In March 2020, the disease COVID-19 reached the pandemic dimension, infection control became imperative and safety measures were needed. Staying at home has become an essential safety behaviour but, the home isolation result in a decrease of moderate-to-vigorous physical activity levels and increase in sedentary behaviour. This nonexercise behaviours lead to an increasingly sedentary lifestyle, known to result in a range of chronic health conditions and to anxiety and depression. There was already a need to find ways to improve health in old age, but the pandemic COVID-19 emphasized the priority of exercise programs in public health policies. In the light of WHO Global action plan 2016-2025, physical activity already remains as a leading factor in health and well-being, with particular attention to the burden of noncommunicable diseases associated with low physical activity levels and sedentary behaviour. Recently the United Nations proclaimed 2021–2030 as announced the Decade of Healthy Ageing, with WHO guiding international action to improve the lives of older adults. In this symposium will be presented results of health interventions, through physical exercise (before and during COVID-19 pandemic), which we believe be undertaken to achieve specific health care goals of older adults and that should be included and prioritized in public health programs and policies.

###### Nuno Loureiro: Effectiveness of a combined exercise program to improve functional fitness in community-dwelling older adults: A randomised controlled trial

Authors: Vânia Loureiro^1^, António Cachola^2^, Tiago Rosa^2^, Adilson Marques^3^, Joanna Gradek^4^, Nuno Loureiro^1^

^1^ Polytechnic Institute of Beja, Portugal

^2^ Office of Associative Movement, Sport and Youth, Municipality of Serpa

^3^ Faculty of Human Motricity, University of Lisbon, Portugal

^4^ University of Physical Education in Krakow, Poland

Purpose: Functional fitness is defined as the ability to independently take-out daily living activities without effort. The progressive decline of functional fitness is a characteristic of the aging process. The present study aims to evaluate the impact of a combined exercise program on functional fitness in community dwelling older adults. Methods: A total of 75 community-dwelling older adults (68, woman) were assigned to a moderate to-vigorous intensity program for 2 days/week. Each participant performed a supervised combined exercise program (water fitness and multicomponent program yoga and Korean dance) for 60 min per day, 2 times per week, for 8 months. The exercise program was supervised by a certified exercise professional. The Senior Fitness Test battery and Fullerton Advanced Balance battery were measured on two different occasions. Results: Using paired t-tests, positive and significant improvements were found in %BF, in five physical fitness components of the Senior Fitness Test and in eight of the balance tests, but not in BMI, sit and reach, standing on one leg and rotation of the head. The level of significance was set top <0.05 Student’s t-test. Conclusion: The positive results that we found with this exercise program suggest that an ease and low-cost implementation could bring an effectiveness improvement on physical fitness and health of the community dwelling older adults. Keywords: physical fitness; multicomponent exercise; SFT; FAB.

###### Miguel Peralta: Grip strength and depressive symptoms among European middle-aged and older adults

Authors: Adilson Marques^1^, Priscila Marconcin^1^, Miguel Peralta^1^, Élvio R. Gouveia^2^, Nuno Loureiro^3^, Vânia Loureiro^3^, João Martins^1^, Margarida Gaspar de Matos^1^

^1^ Faculty of Human Motricity, University of Lisbon, Portugal

^2^ University of Madeira, Portugal

4 Polytechnic Institute of Beja, Portugal

Purpose: To analyse the relationship between grip strength and symptoms of depression considering sex and age in adults from 18 countries. Methods: Cross-sectional data on adults aged ≥50 years from the Survey of Health, Aging, and Retirement in Europe wave 6 (collected in 2015) were analysed. Grip strength was measured hand using a handgrip dynamometer. The EURO-D 12-item scale was used to measure depression symptoms. Multivariable logistic regression analysis was conducted. Results: Men and women who were in the 2nd, 3rd and 4th quartile of grip strength were less likely to have depression symptoms than those in the 1st quartile of grip strength. Having more grip strength decreased the odds of depression symptoms by 30% (OR: 0.70; 95% CI: 0.65, 0.77) and 47% (OR: 0.53; 95% CI: 0.49, 0.57) for adults aged 50-64 years and ≥65 years, respectively, when compared to those with the lowest grip strength. The negative relationship between strong grip strength and depression symptoms were observed among men and women under and over 65 years. Conclusion: There was an association between grip strength and depression symptoms. For clinical practice and geriatric health professionals, assessing adults’ grip strength can be used as a signal to screen mental health. Keywords: handgrip; physical function; European adults; mental health.

###### Bebiana Sabino: Postural balance as predictor of fall in community-dwelling older adults

Authors: Margarida Gomes^1^, Nuno Loureiro^1^, Bebiana Sabino^1^, Priscila Marconcin^2^, Adilson Marques^3^, Vânia Loureiro

^1^ Polytechnic Institute of Beja, Portugal

^2^ Faculty of Human Motricity, University of Lisbon, Portugal

^3^ CIPER, Faculty of Human Motricity, University of Lisbon, Portugal

Purpose: The use of force platform measures could objectively have been developed as tools for evaluating the center of pressure movement patterns of postural control, functional mobility and identify those at fall risk. Design new, more accurate and reliable diagnostic methods allow us to detect early changes of pathologies and control clinical practices based on balance exercise to fall programs. The aim of the present study was to identify determinants of postural stability according COP displacement in community-dwelling older adults. Methods: Data were normally distributed (Shapiro–Wilks statistic); for each balance measurement, independent-samples t-test and one-way ANOVA assess the mean differences between two and three groups in the following dependent variables: (1) medio-lateral (ML) displacement; and (2) anteroposterior (AP) displacement, eyes open (EO) and eyes closed (EC). Results: Experimental results showed medio-lateral displacement under the eyes-open present statistically significant differences (t(22) = 2.086; p= .049) between fallers and non-fallers. Participants categorised as “high concern about falling” had a higher medio-lateral displacement under the eyes-close (F(2;20) = 3.949; p = .036). The participants with the higher medio-lateral displacement under the eyes-close are those who perceive a high concern about falling (r = .444; p = .034). Conclusion: As a preliminary screening tool, force platforms are an effective strategy to identify impairments on postural control, which are a risk factor for forthcoming falls in older people. Keywords: balance, force platform, center of pressure, fall, elderly.

###### Priscilla Marconcin: The association of grip strength with depressive symptoms among middle-aged older adults with different chronic diseases

Authors: Priscila Marconcin^1^, Miguel Peralta^1^, Nuno Loureiro^2^, Vânia Loureiro^2^, João Martins^1^, Eugenia Murawska-Ciałowicz^3^, Adilson Marques^1^

^1^ Faculty of Human Motricity, University of Lisbon, Portugal

^2^ Polytechnic Institute of Beja, Portugal

^3^ University of Physical Education in Krakow, Poland

Purpose: Low grip strength has been associated with an increase in depressive symptoms, independent of age group or gender, although the literature has not investigated this association among different chronic diseases. The present study aims to investigate the association of grip strength and depressive symptoms among middle-aged and older adults with different chronic diseases. Methods: A cross-section of data from the Survey of Health, Ageing, and Retirement in Europe wave 6 (collected in 2015) was analysed. Grip strength was measured by a handgrip dynamometer, and the European Depression Symptoms 12-item scale (EURO-D) was used to assess depressive symptoms. Multivariable logistic regression analysis was conducted. Results: Those in the high strength tertile had 42% (95% confidence interval: 0.50, 0.71; p < 0.005) and 41% (95% confidence interval: 0.50, 0.70; p < 0.001) lower odds of depressive symptoms in the ‘no disease’ and in the ‘metabolic diseases’ groups of participants, respectively, compared with those in the lower strength tertile. No statistically significant relationship between grip strength and depression was observed in the ‘arthritis diseases’ group of participants. Conclusion: The association of grip strength with depressive symptoms must consider, besides gender and age group, the chronic conditions that an individual could have. Keywords: handgrip strength; depressive symptoms; chronic disease; SHAR.

###### Adilson Marques: Physical activity is negatively associated with depression symptoms independently of socioeconomic status

Authors: Miguel Peralta^1^, Priscila Marconcin^1^, Élvio R. Gouveia^2^, João Martins^1^, Vânia Loureiro^3^, Nuno Loureiro^3^, João SantosNone^4^, Adilson Marques^1^

^1^ Faculty of Human Motricity, University of Lisbon, Portugal

^2^ University of Madeira, Portugal

^3^ Polytechnic Institute of Beja, Portugal

^4^ Environmental Health Institute, University of Lisbon, Portugal.

Purpose: Few studies have evaluated the relationship between depression symptoms (DS) and physical activity (PA) considering peoples’ sociodemographic characteristics. This study aimed to analyse the relationship between DS and PA, stratified by sociodemographic characteristics of European adults. Methods: Participants were 29285 adults (13943 men, 47.6%; 15342 women, 52.4%), aged 50.9±17.4 (50.6±17.3 men, 51.1±17.5 women) from the European Social Survey round 7. DS was assessed with the Centre for Epidemiological Studies Depression Scale (CES-D8). Leisure-time PA (LTPA) was self-reported. The analysed sociodemographic characteristics were sex, age, living place, household members, marital status, income, and educational level. The relationship between DS and PA, stratified by sociodemographic variables, was examined by linear regression models. Results: Engaging in LTPA was negative and linearly related to DS, independently of sex, age, place of residence, having children or not, being single or married, being wealthy or poor, employment status, and having a lower or a higher education level. Age was the variable with both the least and the greatest effect of LTPA on DS. The least effect of LTPA on DS was observed in younger adults (β=-0.08, 95% CI: -0.11, -0.05) and the greatest effect in retired people (β=-0.33, 95% CI: -0.36, -0.29). Conclusion: Independently of sociodemographic characteristics, LTPA is associated with DS and can benefit everyone. Public health policies for promoting mental health should include PA promotion as an important strategy for the prevention or treatment of DS. Keywords: sport; exercise; adults; mental health; sociodemographic; European Social Survey.

###### Vânia Loureiro: Home-based exercise program and health literacy intervention during COVID-19 pandemic: Up again senior pilot study

Authors: Vânia Loureiro^1^, Margarida Gomes^1^, Luís Branco^2^, Nuno Loureiro^1^, Ana Alves^1^, Adilson Marques^3^, Margarida Gaspar de Matos^3^

^1^ Polytechnic Institute of Beja, Portugal

^2^ Health Senior Club, Municipality of Viana do Alentejo, Portugal

^3^ Faculty of Human Motricity, University of Lisbon, Portugal

Purpose: Finding ways to optimize health in older age become a priority for public health policy exercise programs. In the light of WHO Global action plan 2016-2025, physical activity remains as a leading factor in health and well-being, with particular attention to the burden of noncommunicable diseases associated with low physical activity levels and sedentary behavior. Given the concerns about the increasing spread of COVID-19, infection control was imperative and safety precautions must be followed. Staying at home has become an essential safety step that can limit infections from spreading widely. The UP Again Senior - home-based exercise program aims to demonstrate the feasibility of an individualised home-based physical activity program including exercise instructions by digital means. Methods: Maintaining regular PA and exercise routine in a safe home environment is an important strategy for healthy living during the coronavirus crisis. The UP AGAIN SENIOR implement a home-based exercise and health literacy intervention during COVID-19 pandemic with 59 community dwelling older adults (≥ 55 years). In the baseline we perform an individual multifactorial assessment, in an environment authorized by the local health delegate. In the end of the assessment, exercise routines will be provided by means of a booklet with the description of all exercises. In the mentioned booklet will be collected data related to the adherence to the program and physical activity behavior (walking time/day; Exercise routine/day). To control health status, motivation and physical performance were designed and implemented regular phone interviews (2 in 2 weeks) and in the participants register in the booklet their arterial pression and glycemia. We also adopt a multiple strategies as ordinary mail; electronic mail and a TV channel (where they can watch the exercise routine). Participants will be exercising at home for 9 months. The exercise program will be an individually tailored exercise program. Progression through the levels in the exercise program will be in consultation with a coach through weekly telephone contact and in consequence of the assessment appointments at pre, middle and post (0, 3, 6 and 9 months). Results: Primary outcome is physical performance measured and relevant secondary and observational outcomes include physical activity level, anxiety and depression symptoms and fall history. Outcome variables were measured at the beginning of the program, after 3, 6, 9 months and at the end of the 12-months of the intervention. Conclusion: The UP AGAINS SENIOR home-based exercise program aims to increase health through nonpharmacological strategy. We believe that Up Again Senior offer examples of approaches that may be implemented to reduce potential risks of COVID-19 isolation, keeping the social distancing but also promoting actions that keep the older adults active and as healthy as possible during the pandemic.

###### Relationship of physical and cognitive performance in community-dwelling older adults and nursing home residents

Symposium Chair:
Prof. Dr. Bettina Wollesen, *Berlin University of Technology, Germany* & Dr. Katrin Müller, C*hemnitz University of Technology, Germany*

In order to protect older adults from dependency on nursing care, interventions to promote physical and cognitive performance could be beneficial. Several studies with older adults revealed that a higher level of physical activity is associated with better physical and cognitive performance, reduced risk of falls as well as improved quality of life, while sedentary behavior is related to increase the risk for chronic diseases, frailty and mortality (Cabeza et al., 2019; Hajek et al., 2017). But less is known about these associations in the group of community-dwelling adults aged 80 years and older or nursing home residents. However, insights into these relationships are crucial to develop innovative, targeted interventions to support the health care system. In this context, the symposium will present results regarding interactions of physical activity, physical as well as cognitive performance among community dwelling older adults and nursing home residents. The first two presentations report on results of the SENDA-Project (“Sensor-based systems for early detection of dementia - a reaction to demographic change”). Longitudinal data of cognitive performance (memory, processing speed, language skills, and executive functions) of non-demented older adults, measured three times with intervals of eight months, describe performance levels in high-agers (>80 years) and enable to compare their trajectories. The results are embedded in the field of tension between aging processes, intra-individual variability and test repetition effects. In the second talk potential associations of physical activity, physical performance and cognitive decline in the same study population will be shown. Subsequently, daily physical activity patterns (e.g. moderate physical activity and sedentariness) of nursing home residents will be presented. In the last talk, the ability to walk as a crucial factor for independent living and quality of life in older adults will be discussed. For this, data of physical and cognitive performance of nursing home residents with and without the ability to walk of the project PROCARE (“Prevention and occupational health in long term care”) will be reported. Overall, the final discussion of the integrated presentations will summarise the implications for future health care intervention practice for older adults and nursing home residents.

###### Stephanie Fröhlich: Trajectories of cognitive performance in community-dwelling adults older than 80 years over a 16-month period

Authors: Stephanie Fröhlich^1^, Katrin Müller^2^, Claudia Voelcker-Rehage^1^

^1^ University of Münster, Department Neuromotor Behavior and Exercise, Germany

^2^ Chemnitz University of Technology, Institute of Human Movement Science and Health, Germany

Few studies present data for change in cognitive performance in high-agers (>80 years), even though this data is needed to understand the capabilities and limitations of this growing population. In addition, accelerated rates of change could be related to neuropathological processes. We recruited 244 non-demented older adults (M = 82.5 years, SD = 2.5, 123 males) as part of the SENDA study (a prospective cohort sequential study) at Chemnitz University of Technology, Germany and measured cognitive performance at baseline and follow-ups after eight (n = 162) and 16 months (n = 89). Participants without follow-ups did not differ in age or education from participants with follow-ups but had lower baseline Montreal Cognitive Assessment scores. Processing speed (Digit Symbol Substitution Test), executive function (Trail Making Test) and semantic memory (Wordlist Recall) were measured. Their trajectories were modelled with linear mixed models using age as predictor with random intercept and slope. Additional fixed effects included repetition, sex and education. Results indicated that there was no systematic practice effect for processing speed, while performance on the other two tests profited from repeated measurements. The overall trajectory of cognitive change indicated decline in processing speed (b = -0.6, t(102.24) = -2.70, p = .01) and no change in the other domains. Model fit was not increased by including a random slope for the effect of age indicating that individual trajectories could not be properly modelled with linear growth curves. The project was funded by ESF and Sächsische AufbauBank-Förderbank (Project-Number: 100310502).

###### Katrin Müller: Relationship of physical activity and cognitive performance in community-dwelling older adults and nursing home residents

Authors: Katrin Müller^1^, Stephanie Fröhlich^2^, Alexandra Clauß^1^, Claudia Voelcker-Rehage^2^

^1^ Chemnitz University of Technology, Institute of Human Movement Science and Health, Germany

^2^ University of Münster, Department Neuromotor Behavior and Exercise, Germany

Due to the lack of effective treatments of dementia or its pre-stages, early detection of mild cognitive impairment (MCI) is essential, e. g., to provide health care interventions. It is discussed whether motor decline and reduced activities of daily living (ADL) are promising prodromal markers of MCI. The objective of this study was to examine predictors for MCI in a group of community-dwelling adults 80 years and older (OA). Cognitive status was assessed by a two-step classification process using the Montreal Cognitive Assessment (MCI: score < 26) and the Consortium to Establish a Registry for Alzheimer’s Disease (MCI: at least one test ≥ 1.5 SD below standard) in 154 OA (age: M = 82.52, 75 male). Physical performance was measured by use of the Short Physical Performance Battery (SPPB). ADL was assessed with the Nürnberger-Alters-Alltagsskala. Binary logistic regression analysis also included age, sex, and comorbidities as possible predictors. Analysis (Nagelkerkes R² = .331, p < .001) indicated that SPPB (Exp(B) = 0.608 p = .006) and ADL (Exp(B) = 1.331, p = .000) predicted MCI in OA but not age, sex, and comorbidities. The results indicated that reduced ADL as well as a poorer physical performance are associated with MCI in OA. To further confirm this relationship by longitudinal analysis, measurements will be repeated every eight months up to three times. Tailored interventions might target physical performance to maintain cognitive state and to prevent dementia. The project was founded by ESF and Sächsische AufbauBank-Förderbank (Project-Number: 100310502).

###### Tina Auerswald: Application of activity trackers among nursing home residents - Daily physical activity and sedentary behaviour and feasibility

Authors: Tina Auerswald^1^, Jochen Meyer^2^, Kai von Holdt^2^, Claudia Voelcker-Rehage^3^

^1^ Chemnitz University of Technology, Department of Sport Science, Germany

^2^ OFFIS - Institute for Information Technology Oldenburg, Germany

^3^ University of Münster, Department Neuromotor Behavior and Exercise, Germany

The number of dependent elderlies living in care institutions such as nursing homes is expected to increase. Reduced mobility and mobility loss represent some of the most common risk factors for the need of care in old age. Accurate quantification of nursing home residents’ physical activity is necessary to derive consensus guidelines as well as exercise intervention studies for this frail population. Thus, the primary aim of the study was to assess physical activity and sedentary behavior of nursing home residents. Furthermore, usage behavior, usability, acceptance and motivational impact of an applied activity tracker were examined. Physical activity and sedentary behavior were measured among 22 residents (68 to 102 years) by use of a commercial activity tracker (Fitbit Zip) worn during waking hours for 77 days on average. Usability, acceptance and motivational impact of the tracker were examined using a specially developed questionnaire. Participants walked on average 1007 ± 806 steps per day and spent on average 75.8 % of their waking time sedentary. The tracker was used for 65.4% of the individual study period and acceptance rate was high (94.4%). The average steps/day increased significantly within the first five weeks of wearing the activity tracker. Participants with a usage time of ≥ 50% walked significantly more steps per day than those with a lower usage. Tested Nursing home residents were highly inactive and sedentary. The results support the feasibility of a long-term application of activity trackers to assess or even increase physical activity behavior.

###### Katharina Zwingmann: Comparison of physical and cognitive status of nursing home residents with and without the ability to walk

Authors: Katharina Zwingmann^1^, Julian Rudisch^2^, Thomas Cordes^3^, Bettina Wollesen^3^, Claudia Voelcker-Rehage^2^

^1^ Chemnitz University of Technology, Department of Sport Science, Germany

^2^ University of Münster, Department Neuromotor Behavior and Exercise, Germany

^3^ University of Hamburg, Germany, Department of Human Movement and Training Science

In Germany, 818.000 nursing home residents are living in about 14.480 nursing homes. Living in a stationary care setting and in dependency, nursing home residents consider their cognitive as well as their walking abilities as central components for a maximized self-determination and participation level. The positive impact of the ability to walk on health-related quality of life in institutionalised elderly persons has already been shown. It is, however, unclear whether other parameters differ between residents with and without the ability to walk. Therefore, the aim of this study was to reveal central differences in upper extremity physical and cognitive parameters of nursing home residents with and without the ability to walk. So far, 124 residents of six nursing homes between 57 and 98 years of age (34 male; 86 with ability to walk) at different project locations participated in the study (further participants will be included). Anthropometric data, health status, satisfaction with life, cognitive status, and manual dexterity were measured cross-sectionally. Data were analysed by univariate ANOVAs comparing the group of nursing home residents with and without the ability to walk. Groups differed significantly in age, height, physical health status (SF-12) and manual dexterity (Purdue pegboard right, left, both hands), but not in their cognitive status (MoCA). Independent walking in nursing home residents seems to serve as an important factor for the conservation of physical and psychosocial resources.

###### ADYMA: Psycho-social and behavioural impact of an adapted physical activity program for seniors living in residential-based communities

Symposium Chair:
Prof. Dr. Sebastien Chastin, *Glasgow Caledonian University, United Kingdom*

The time seniors spend in a residential based community affects both their level of physical activity and their level of sedentary behaviours (Lord et al., 2011). Therefore, maintaining significant levels of physical activity and reducing sedentary behaviours in this population confers multiple health benefits (Cunningham & Sullivan, 2020). Recent studies evaluating physical activity programs and sedentary behaviours in a residential-based community for older people reveal that the programs often have low attendance rates, with difficulty motivating residents or maintaining those who participate in these programs (Hawley-Hague et al., 2016). For this reason, effective interventions are highly needed to promote physical activity and reduce sedentary behaviours in residential-based community settings. The present symposium has three main objectives: 1. To update the knowledge of the effectiveness of interventions to reduce sedentary behaviour in older people living independently in residential-based communities. 2. To present a program to promote physical activity in senior residential-based communities based on collaboration between a scientific laboratory and an association. 3. To show the preliminary results of a multiple-residences intervention program to promote mobility in seniors. In this symposium there will be three communications: First, Sebastien Chastin will present the results of a literature review about the effectiveness of interventions aimed at reducing sedentary behaviour amongst older adults living independently in the community compared to control conditions involving either no intervention or interventions that do not target sedentary behaviour. The next two communications will focus on the articulation of an intervention research project that addresses mobility among seniors living in residential-based communities. Amélie Baghdiguian and Yannick Manet will present Adyma: a program to encourage mobility and promote healthy ageing among seniors living in residential-based communities. Two principles guide Adyma’s program: 1) Accessibility for all older people (including the most vulnerable) and 2) an evaluation that establishes benchmarks for the evolution of each person within the program. Ariane Gautier, Gonzalo Marchant and Emma Guillet–Descas will present the preliminary results of the effects of Adyma program on psychological, social and behavioural aspects.

###### Sebastien Chastin: Interventions to reduce sedentary behavior in older adults: A Cochrane systematic review of the literature

Author: Sebastien Chastin^1^

^1^ Glasgow Caledonian University, United Kingdom

Background: Older adults spend a large part of their day sitting down. High levels of this behaviour have been linked to increased risk factors for health and premature death. Objectives: This presentation is about a systematic review that evaluated the effectiveness of interventions to reduce sedentary behaviour of older adults living independently in the community compared to control conditions. Method: Different sources of information were searched up to 4 June 2020, including databases (e.g. MEDLINE), scientific article platforms (e.g., ICTRP), reference lists of articles and some authors. Randomised controlled trials (RCTs) and cluster-randomised controlled trials (Cluster-RCTs) with interventions designed to reduce sedentary time in older adults (aged 60 years and over) were selected. Data analysis included titles, abstracts and full articles. There was a data extraction, and risk of bias assessed independently. Results: There were only seven low-quality studies. The interventions decreased sedentary behaviour by ~ 40 minutes, but this is not statistically significant, and the evidence is low. Conclusion: It is not clear whether these interventions change the function or health of older adults. Only individual behaviour change intervention has been tested. We have no evidence of environmental or policy change. Studies with larger sample sizes, objective monitoring and multi-dimensional approaches are needed.

###### Amélie Baghdiguian & Yannick Manet: ADYMA: A program to encourage mobility and promote healthy aging among seniors living in residences

Authors: Amélie Baghdiguian^1^, Yannick Manet^1^

^1^ ADYMA Association, France

Program description: Adyma focuses on two issues about loss of autonomy in seniors: falls and isolation. Adyma’s mission is to develop seniors’ desire and power to act on their well-being and health to preserve their autonomy. Two principles guide Adyma’s program: 1) accessibility for all older people (including the most vulnerable) and 2) an evaluation that establishes benchmarks for the evolution of each person within the program. Adyma’s originality is that the program is based on a global approach: It is aimed at both the seniors and the professionals in gerontology. Each person is subject to a personalised mobility diagnosis that determines a specific program that mixes individual and collective coaching. Adyma relies on a three-dimensional approach: physical, social and psychological, to create synergies among the participants. It is driven to build an ecosystem of mobility all around the elderly in which every actor is involved: professionals, families, volunteers, people in the neighbourhood. Adyma provides its audiences with some dedicated human support, all graduated in Adapted physical activity and trained with Adyma’s method. Each coach draws up a specific program at the residence. Furthermore, Adyma provides training for professionals and digital tools to support the program and manage the participants’ evolution.

###### Ariane Gautier & Gonzalo Marchant: Embedding psycho-social approach in a program designed to promote mobility in seniors living in residence

Authors: Ariane Gautier^1^, Emma Guillet–Descas^1^, Gonzalo Marchant^1^

^1^ Claude Bernard University Lyon 1, France

Background: Keeping seniors mobile and independent through any physical activity type, and any duration can improve health and well-being in this population (WHO, 2020), reducing health risk (McPhee et al., 2016). For that reason, it is necessary to study mobility with an approach that combines psychological, social and contextual aspects (Kanning &Schlicht, 2008). Objectives: This project aims to characterise seniors living in residences by adopting a psychological, social and behavioural approach to evaluate and adapt a mobility program for this population. Method. A total of 55 people (89% women) aged 83 years on average (SD=8.1) answered questionnaires measuring demographical, psycho-social variables, and contextual variables. Also, an evaluation of physical activity and sedentary behaviours was carried out with accelerometers on 24 participants in that sample. Results: On average, people have been living in residence for five years, presenting on average two health problems, and practicing at least one exercise or physical activity. Personality analysis showed that participants had high levels of conscientiousness and agreeableness. In terms of physical self-esteem, participants declared low general self-esteem, endurance and strength. For social provisions, seniors presented high levels of attachment, social integration, reliable alliance, and guidance but low reassurance of worth levels. The objective measures of behaviours showed that seniors spent on average 12 hours per day sitting or lying (SD=4.9), one hour in light physical activity (SD=0.6), and 18 minutes in moderate-to-vigorous physical activity (SD=24.7). Conclusions: A program to promote mobility should tap psycho-social factors linked to physical activity and sedentary behaviours, considering that this population could be highly sedentary and that light physical activity characterises their movements.

###### Healthy aging: From the brain to the muscles

Symposium Chair:
Prof. Dr. Anita Hökelmann, *Otto-von-Guericke-University Magdeburg, Germany*

With increasing age and physical inactivity, muscle atrophy (as sarcopenia) and cognitive decline increases with age. As the body is an interactive system, the loss of muscle mass and cognitive function are related, and it is assumed to be bidirectional. The functional limitations in several physiological systems cause non-physiological age-related cognitive decline and neurological disorders contribute to sarcopenia and impaired functioning. An evaluation of cognitive interventions aiming to improve mobility in elderly and assessing clinical effectiveness on gait performance is conducted. Combining physical with cognitive activity seems to be ideal for intervening the bidirectional relation of muscles and the brain. For combination of these aspects, dancing interventions appear to be suitable; their effects on brain structure and cognition in older adults are evaluated. Tensiomyography, a sensitive tool for detecting muscle-dysfunction in early stages, is used to detect neuromuscular alterations after inactivity (bed rest) and subsequent recovery in young and old subjects. It detects alterations earlier than using ultrasound for seeking architectural changes. The importance of an active lifestyle due to early changes resulting from inactivity is stressed. To determine the muscle performance of seniors, different skeletal muscle contractile parameters are measured with tensiomyography. A comparison of neuromuscular function in healthy elderly and elderly with mild cognitive impairments (MCI) and between gender is conducted. Furthermore, the purpose is to explore neurophysiological correlates of cognitive performance and to investigate whether this multimodal approach can aid in early identification of individuals with MCI. These early stadiums of not natural cognitive decline are an important phase in which intervention strategies can be applied to minimise risk factors, to improve cognitive status and to prevent a manifestation of dementia. Since the number of persons diagnosed with dementia will continue to rise in the future due to longevity in the Western societies, neurodegenerative diseases are a main issue. Additionally, age is the major risk factor for Alzheimer’s. In this symposium the bidirectional relationship between changes in muscle mass and cognitive function is explored in order to contribute to the understanding of the body as an interactive system.

###### Anita Hökelmann & Achraf Ammar: The effects of physical activity - dance and sport intervention- on brain structure and cognition in elderly

Authors: Anita Hökelmann^1^, Achraf Ammar^1^ ,Patrick Müller^2^, Mandy Knoll^1^, Kathrin Rehfeld^1^, Notger Müller^2^

^1^ Otto-von-Guericke-University Magdeburg, Germany

^2^ DZNE Magdeburg, Germany

The positive correlation and beneficial effect of physical activity on cognitive functions was reported in the latest WHO recommendations (”Risk reduction of cognitive decline”, 2019). Several theories attempt to explain the neuroprotective and neuroplastic effects of physical activity. Kirk-Sanchez and McGough (2014) report that exercise affects neuroplasticity of the brain via two mechanisms of actions. They explain that endurance sports improve cardiovascular health and increase the level of neurotrophic factors. These stimulate neurogenesis, synaptogenesis and neuroplastic processes, resulting in improved cognitive performance. We created a 5-year study with 60 seniors with two types of intervention strategies. Our investigations have shown that 5 years of intensive coordination training, using dance sequences, increased significantly the volumes of the brain of dancers more than those in endurance and strength training groups, using repetitive movements. The process started already after a training period of 6 months and lasted over 5 years. The increase of volume can be registered in white and grey matter parts of brain (FWE p>.05). Additionally, an improvement of cognitive abilities and balance was found. The cardiovascular fitness levels of both groups remained constant over the time course of the interventions. Dance movements require the simultaneous coordination of several parts of the body in different directions and adjustment to the varying rhythms of the music (polycentric and polyrhythmic). These features included also the requirement to constantly learn new choreographies (i.e., memory), integrate multisensory information, coordinate the entire body, and navigate in space. We assume that a long-term dancing/ coordination intervention could be superior to repetitive physical exercise in inducing neuroplasticity in the human brain. We presume that this advantage is related to the multimodal nature of dancing, which combines physical, cognitive and coordinative challenges.

###### Rado Pišot: Early neuromuscular alterations in older compared to younger men following prolonged bed rest and subsequent recovery

Authors: Rado Pišot^1^, Boštjan Šimunič^1^, Marco Narici^1,2^, Carlo Reggiani^1,2^, Gianni Biolo^3^

^1^ Science and Research Centre Koper, Slovenia

^2^ Department of Biomedical Sciences, University of Padova, Italy,

^3^ Department of Medical, Surgical and Health Sciences, University of Trieste, Italy

Purpose: Physical inactivity induces rapid declines in musculoskeletal, cardiovascular and metabolic systems. Decline strongly depends on participants age and fitness. The use of physical inactivity analogues (e.g bed rest, immobilization, suspension, dry-water immersion) is very popular, where bed rest is a gold standard. However, only few institutions could develop bed rest study in a complex hospital setting. Institute for Kinesiology Research conducted a series of bed rest studies and were the first to compare declines in younger and older adults after 14-day bed rest. The aim of this presentation will be an overview of neuromuscular changes after periods of bed rest with an emphasis in comparing younger and older participants as well as presenting early cascade of declines. Methods: Younger and older men underwent 10- to 35-day bed rest in Valdoltra orthopaedic hospital (2006-08, 2012) and General Hospital Isola (2019). Skeletal muscles were assessed at a single fibre level (size, force, tension) and whole muscle level (size, force, power, contraction time and displacement, architecture) and function (gait, balance). Furthermore, early biomarkers of neuromuscular junction were assessed in blood or muscle biopsies (NCAM – neural cell adhesion molecule, CAF – C-terminal agrin fragment). Results: Findings suggest severe muscle atrophy in leg muscles (~20%) after 35-day bed rest, being largest fast twitch fibres in postural distal muscles. Even higher declines in muscle force and power could not be explained solely by declines in muscle size and architecture. About 25% of the muscle force decline variance must be explained by neural factors. Being even higher in muscle dynamic actions (e.g. explosive power). Muscle size declines after a week of bed rest, whereas contractile properties decline already after 24-72 hours of bed rest. Changes in older participants are preceding those in young; and even more importantly, declines in older are less reversible after the bed rest. Therefore, in our latest study an emphasis was given to study early biomarkers and cascade of muscle atrophy, e.g. neuromuscular junction. Initial and partial myofiber denervation was observed from increased NCAM positive fibers and increased CAF concentration was observed after 10-day bed rest. Conclusion: Our findings suggest that older participants decline at a faster rate with slower recovery, which is of highest clinical importance. Furthermore, only few days of unloading are enough to induce a decline of skeletal muscle mass and function were the muscle force is lost at a faster rate than mass. In contemporary society physical inactivity is becoming very important public health problem inducing a need to study its consequences at early stages as well to design intervention programmes to counteract those changes. Facing time of the lock down because of COVID – 19 the problem of inactivity is becoming new reality.

###### Boštjan Šimunič: Age-related slowing of skeletal muscles in non-athletes, power and endurance master athletes

Authors: Boštjan Šimunič^1^, Rado Pišot^1^, Jörn Rittweger^2^, Hans Degens^3^

^1^ Science and Research Center Koper, Slovenia

^2^ Institute of Aerospace Medicine, German Aerospace Center, Cologne, Germany

^3^ School of Healthcare Science, Manchester Metropolitan University, Manchester, United Kingdom

Purpose: In nonathletes muscle mass shows a progressive decline of as much as 1% to 1.5% per year after about the age of 50. The age-related loss of muscle mass is a consequence of loss and atrophy of muscle fibers. Since many of the age-related changes in skeletal muscle are similar to those induced by disuse it is likely that the decrease in physical activity in old age is a major contributor to the muscle wasting during aging. Master athletes, however, maintain high levels of physical activity and suffer from fewer morbidities, thereby providing a unique human research model to disentangle the effects of disuse and comorbidities from aging per se. There are indications of disproportional age- and physical inactivity-induced muscle wasting between muscles. Therefore, the aim of this study was to assess age-related changes in the Tensiomyographic contraction time (Tc) of the vastus lateralis (VL), gastrocnemius medialis (GM), and biceps femoris (BF) muscles in older non-athletes and master athletes. Methods: Tc was assessed in VL, GM and BF muscles in older non-athletes (age = 62.1 ± 12.7 years; males n = 133; females n = 246), power (age = 56.9 ± 13.5 years; males n = 100; females n = 78) and endurance master athletes (age = 56.5 ± 14.5 years; males n = 76; females n = 73). Furthermore, in a subsample of 17 master athletes we obtained VL biopsies for myosin heavy chain (MHC) determination. Results: We found an age-related slowing in all muscles, irrespective of discipline, where endurance master athletes had the longest and power master athletes had the shortest Tc. Our findings were confirmed also by MHC estimation, where power athletes had lower amount of MHC type 1 than endurance athletes. Conclusion: TMG revealed that the age-related slowing of muscle contractile properties occurs particularly in endurance athletes. Here we suggest that this may be related to their high proportion of type I and IIa fibers that have been reported to exhibit an age-related slowing independent of shifts in myosin heavy and light chain composition.

###### Uroš Marušič: The role of enhanced cognition for mobility improvements: A neurophysiological perspective

Authors: Uroš Marušič^1,2^, Rado Pišot^1^

^1^ Science and Research Center Koper, Slovenia

^2^ Alma Mater Europaea – ECM

Prolonged immobilisation or inactivity following injury and/or surgery can lead to motor dysfunction, which prevents rapid recovery and increases costs to the national health care system. It is therefore important to develop interventions that reduce the deterioration of motor performance during immobilisation or inactivity. In three different studies (randomised controlled trials, RCTs), we tested the effectiveness of non-physical training to mitigate the deterioration of motor performance (i.e. walking efficiency), focusing on neurophysiological adaptations. In two RCTs, healthy community dwelling older adults underwent either a 24-session or 12-session of computerised cognitive training under usual or bed rest conditions, respectively. In third RCT, patients who were scheduled to total hip arthroplasty surgery, underwent a 24-session of non-physical motor imagery training. The application of non-physical training was feasible and was generally well received by the older individuals, who had no experience with such procedures. The addition of non-physical training had a positive effect on distal untrained domain, such as mobility. More specifically, mobility-related improvements were accompanied by improvements in neurophysiological parameters (shorter P200 latency) and blood biomarkers (concentrations of plasma Brain-derived neurotrophic factor (BDNF)). In summary, non-physical approaches provide an affordable, safe and not very time-consuming tool to optimize the individual functional training or rehabilitation process. It allows the start of training of very frail populations or the rehabilitation process in the early post-operative phase, when the patient is not able to perform regular physical training. While future studies must aim to investigate the long-term effects, the use of non-physical training is recommended, especially when the extent and scope of physical training is limited or impossible.

###### Bernhard Grässler: Neurophysiological correlates of cognitive performance in elderly: Heart rate variability as part of multimodal measurement approach

Authors: Bernhard Grässler^1^, Anita Hökelmann^1^

^1^ Otto-von-Guericke-University Magdeburg, Germany

Heart rate variability (HRV) describes the beat-to-beat fluctuations in the ECG and is considered as a marker of autonomic function. Deteriorations in specific brain areas belonging to the central autonomic network (CAN) provokes cognitive and cardiac autonomic impairment. Whether autonomic dysfunction is present prior to a cognitive decline, or is only a symptom of cognitive decline, is still a matter of debate. Thus, the aim of the present study was to examine HRV of 22 MCI subjects and 29 healthy controls (HRV). Linear (RMSSD and HF) and non-linear indices (SD2, D2, and DFA1) were analysed. These parameters were chosen, as they reflect vagal activity. According to literature, we assumed a relation between high vagal activity and better self-regulation, adaptation, and cognitive functioning. HRV analysis was performed on a 10-minute electrocardiographic recording in sitting position with free breathing. The last five minutes were used for analysis. Our results showed significant differences between MCI and HC in SD2 (24.24 ±11.14 vs. 35.03 ±18.07), D2 (0.47 ±0.92 vs. 1.20 ±1.42), and DFA1 (0.84 ±0.33 vs. 1.05 ±0.29). The differences between MCI and HC for RMSSD (25.05 ±10.49 vs. 27.57 ±15.13) and HF (162.00 ±107.28 vs. 283.70 ±295.74) were not significant. Our results suggested autonomic deficits in MCI compared with HC. The results were significant for non-linear, but not for the traditional linear parameters, suggesting a reduced complexity of cardiac rhythm in MCI. More research is needed to determine the prognostic relevance of HRV as a predictor of cognitive decline.

###### Nicole Halfpaap: Neuromuscular functions of MCI patients in comparison to healthy seniors - muscle contractile properties measured with Tensiomyography

Authors: Nicole Halfpaap^1^, Berit Kristin Labott^1^, Achraf Ammar^1^

^1^ Otto-von-Guericke-University Magdeburg, Germany

Introduction: The musculature is an important organ system for locomotion of humans in sporting as well as everyday life. With advancing age, people lose muscle mass and performance (Rodriguez-Ruiz, D. 2013). In bed rest studies by Simunic et al. (2019), early atrophic processes could be determined by means of tensiomyography (TMG), especially in the muscle belly displacement (Dm), even before a change in the muscle composition becomes visible (Simunic, B. 2019). In this study, TMG was used to identify possible early features in seniors with mild cognitive impairment (MCI) in muscle performance. Method: During the study, the focus was on the TMG parameters contraction time (tc), muscle belly displacement (Dm) and reaction time (td) of certain muscles. In this cross-sectional analysis, the conditions of healthy seniors (n=33) aged 68,9 ± 6,9 years and seniors with MCI (n=28) aged 71,3 ± 6,1 years were compared. Results: Significant differences between groups could be found in m. semitendinosus left in tc (p= .047), m. tibialis anterior (TA) left in td (p= .020), TA right in tc (p= .044) and m. flexor digitorum right in tc (p= .035). Conclusion: TMG appears to be a well-functioning method for determining differences in contractile properties of the measured muscles between healthy seniors and seniors with MCI. In particular, the contraction time and reaction time are important parameters that change early on MCI. Keywords: Tensiomyography, muscle performance, MCI. References: Rodriguez-Ruiz, D. (2013): Effects of age and physical acivity on response speed in knee flexor and extensor muscles, Eur Rev Aging Phys Act (2013) 10:127-132. Simunic, B. (2019): Tensiomyography detects early hallmarks of bed-rest-induced atrophy before changes in muscle architecture. J Appl Physiol 126: 815–822, 2019; doi:10.1152/japplphysiol.00880.2018.

###### Life-space mobility in old age

Symposium Chair:
Dr. Eleftheria Giannouli, *Division of Sports and Exercise Medicine, Department of Sport, Exercise and Health, University of Basel, Switzerland &* PD Dr. Timo Hinrichs, *Division of Sports and Exercise Medicine, Department of Sport, Exercise and Health, University of Basel, Switzerland*

Mobility is broadly defined as the ability to move – either by walking, if necessary supported by aids, or by using means of transportation. Important aspects of a person’s mobility in real life are captured by the so-called life-space mobility. Life-space mobility describes the physical and social environment which a person visits in real life, which also distinguishes between in- or out-of-home mobility. Hence, it is defined as the space in which a person moves in daily life (e.g. home, neighbourhood, city or beyond) while also considering frequency of travel and need of assistance. In that sense, mobility encompasses both a person’s independent mobility, requiring mobility-related physical activity (e.g. walking), as well as all movement supported by mobility aids and/or means of transportation. Particularly in old age, life-space mobility is very important for social participation, but it has also been shown that it is closely connected with a person’s health status, physical and cognitive functioning and quality of life. That is why a restricted life-space mobility is regarded as a red flag for physical disability and even a higher mortality risk. Given the importance of life-space mobility for active and healthy aging, this symposium will present recent reviews as well as research articles addressing a wide range of topics related to life-space mobility. The symposium will start with a scoping review about subjective and objective assessment tools and frequently reported life-space mobility outcomes (Eleftheria Giannouli), followed by an analysis of multidisciplinary factors within the motility framework and their relationship with life-space mobility in frail older adults (Julia Seinsche). The symposium will further cover intervention approaches to increase life-space mobility in community-dwelling older adults and nursing home residents (Carl-Philipp Jansen), and close with life-space mobility of older adults before and during Covid-19 (Erja Portegijs).

###### Eleftheria Giannouli: Life-space mobility in patients with neurological diseases: A scoping review

Authors: Eleftheria Giannouli^1^, Andrea Roffler^1^, Roland Rössler^1^, Timo Hinrichs^1^

^1^ Division of Sports and Exercise Medicine, Department of Sport, Exercise and Health, University of Basel, Switzerland

Life-space (LS) mobility is an indicator of health status, physical and cognitive functioning and quality of life. Motor and cognitive impairments that are often evident in neurological conditions can cause a reduction of LS mobility, which in turn can lead to depression, loneliness and other adverse outcomes. Therefore, valid and reliable tools are essential for early risk detection and for the evaluation of interventions aiming at improving LS mobility. Numerous subjective and objective assessment tools and outcomes have been reported. Apparently, there is no consensus on “standard tools and outcomes, hindering the comparability of studies. Aiming to provide an overview of tools and outcomes used to assess and quantify LS mobility in patients with neurological conditions, a systematic review was conducted. PubMed, Scopus and Cinahl were searched using combinations of keywords consisting of the term “life-space”, six predefined neurological diseases and potential subjective and objective assessment methods. 41 studies fulfilled the inclusion criteria: reporting at least one quantitative outcome and including participants suffering from at least one of the predefined diseases. Most studies were conducted in patients with mild cognitive impairment and stroke, only few in patients with dementia, Parkinson’s disease, multiple sclerosis and polyneuropathy. The vast majority of studies using subjective assessments used the LS Assessment questionnaire. Studies using objective assessments mostly used the Global Positioning System (GPS), however, without being consistent in methodology and reported outcomes. For future studies using GPS, a consensus on methodology and outcomes is needed, facilitating the comparability across studies and patient groups.

###### Julia Seinsche: Motility in frail older adults: First insights into its relationship with physical activity and life-space mobility

Authors: Julia Seinsche^1^, Wiebren Zijlstra^1^, Eleftheria Giannouli^1^

^1^ Institute of Movement and Sport Gerontology, German Sport University Cologne, Germany

In order to design effective interventions to prevent age-related mobility loss, it is important to identify influencing factors. The concept of “motility” by Kaufmann et al. subdivides such factors into three categories: “access”, “skills”, and “appropriation”. The aim of this study was to assemble appropriate quantitative assessment tools for the assessment of these factors in frail older adults and to get first insights into their relative contribution for life-space and physical activity-related mobility. This is an exploratory cross-sectional study conducted with twenty-eight at least prefrail, retired participants aged 61–94. Life-space mobility was assessed using the “University of Alabama at Birmingham Life-space Assessment” (LSA) and physical activity using the “German Physical Activity Questionnaire” (PAQ50+). Factors from the category “appropriation”, followed by factors from the category “skills” showed the strongest associations with the LSA. Factors from the category “access” best explained the variance for PAQ50+. This study’s findings indicate the importance of accounting for and examining comprehensive models of mobility. The proposed assessment tools need to be explored in more depth in longitudinal studies with larger sample sizes in order to yield more conclusive results about the appropriateness of the motility concept for such purposes.

###### Carl-Philipp Jansen: Intervention effects on life-space mobility: Current evidence and future perspectives

Author: Carl-Philipp Jansen^1^

^1^ Robert-Bosch-Krankenhaus Stuttgart, Germany

Background: Although the association of life-space mobility with various clinically meaningful measures in older populations such as apathy, depression, social participation, and physical functioning has been shown, there is limited evidence on intervention effects on life-space mobility outcomes. In the recent past, several approaches to improve life-space mobility have emerged, also using sophisticated, technology-based assessment methods. Methods: This talk gives an overview of empirical findings on interventions to improve life-space mobility across different senior cohorts, derives missing pieces in the current evidence, and proposes future steps to fill these gaps. Results and Discussion: Interventions to promote life-space mobility are scarce but show that it can be enhanced through functional training and physical activity promotion, especially in samples with mobility impairment. In community-dwelling populations, GPS-derived data provide a clear picture of the extent of older individuals’ mobility exertion in everyday life and with high ecological validity. Outcomes such as number of trips, ellipse area, and maximum distance travelled help understanding where older people move and how they are actually utilizing their individual environment. However, not all of these parameters are responsive to intervention measures. When it comes to more impaired samples such as nursing home residents, other measures are applied with much more measurement intricacies and risk of bias regarding intervention effects. Recently developed methods and instruments as well as the anticipation of ongoing technological development strengthen the usefulness and feasibility of technology-based life-space assessments in gerontological research, which comes with high potential for future evaluation of intervention effects.

###### Erja Portegijs: Older adults´ activities in the life-space before and during COVID-19 restrictions

Authors: Erja Portegijs^1^, Kirsi Keskinen^1^, Essi-Mari Tuomola^1^, Timo Hinrichs^2^, Milla Saajanaho^1^, Taina Rantanen^1^

^1^ Faculty of Sport and Health Sciences and Gerontology Research Center, University of Jyväskylä, Finland

^2^ Division of Sports and Exercise Medicine, Department of Sport, Exercise and Health, University of Basel, Switzerland

Background: In response to the COVID-19 epidemic threat, governments implemented emergency regulations restricting residents’ opportunities for activity participation and commending keeping distance from other people. Map-based questionnaires (MQ) may be used to assess spatial aspects of life-space mobility and nature of activities. We studied how older adult’s actual activity destinations (obtained through MQ; counts, frequency of visitation, and distance from home) changed after two months of COVID-19-related regulations compared to two years prior. Methods: Prospective analyses of 75-, 80-, and 85-years-old AGNES participants. At baseline, 901 participants completed the MQ, and a slightly better functioning subsample (n=44) also completed it during COVID-19. Activity destinations were any destinations for physical exercise, destinations facilitating one’s outdoor mobility, and destinations for other activities, which participants reported to have visited several times during the past month. Activity destinations were located on a map and distances between participants’ homes and destinations were computed. Results: At baseline and follow-up, activity destination variables correlated positively with life-space mobility scores (Life-Space Assessment, R=0.1-0.5, p<.05, n=901). During COVID-19, participants reported fewer activity destinations (median=4.0, IQR=5.0) than pre-COVID-19 (median=7.0, IQR=3.0; GEE Poisson loglinear p<.001, n=44), and destinations were typically located closer to home. At baseline, a variety of activity destinations was reported, but during COVID-19, predominantly destinations for physical exercise were. Conclusions: During COVID-19, more activities were done close to home than pre-COVID-19. Older adults seemed to have diligently implemented guidelines to refrain out-of-home activities other than outdoor physical activity, and limited other activities to those necessary for daily life.

### ORAL PRESENTATIONS

#### Physical activity and cognitive functioning

###### Wouter Vints: Resistance training and cognitive aging: Muscle-brain crosstalk

Authors: Wouter Vints^1,3^, Oron Levin^1,2^, Nerijus Masiulis^1^, Mindaugas Kvedaras^1^

^1^ Lithuanian Sports University, Kaunas, Lithuania

^2^ KU Leuven, Department of Movement Sciences, Movement Control & Neuroplasticity Research Group, Heverlee, Belgium

^3^ Adelante Centre of Expertise in Rehabilitation and Audiology, Hoensbroek, Netherlands

Physical activity is known to increase brain health. In the elderly population, it can delay or even reverse age-related neurodegeneration and cognitive decline, as well as improve the outcome of disorders like Alzheimer’s disease. It was recently proposed that this exercise-effect results in part from a muscle-brain endocrinal loop. During exercise muscles secrete more than 600 myokines, of whom some might cross the blood-brain barrier and interact with the brain. However, studies examining these myokines behind the blood-brain barrier are limited. Therefore, we will measure a wide range of blood and brain biomarkers before and after a 12-week resistance training program or no training in older adults. Data collection from 70 older adults with intact cognitive functioning or mild cognitive impartments (MCI) began in August 2020 and is expected to last until March 2021. Neurometabolites will be measured with proton magnetic resonance spectroscopy (1H-MRS) and brain volume will be assessed with magnetic resonance imaging (MRI). Cognitive function will be scored with the Automated Neuropsychological Assessment Metric (ANAM) and the Montreal Cognitive Assessment (MOCA). Training induced changes in levels of brain metabolite (e.g., N-acetylaspartate - NAA and myoinositol - mIns), muscle metabolic markers (e.g., PGC1α), brain-derived neurotrophic factor (BDNF), and cognitive performance scores of the studied groups will be examined. We will be the first to measure both resistance training-induced blood and brain biomarker levels in a single study in an elderly population. The neurochemical pathways that are targeted by exercise in the brain will be discussed in depth.

###### Bohumila Krčmárová: The influence of dance on the quality of life of older females

Authors: Bohumila Krčmárová^1^; Dominika Ivaničová^1^; Matúš Krčmár^1^

^1^ Constantine The Philosopher University in Nitra, Slovakia

The research aimed to investigate the effect of a 12-weeks dance program on health-related quality of life (HRQOL) in older females. 28 untrained healthy older females were participated in the program (age: 68.5 ± 4.4 years; weight: 73.7 ± 12.4 kg; BMI: 27.1 ± 4.1 kg/m2). The participants were randomly allocated into control (CON, n=14) and experimental (EX, n=14) groups. The CON was asked to maintain the same lifestyle during the time of the program. The HRQOL was measured by a standardised questionnaire SF-36 at the beginning of the experiment and after completing the 12-weeks dance intervention. The HRQOL questionnaire was evaluated using a standard scoring manual. Data normalisation was assessed using the Shapiro-Wilk test and intragroup differences were evaluated using Wilcoxon T-test and the rank test. Intergroup differences were evaluated by Mann-Whitney U-test. The results showed a significant improvement (p≤0.05) of EX in RP domains (restriction of activities due to physical health), BP (pain), GH (general health), VT (vitality), MH (mental health), PHS (overall physical health), MHS (overall mental health) and SF36 (overall quality of life). There were no significant changes in the CON. When comparing absolute values between the CON and EX groups, statistically significant results with p≤0.05 were obtained in BP, VT, MHS, and SF 36 and with p≤0.01 in GH, MH, and PHS domains. The results of the study showed that a 12-weeks dance intervention had a significantly positive impact on the HRQOL in older females. This article is part of the VEGA project 1/0351/20 “The influence of dance, strength training, and their combination on cognitive functions, quality of life, functional fitness and level of motor skills of older females.”

###### Oron Levin: Motor and cognitive inhibition in high and low fit old and young adults

Mona Abu Rukun-Halaby^1,2^, Oron Levin^3^, Ronnie Lidor^1,2^, Michal Arnon^1^, Yael Netz^1^

^1^The Academic College at Wingate, Israel

^2^Haifa University, Israel

^3^Katholic University Leuven, Belgium

Background: Structural and functional changes in the brain with advancing age affect inhibitory mechanisms. Physical fitness is known to attenuate these changes, but more is known on cognitive than on motor inhibition. Aim: To assess whether fitness is a moderator of both inhibitions in old and young adults. And to examine the link between cognitive and motor inhibition. Method: Sixty individuals aged 60+, and 30 aged 22-30 performed a maximal exercise test for assessing their fitness (predicted peak VO2). Their cognitive and motor inhibition were assessed by vocal-Stroop-test and by a new multi-limb task (CRT) respectively. Data Analysis: Two-way ANOVAs (age X fitness) were performed for assessing differences in motor and cognitive inhibitions. Pearson correlation examined the relationships between both forms of inhibition. Results: Both age (F=16.2, p=0.000) and fitness (F=8.94, p<0.004) were significant in Stroop (Interference). In the CRT, significant main effects were indicated on age (F=12.0, p<0.001), but not on fitness (F<1). For both tasks, number of errors old > number of errors young, and number of errors of low fit individuals > number of errors of high fit individuals mainly in the old. No significant age x fitness interaction (both F<1.73, p>0.1). Conclusions: Associations between cognitive and motor inhibitory processes are demonstrated mainly in the older adults. Physical fitness is a moderator of inhibition in old more than in young and in cognitive than in motor inhibition. Parallel declines of cognitive and motor inhibition with age suggest that both inhibitory processes are mediated by overlapping neural pathways.

###### Soledad Ballesteros: The effects of combined physical and cognitive training on cognition in healthy older adults: A systematic review and three-level meta-analysis

Authors: Soledad Ballesteros^1^, Jennifer Rieker^1^, Mónica Muiños^2^, José Manuel Reales^1^

^1^ Universidad Nacional de Educatión a Distancia, Spain

^2^ Universidad Internacional de Valencia, Spain

Both, physical exercise and cognitive training help to maintain cognition in the older mind. The question is whether combined physical and cognitive training might produce synergistic effects. I will present the results of a systematic review and a three-level meta-analysis conducted to assess the effects of combined training in comparison to each of its training components alone on cognitive and physical functions in healthy older adults. The effects of combined (cognitive+physical) training and cognitive and physical training alone in healthy older adults were systematically reviewed using MEDLINE, PsycInfo, and Cochrane Central Register of Controlled Trials (CENTRAL) databases to identify the relevant studies. We calculated 1130 effect sizes from 54 published intervention studies. Combined training produced increased improvement in executive functions, whereas combined training and cognitive training alone produced similar positive effects on attention. The effects on memory, language, processing speed, and global cognition did not differ as a function of the training regime. Multi-domain training was more effective when the two training components were delivered simultaneously rather than sequentially or on separate days. Both combined and physical training produced significant and greater improvements in fitness than cognitive training. We concluded that combined training is more effective than single-domain training in improving executive functions in older adults, particularly when the two components are delivered simultaneously.

###### Karolina Talar: Augmenting the effects of aerobic training by transcranial direct current stimulation on cognition in older adults with and without cognitive impairment: A systematic review of randomised controlled trials

Authors: Karolina Talar^1^, Martijn van Haren^2^, Miguel Fernandez del Olmo^3^, Tibor Hortobágyi^2^, Ewa Kałamacka^1^

^1^ Institute of Basic Sciences, Faculty of Rehabilitation, University of Physical Education in Krakow, Poland

^2^ Center for Human Movement Sciences, University of Groningen, University Medical Center Groningen, Groningen, Netherlands

^3^ Area of Sport Science, Faculty of Sports Sciences and Physical Education, Center for Sport Studies, King Juan Carlos University, Madrid, Spain

Background: Aerobic training (AE) may slow age-related cognitive decline. However, such cognition-sparing effects are not uniform across cognitive domains. The unexplored possibility exists that a priming of brain cognitive networks that are less responsive to aerobic exercise by non-invasive brain stimulation (NIBS) could potentiate the effects of AE on cognition. Indeed, transcranial direct current stimulation (tDCS), a non-invasive and safe therapeutic form of NIBS, is knows to augment the effects of cognitive training on cognition in older adults with mild cognitive impairment (MCI). Aim: The purpose of this systematic review is to determine if tDCS could augment the effects of AE on cognitive function in older adults with and without cognitive impairment. Methods: Using a PRISMA-based systematic review, we compiled studies that examined the effects of AE alone, tDCS alone, and AE and tDCS combined on cognitive function in older adults with and without MCI. Using appropriate key words and syntax in PubMed, Scopus and Web of Science searches up to December 2020, we focused on ‘MoCA’, ‘MMSE’, ‘Mini-Cog’, ‘GPCOG’ (measures) and ‘cognition’, ‘cognitive function’, ‘cognitive’, ‘cognitive performance’, ‘executive function’, ‘executive process’, ‘attention’, ‘memory’, ‘memory performance’ (outcome terms). We included randomised controlled trials in humans if available in English full text over the past 20 years, with participants’ age over 65. We assessed the methodological quality of the included studies by the Physiotherapy Evidence Database (PEDro) scale. Results: In older adults without MCI, AE alone, conducted for up to 26 weeks, improved four domains of cognitive function in a range of effect sizes of 0.10 to 0.67. Likewise, tDCS alone, conducted for up to 14 sessions, improved 4 domains of cognitive function in a range of effect sizes of 0.14 to 0.86. In older adults with MCI, AE alone, conducted for up to 8 weeks, improved 3 domains of cognitive function in a range of effect sizes of 0.11 to 0.47. In older adults with MCI, tDCS alone, conducted for up to 30 sessions, improved 3 domains of cognitive function in a range of effect sizes of 0.27 to 1.79. In neither age group were there any studies yet that have examined our hypothesis that tDCS would potentiate the effects of AE on cognition in the two age groups. Conclusion: Aerobic exercise training with tDCS intervention in healthy and impaired older adults provided evidence that tDCS has the potential for augmenting the effects of AE on number of domains of cognition. The use of both techniques may not only prevent cognitive deterioration in the older adults, but also bring many benefits in neurorehabilitation. Key words: Ageing; Aerobic training; tDCS; cognition; Systematic review.

#### Physical activity in clinical settings

###### Tim Stuckenschneider. Disease-inclusive – Time to rethink exercise classes?!

Authors: Tim Stuckenschneider^1^, Vera Abeln^1^

^1^ German Sport University Cologne, Germany

Age is a dominant risk factor for diseases such as Alzheimer’s and Parkinson’s disease (PD). A physically active lifestyle and exercise are recommended not only to prevent but also to slow down and/or alleviate symptoms of the aforementioned diseases. As such, exercise needs to be accessible to older adults with or without disease, which may be particularly challenging in rural areas with limited classes offered. Disease-inclusive exercise classes may present a solution, however, need to be analysed regarding their efficacy for all individuals independent of their health status.
In this feasibility study, 24 older adults (mean age 70.1 ± 5.7) with (n=10) or without PD (n=14) were enrolled in an eight-week multimodal exercise training and participated in at least two supervised exercise classes a week. Physical fitness, cognitive performance and quality of life were assessed before and after. Outcomes were compared across the time and between groups using repeated measures ANOVA. Data analysis showed no differences for quality of life or cognitive performance. Physical fitness, displayed as 6-minute walking test (p<0.001) and 30s chair rise (p=0.001), improved significantly after the intervention. Both, individuals with and without PD, significantly increased walking distance (healthy: p=0.005; PD: p=0.021), and increased the number of 30s chair rising (healthy: p=0.009; PD: p=0.001). An eight-week multimodal exercise intervention was effective in improving physical fitness in individuals with or without PD. Therefore, it may be time to rethink exercise classes to increase accessibility for everybody and to set an example for an open and inclusive society.

###### Mona Ahmed: Users' and other stakeholders' needs in the development of a personalised integrated care platform (PROCare4life) for older people with dementia or Parkinson disease

Authors: Mona Ahmed^1^, Mayca Marin^2^, Raquel Bouça-Machado^3^, Daniella How^1^, Elda Judica^4^, Peppino Tropea4, Ellen Bentlage^1^, Michael Brach^1^

^1^ Institute of Sport and Exercise Sciences, University of Münster, Germany

^2^ Asociación Parkinson Madrid, Madrid, Spain

^3^ Campus Neurológico Sénior, Torres Vedras, Portugal

^4^ Department of Neurorehabilitation Sciences, Casa di Cura del Policlinico, Milan, Italy

Background: PROCare4Life (Personalized Integrated Care Promoting Quality of Life for Older Adults) is a European project that proposes an integrated, scalable, and interactive care platform targeting older people suffering from neurodegenerative diseases, their caregivers, and socio-health professionals, aiming to improve the care coordination and the quality of life of these target groups. This will include physical activity recommendations. Objective: To investigate users’ needs and requirements regarding the proposed platform. Methods: Mixed qualitative and quantitative study design, including 2 online surveys, 40 interviews, and 4 workshops. Main results: The study included over 200 participants; 90 patients diagnosed with dementia or Parkinson disease, 77 caregivers, 20 socio-health professionals, and 30 other stakeholders, from Germany, Italy, Portugal, Romania, and Spain. Mode values regarding patients: age rang 65-85 years, 33% diagnosed in the last 5 years, 51% had co-morbidities, 63% received help from informal caregivers, 95% home-dwelling, 64% with partner. Health: 38% as fair, 27% had emotional problems which restrained 30% of them from doing daily activities. 64% stated that their physical health was limiting from carrying physical (64%) or daily (67%) activities. Caregivers: 46-75 years old (66%), informal (90%), insufficient time to take care of themselves (32%). Stiffness, loss of balance and gait problems of the patients were the most symptoms consistently concern both patients and caregivers. 83 % of the caregivers use smart phones, while patients mostly do not use technology foreseen for the platform. 60% of the patients and 72% of the caregivers are willing to use the proposed platform. Results were supported by interviews and applied in the workshops. Discussion: Important hints for usage of technology and designing the new platform were received. The methodology proved to be viable in terms of user-centred design of the platform.

###### Michael Brach: Exercise groups for long-term cardiac rehabilitation in Germany: Feasability and acceptance survey three month after changes in medical care

Authors: Michael Brach^1^, Mascha Nottelmann^1^, Johanna Dransmann^1^, Vanessa Hübert^1^, Klaus Völker^1^, Shu Ling Tan^1^

^1^ Institute of Sport and Exercise Sciences, University of Münster, Germany

Background: Exercise groups are one of the most important elements of cardiac rehabilitation. In Germany, a supervising physician has to be present during each group session, which is a limiting factor in meeting the increasing demand for cardiac rehabilitation groups. The National Paralympic Committee Germany, responsible for rehabilitation sports, has developed a “supervisor conception”, including alternative versions of emergency care. Attending physicians are substituted by paramedics (v1), by a physician on call (v2), or by a specially educated instructor (v3). The conception has been implemented during a feasibility project in two German federal states, Saxony and Lower Saxony. Acceptance, perceived safety and feasibility of the alternatives and of the supervisor conception as a whole have been studied in comparison to the conventional physician attendance. Methods: Based on three focus group discussions, questionnaires for participants, instructors and organisers of the exercise groups, and interview guidelines for physicians have been developed. Criteria of feasibility and acceptance have been discussed and agreed in advance. These included strong rejection as main parameter, which characterised low trust in the different ways of emergency care and was defined by marks of the two outer points of uni- and bipolar Likert-type items. Measurements were taken three months after implementation of the different forms of medical care. Results: 15 clubs with 42 heart groups took part in the study. Questionnaires were retrieved from 446 participants, 30 instructors, 21 organisers, and 13 paramedics. 14 physicians were interviewed. Percentage of participants rejecting strongly the core questionnaire items were 8.9 (v1), 9.2 (v2), 18.8 (v3). Rejection rates of the other players range from 0% to 29.2%, with the exception of paramedics judging on v3 (50% strong rejection). Conclusion: For the main criteria, strong rejection below 50%, was fulfilled for v1 and v2, these alternatives should continue to be pursued. Emergency care by trained exercise instructors (v3) needs further analysis.

###### Sylwia Mętel: Chest mobility of older adults participating in speleotherapy combined with pulmonary rehabilitation

Authors: Sylwia Mętel^1^, Magdalena Kostrzon^2^, Justyna Adamiak^1^

^1^ Institute of Applied Sciences, University of Physical Education in Krakow, Poland

^2^ “Wieliczka' Salt Mine Health Resort, Wieliczka, Poland

Aim: Evaluation of the influence of pulmonary rehabilitation conducted in the underground salt chambers on the chest mobility of the elderly. Methods: The study was conducted between March 2019 and December 2019 and included 51 individuals 65 years of age and older with chronic respiratory conditions. The patients underwent the chest mobility test with the measuring tape before and after an outpatient pulmonary rehabilitation conducted 135 meters below the Earth's surface, for a period of 3 weeks (6 hours a day, for 5 days a week) in the “Wieliczka” Salt Mine Health Resort. Results: The test group for the eventual trial included 44 patients with mean age 68.8±2.9 years, mean BMI 28.5±3.6: 28 women and 17 men with the mean age 68.5±3.2 years and 69.4±2.5 years and mean BMI 28.4±3.8 and 28.6±3.5, respectively. Conditions taken as an indication for pulmonary rehabilitation combined with speleotherapy included chronic diseases of the lower respiratory tract (59%, 26 patients) such as: bronchial asthma, COPD, and bronchiectasis, as well as chronic diseases of the upper respiratory tract (41%, 18 patients) such as: sinusitis, pharyngitis, and laryngitis. The average chest circumference difference between the maximum inhalation and exhalation assessed in a free-standing position increased significantly (p≤0,05) from 4.5±5.5cm before the stay to 5.4±2.8cm after the stay. Both for patients with lower and upper respiratory tract disorders the average increase of chest mobility was 0,9 cm (p≤0,05).
Conclusions: Speleotherapy combined with pulmonary rehabilitation increases the chest mobility of the examined older adults as measured by the chest mobility test and should be considered important for the elderly with chronic respiratory diseases to promote healthy aging.
Keywords: Chest mobility, pulmonary rehabilitation, elderly people, speleotherapy.

#### Motor performance and functional fitness

###### Kaisa Koivunen: Birth cohort differences in maximal physical performance in 75- and 80-year-old men and women: A comparison of two cohorts over 28 years

Authors: Kaisa Koivunen^1^, Elina Sillanpää^1^, Matti Munukka^1^, Erja Portegijs^1^, Taina Rantanen^1^

^1^ Faculty of Sport and Health Sciences and Gerontology Research Center, University of Jyväskylä, Finland

Introduction: Whether increased life expectancy is accompanied by increased functional capacity in older people at specific ages is unclear. Performance-based measurements reflect functional age better than self-assessments. We compared similar validated measures of maximal physical performance in two population-based older cohorts born and assessed 28 years apart. Material and methods: Participants in the first cohort were born in 1910 and 1914 and were assessed in 1989-1990 (Evergreen project, n=500). Participants in the second cohort were born in 1938 or 1939 and 1942 or 1943 and were assessed in 2017-2018 (Evergreen II, n=726). In both cohorts, participants were assessed at age 75 and 80 years and were recruited from the Finnish population register. All community-dwelling persons in the target area were eligible. Both cohorts were interviewed at home and examined at the research center with identical protocols. Maximal walking speed, maximal isometric grip and knee extension strength, forced vital capacity (FVC), forced expiratory volume in 1 second (FEV1) and peak expiratory flow (PEF) were assessed. Results: Walking speed was on average 0.2-0.4 m/s faster in the later than earlier cohort, depending on age group and sex. In grip strength, the improvements were 11-33%, and in knee extension strength 20-47%. In FVC, the improvements were 14-21% and in FEV1 0-14%. Cohort differences in PEF were not observed. Conclusions: The later cohort showed markedly and meaningfully higher results in the maximal functional capacity tests, which suggest that increased life expectancy is accompanied by an increased number of years lived with good functional ability.

###### Magdalena Majer: Changes in functional fitness and somatic indicators of women participating in the “Active and Healthy Senior” project

Author: Magdalena Majer^1^, Maria Gacek^1^, Grażyna Kosiba^1^, Joanna Gradek^1^

^1^ University of Physical Education in Krakow, Poland

Introduction: The project is carried out at the University of Physical Education in Cracow (POWR.03.01.00-00-T225/18), under the operational program Knowledge Education Development 2014-2020. The aim of the program is to improve the psychophysical fitness of elderly people through participation in recreational activities (health training) and lectures on the determinants of the aging process and delaying involutional changes. Health training is carried out in 4 kinds of physical activity: Nordic walking, smovey®, psychophysical exercises (BodyArt) and aqua aerobics. The program lasts 9 months, during which each person participates in 3 selected activities (3 modules of 3 months). Exercises take place twice a week, and a single training session lasts 90 minutes. Aim of the research: The aim of the research was to evaluate changes in selected fitness and somatic indicators among women participating in the project "Active and Healthy Senior" after completing a 3-month health training module. Material and methods: The study was conducted among 200 women aged 53-85 years (66.54 ± 5.41). Body composition was assessed by electrical bioimpedance and physical fitness by the Fullerton Functional Fitness Test. The statistical analysis of the results was carried out using the Mann-Whitney U test, at the significance level p <0.05. Results: The research showed significant changes in the values of functional fitness indicators in the following areas: muscle strength (p <0.001), upper limb mobility (p <0.001), pelvic and lower limb muscle strength (p <0.001), lower body flexibility (p <0.001) ) and strength (p <0.001). There were no differences in agility and dynamic balance (p = 0.112). Among the analysed somatic indices, changes were found only in the body mass and lean mass of the left arm, but they did not reach the level of statistical significance (p = 0.077). Conclusions: Participation in the 3-month health training module had a positive effect on changes in some indicators of the functional fitness of senior women.

###### Nikola Stračárová: Influence of physical activity on reaction rate in a selected group of seniors

Authors: Nikola Stračárová^1^, Lenka Svobodová^1^, Martin Sebera^1^, Alena Skotáková^1^

^1^ Faculty of Sports Studies, Brno, Czech Republic

We live in modern times, people live longer, medical care is improving, while fewer children are being born, the population is aging. Life expectancy often does not mean prolonging life to the same quality as young people. Seniors face many disabilities, they are not as fast and agile, and life can be more difficult for them. Public transport, various automated call, and reservation systems in offices, self-service cash registers, can be for seniors whose cognitive abilities are limited and the reaction time extended, made difficult or outright impossible. If we improve the speed of response in the aging population, we can involve the aging part of the population more in full life, improve their independence and at the same time slow down the process of mental aging. Aim: The aim of this research was to determine the effect of three different types of training on selective response rate and motor response rate in the senior population. Methods: We compare the effect of three different types of training, namely resistance training - included 3 sets of 8 to 10 repetition at 75% of one-repetition maximum focused on the large muscle groups, balance training - included 4 sets of 8 to 10 repetition, on the bossu or with gym ball and Combination training - once a week resistance training, once a week balance training. We conducted a nine-week intervention training program. Twice a week, 60 min, 9 weeks. The study includes seniors aged 61 to 82 with an average age of 69.9 years, who were divided into three groups according to the type of training resistant, balanced, and combined. The number of participants was 71, of them 8 men and 63 women (n = 71; 63 women and 8 men). The diagnostics of the monitored parameters took place before the start of the intervention program and after its end. It was measured by the VTS test-Vienna test system (Schuhfried GmbH) test to determine the reaction time and motor reaction rate. It measures the selective response to a visual and acoustic stimulus. It is a widely used system with its own standards, which are not standardised. The research performed an analysis based on raw data on the average reaction time, the average motor reaction time, the number of correct, erroneous, incomplete responses, and the number of reactions that did not occur. These data make it possible to evaluate the concentration of attention, deficits of attention, reaction rate, and motor speed examined in a selective reaction process. Processing of results: determination of materiality and presentation of coefficients “effect size” and Cohen’s D. Results: We know from previous research that there is an improvement. More detailed results of our research will be published at the conference.

###### Agnieszka Kowalska: Evaluation of the impact of an exercise program implemented in an outdoor gym on the physical fitness people over 60 years old

Author: Agnieszka Kowalska^1^

^1^ University School of Physical Education in Kraków, Poland

Aim: The aim of the study was to assess the impact of four-week training in an outdoor gym on the physical fitness of people in the age 60-74, measured using Fullerton Test. Material and methods. The study material was a group of 31 people, of which 26 people (18 women and 8 men) completed the study (x̅=66,8± 4). Before and after training in outdoor gym subjects subjectively assessed their physical fitness, muscle strength, flexibility, endurance and well-being, using proprietary rating scale and also performed Fulleron Test. Training in outdoor gym took place 3 times a week, in 45-minute session for 4 weeks. Results: The four-weeks training in outdoor gym allowed for a statistically significant improvement in subjective assessment of physical fitness, muscle strength, flexibility, endurance and well-being. All the parameters assessed in the Fullerton Test improved statistically and the value of the BMI index decreased. Conclusions: Training in the outdoor gym has a significant impact of the fitness components of Fullerton Test, decreased the value of BMI and subjective assessment of health and well-being.

###### Yael Netz: Prescribing individualised exercise programs based upon remote assessments of motor fitness: A pilot study among healthy people aged 65 and over

Authors: Eti Benmoha^1^, Jeremy M Jacobs^2^, Ziv Yekutieli^3^, Michal Arnon^1^, Esther Argov^1^, Keren Tchelet^3^, Yael Netz^1^

^1^ The Academic College at Wingate, Israel

^2^ Hebrew University of Jerusalem, Jerusalem, Israel

^3^ Montfort Brain Monitor LTD, Israel

Background: Recent updated guidelines for physical activity emphasize multiple fitness components among people aged >65. The age-related increase in variability of fitness necessitates an accurate individualised assessment, prior to the optimal prescription of personalised exercise programs. Accordingly, we developed a novel tool to remotely assess balance, flexibility, and strength using smartphone sensors (accelerometer/gyroscope), and subsequently deliver personalised exercise programs via smartphone. Methods: We enrolled 52 healthy volunteers (34 females) aged 65+, with normal cognition and low fall risk. Baseline data from remote smartphone fitness assessment were analysed to generate 42 fitness digital markers, subsequently used to guide personalized exercise programs (five times/week for six weeks) delivered via smartphone. Programs included graded exercises for upper/lower body, flexibility, strength, and balance (dynamic, static, vestibular). Participants were retested after six weeks. Results: Average age was 74.7±6.4 years; adherence was 3.6±1.7 exercise sessions/week. Significant improvement for pre/post testing was observed for 10/12 digital markers of strength/flexibility for upper/lower body (Sit-to-Stand repetitions/duration; arm-lift duration; torso-rotation; arm-extension/flexion). Balance improved significantly for 6/10 measures of tandem-stance, with consistent (non-significant) trends observed across 20 digital balance markers of tandem-walk and one leg-stance. Balance tended to improve among the 37 participants exercising ≥3/week. Conclusion: The high adherence and improved fitness serve as proof of concept, confirming the benefits of remote fitness assessment for guiding home personalised exercise programs among healthy people aged > 65. Future research is necessary to determine the potential benefits for specific patient groups, such as those with frailty, deconditioning, cognitive and functional impairment.

#### Movement, activities and lifestyles

###### Veronique Wolter: Care providers and sports clubs? Chances and barriers of joint activities for older people in stationary and ambulant settings

Author: Veronique Wolter^1^, Miriam Dohle^1^, Lisa Sobo^1^, Cara Stemski^1^

^1^ TU Dortmund University, Germany

Physical activity in a group brings high and long-term added value for the participants. Especially for older people in need of care this development is dependent on interdisciplinary thinking and the networking of local structures. Studies underline the consideration of the communication and access options that are needed to be able to promote the target group´s health through exercise offers. Interestingly, local sports clubs are repeatedly mentioned in national and international publications as competent district partners, but in municipal practice - possibly due to very different basic structures - less thought is given than would be sensible. Particularly in the field of care, outpatient and inpatient, there have so far been more private initiatives or short-term sponsored collaborations with sports clubs. The project “Moving Nursing Homes and Care Providers” is coordinated by the State Sports Federation of North Rhine-Westphalia, Germany. Local sports clubs cooperate with outpatient and inpatient care providers and start new sports programs for older people in need of care. As part of the scientific evaluation (2020-2022), the perspectives involved - care providers, local sports clubs and (non-)participants - are equally considered and their motives and needs are analysed. This talk presents the first results from the qualitative interviews with the organisational level about promoting the opportunities of social participation of older people in their different living environments through sports groups. Chances and barriers to the organisation and sustainable design of a cooperation between the big players “care” and “organised sport” on the local level will be discussed.

###### Manca Peskar: Age-related slowing of hand and food reaction times

Authors: Manca Peskar^1^, Uroš Marušič^1^

^1^ Science and Research Centre Koper, Slovenia

Aging is associated with a higher risk of falls and behavioural slowing in tasks requiring speeded responses. Nevertheless, rapid movements in response to environmental perturbations might be crucial for avoiding dangers, such as stepping over an unexpected obstacle or rebalancing after a brief incident of balance loss. Therefore, the current pilot study investigated age-related differences in hand and foot response times of twenty healthy individuals. Ten young adults (mean age = 32 years) and ten older community-dwelling adults (mean age = 69 years) participated in neurophysiological measurements. The psychomotor vigilance task was administered during electroencephalographic (EEG) recordings to extract stimulus- and response-locked event-related potentials (s-ERP and r-ERP). Our preliminary results show that hand and foot response times were significantly shorter in young as compared to older adults (delta 27ms and 18ms, p<0.05, respectively). Moreover, s-ERP revealed shorter P200 latency (p<0.05) while r-ERP revealed larger peak amplitude (p<0.05) in older as compared to younger adults. The age-related differences in ERPs are elucidating brain mechanism involved in behavioural slowing of healthy aged individuals and carry the potential of being used as neural markers of deteriorating sensorimotor integration. Simultaneously, these markers might offer new possibilities to monitor and assess the efficacy of sensorimotor and cognitive training interventions aiming at promoting healthy aging.

###### Lenka Svobodová: Variables interfering into the quality of standing balance in older adults

Authors: Lenka Svobodová^1^, Martin Sebera^1^, Alena Skotáková^1^, Marta Gimunova^1^

^1^ Faculty of Sports Studies, Brno, Czech Republic

Background: The ability to maintain a standing balance influences the risk of falling daily living activities. As age increases, neuro-musculoskeletal and sensory changes may lead to impaired postural control and difficulties in maintaining balance. Objectives: This paper provides an overview of the variables interfering into the quality of the standing balance, one of the limiting risk factors of falling. Participants: One hundred four independent older adults (23 men with the mean age of 73.0; 80 women with the mean age of 74.5). Main measures: Standing balance was measured during a narrow bipedal stance on the platform with closed eyes (Zebris FDM, Medical GmbH). Types of falls, sensory deficit, and physical fitness were assessed using a questionnaire. For finding the best predictors, we used the data mining statistical method C&RT (Classification and Regression Tree). This method is used to classify cases based on a set of predictor variables. Unlike linear or nonlinear regression-like algorithms, this module will find hierarchical decision rules to provide the optimal separation between observations regarding a categorical or continuous criterion variable. Results: Among the examined predictor variables, narrow bipedal stance with closed eyes is influenced by the following variables ordered by the importance: falls with treatment, falls with hospitalisation, sensory deficit, and physical fitness. Conclusion: Age-related deficits in the standing balance seem to be essentially responsible. Further identification of variables input into the quality of the standing balance is suitable for designing a program to develop balance abilities and practice compensation strategies.

###### Stephanie Schmidle^:^ Insights from monitoring activities of daily living in elderlies with a smartwatch

Author: Stephanie Schmidle^1^, Philipp Gulde^2^, Joachim Hermsdörfer^1^

^1^ Human Movement Science, Technical University of Munich, Germany

^2^ Center for Clinical Neuroplasticity Medical Park Loipl, Bischofswiesen, Germany

Frailty is accompanied by limitations in activities of daily living (ADL). These are associated with reduced quality of life, institutionalisation and higher health care costs. Long-term monitoring ADL could allow creating effective interventions and thus reduce the occurrence of adverse health outcomes. The main objective of this study was to evaluate if ADL task performance can be assessed by a smartwatch’s accelerometer, and whether these measures can differentiate individual’s frailty. ADL data was obtained from twenty-seven elderly who performed two ADL tasks. Acceleration data of the dominant hand was collected using a smartwatch. Participants were split up in three groups, F (frail, n = 6), P (pre-frail, n = 13) and R (robust, n = 8) retrospectively. Measures were calculated from the vector product: Trial duration (TD), relative activity (RA), peak standard deviation (STD), peaks per second (PPS), peaks ratio (RATIO), acceleration per second (AccS), weighted sum of acceleration per second (SUM), signal to noise ratio (S2N), mean peak acceleration (MPA) and the 95th percentile of acceleration peaks (Max95). STD, PPS, SUM and Max95 showed good reliability over both tasks (r = 0.44 - 0.69). Three parameters (STD, PPS, MPA) revealed significant results differentiating between groups (effect sizes 1.30–1.70). Multiple linear regression showed that only STD significantly correlated with the Fried score (R2 = 0.25). The results demonstrate that ADL task performance can be assessed by smartwatch-based measures and further allows drawing conclusions on the frailty status of elderly, although the predictability of the exact Fried score was limited.

###### Ellen Bentlage: Practical recommendations for maintaining active lifestyle during the COVID-19 pandemic: A systematic literature review

Authors: Ellen Bentlage^1^, Achraf Ammar^2^, Daniella How^1^, Mona Ahmed^1^, Khaled Trabelsi^3,4^, Hamdi Chtourou^3,5^, Michael Brach^1^

^1^ Institute of Sport and Exercise Sciences, University of Münster, Germany

^2^ Institute of Sport Sciences, Otto-von-Guericke University, Magdeburg, Germany

^3^ High Institute of Sport and Physical Education of Sfax, University of Sfax, Tunisia

^4^ Research Laboratory: Education, Motricité, Sport et Santé, Sfax, Tunisia

^5^ Research Unit: Physical Activity, Sport, and Health, Tunis, Tunisia

Dimished volumes of habitual physical activity and increased sedentary levels have been observed as a result of COVID-19 home confinement. Consequences of inactivity, including a higher mortality rate and a poorer general health and fitness, have been reported. This systematic review aimed to provide practical recommendations for maintaining active lifestyle during pandemics. In May 2020, two electronic databases were used to search for relevant studies. A total of 1206 records were screened by two researchers. Thirty-one relevant studies were included. This systematic review revealed that reduced physical activity levels are of serious concern during home confinement in pandemic times. The recommendations provided by many international organisations to maintain active lifestyles during these times mainly target the general population, with less consideration for vulnerable populations (e.g., older adults, people with health issues). Therefore, personalised and supervised physical activity programs are urgently needed, with the option to group-play physical activity programs (e.g., exergames). These can be assisted, delivered, and disseminated worldwide through information and communication technology solutions. If it is permitted and safe, being active outside in daylight is advised, with an effort level of mild to moderate using the rating of perceived exertion scale. Relaxation techniques should be integrated into the daily routine to reduce stress levels. On the evidence base and levels of the included articles in this review, the results need to be interpreted with caution. Given that policies are different across regions and countries, further research is needed to categorize recommendations according to different social-distancing scenarios.

### POSTER PRESENTATIONS

#### Poster Session 1

###### Ivan Serbetar: The influence of fitness training program on resistance, flexibility and body fat mass of middle-aged women

Authors: Ivana Nikolic^1^, Ivan Serbetar^1^

^1^ Faculty of Teacher Education, University of Zagreb, Croatia

Adult age typically involves an overall decline in motor abilities of women. In addition to reduced mobility and functional limitations, health-related threats like osteoporosis, sarcopenia and increased risk of falling are more likely to occur in later aging. Exercise program was carried out with the aim to improve the arm strength, upper body strength and overall flexibility. The participants were 31 females aged 39-63 years (Mean=47.5; SD=6.19). A twelve-week cardio-endurance aerobic and strength program with three exercise sessions per week, each lasting 60 minutes, was conducted. The program included 35 single training sessions, each consisting of a 5-minute warm-up, 20-minute cardio endurance exercises, 25-minute resistance exercises, and 10 minutes of stretching. Pre- and post- assessment, carried out in September and January, respectively, included subscapular and abdominal skinfold measures, sit-ups, knee push-ups, and sit-and-reach test. Data were analysed using ANOVA. Statistically significant changes between the two assessments, which indicated improvement, were found in sit-up (p=.000), push-up (p=.010) and sit-and-reach tests (p=.050). Although the reduction of body fat was not observed, strength and range of motion of subjects improved significantly, which justify implemented exercise program and encourages participants to pursue lifelong healthy habits.

###### Gintarė Katkutė: Neurochemical correlates of balance stability and dual-task effects in older adults with intact cognitive functioning and mild cognitive impaired patients

Authors: Gintarė Katkutė^1^, Vida Česnaitienė ^1^, Margarita Drozdova-Statkeviciene^1^, Simona Kusleikiene^1^, Kristina Valatkeviciene^2^, Rymante Gleizniene^2^, Kazimieras Pukenas^1^, Oron Levin^1,3^, Nerijus Masiulis^1^

^1^ Lithuanian Sports University, Kaunas, Lithuania

^2^ Department of Radiology, Kaunas Clinics, Kaunas

^3^ KU Leuven, Department of Movement Sciences, Heverlee, Belgium

Aging is associated with gradual alterations in the neurochemical characteristics of the brain, which can be assessed in-vivo with proton-magnetic resonance spectroscopy (1H-MRS). However, the impact of these age-related neurochemical changes on balance control is still poorly understood. Here, we address this knowledge gap by examining the associations between neurochemical integrity of the aging brain and posturographic measures of balance stability obtained during single (ST) and dual-tasks (DT) in older adults (age-range 60-85y). 1H-MRS date were collected form voxels in the left hippocampus (LHC), left medial temporal cortex (LMTC), posterior cingulate cortex (PCC), left sensorimotor cortex (LSM1), and right dorsolateral prefrontal cortex (RDLPFC). Neurometabolites of interest were Nacetylaspartate (NAA), glutamate-glutamine complex (Glx), choline (Cho) and myoinositol (mI) that were normalised to the level of creatine (Cr) from each voxel. Balance stability was estimated through the calculation of the center-of-pressure (CoP) velocity (Vcop) and dual task cost (DTC) was calculated as a percentage of change in Vcop from single to dual task conditions. The Montreal Cognitive Assessment (MoCA) test was used to evaluate the cognitive status of all participants. Observations revealed that higher Cho/Cr ratio in the PCC, higher mIns/Cr in the LHC and lower NAA/mIns in the LHC were related to high DTC in older individuals that were diagnosed with mild cognitive impairment (MCI) but not in older individuals with apparently intact cognitive functioning (MoCA ≥ 26). These findings highlight the role for 1H-MRS to study neurochemical correlates of balance control deficits in MCI.

###### Rodrigo Gallardo: Sedentarism, mental health and presence of pathologies on Chilean older adults during confinement due to COVID-19 pandemic

Authors: Rodrigo Gallardo^1^, Orlando Gallardo^2^, Karen Gallardo^3^

^1^ Departamento de Ciencias del Deporte y Acondicionamiento Físico, Universidad Católica de la Santísima Concepción – Concepción, Chile

^2^ Departamento de Educación Física, Facultad de Educación, Universidad de Concepción – Concepción, Chile

^3^ Escuela de Educación Física, Facultad de Educación, Universidad de las Américas - Concepción - Chile

Scientific evidence has demonstrated that sedentarism is related to the prevalence of numerous pathologies and a higher level of anxiety and depression. Confinement due to the COVID-19 pandemic has significantly impacted increasing sedentary lifestyles, especially in older adults. This study aimed to analyse the relationship between sedentary time due to confinement in Chilean older adults and anxiety and depression levels. Furthermore, this research intends to compare whether the sedentary time is higher when they suffer different types of pathologies. The sample comprised of 23 men and nine women (age=66.35±7.16). The sedentary item of the International Physical Activity Questionnaire (IPAQ-short form), an ad-hoc questionnaire to collect information about the presence of pathologies, and the Spanish version of the Hospital Anxiety and Depression Scale (HADS) were applied. Pearson coefficient showed a significant correlation between people who spent more time sitting weekly with higher levels of anxiety (p <.05) and depression (p<.001). According to the number of pathologies, a significant difference between sedentarism levels was calculated through ANOVA with posthoc Bonferroni test. Subjects who presented more than one pathology spent more time sitting weekly (p<.05). A multivariate regression analysis indicated that sitting weekly time spent by older adults can be predicted by a linear combination of the level of anxiety and depression and the presence of more than one pathology (p<.05). These findings suggest that emotional state and pathologies can be a risk factor against reducing the sedentary lifestyle. Future research should seek to gather more population under different confinement conditions.

###### Gabrielle McKee: Why older adults engage in a physical activity app: A qualitative analysis

Authors: Dr. Aileen Lynch^1^ Dr. Fintan Sheerin^1^, Dr. Sean Kilroy^1^, Dr. Gabrielle McKee^1^

^1^ School of Nursing and Midwifery, Trinity College Dublin, Ireland

Background: Approximately 30% of older adults meet international physical activity (PA) guidelines. While many apps promoting PA are available few are targeted for older adults or include substantial behavioural change techniques. Goals and planning are among the most successful behavioural change techniques used in PA apps including goal setting behaviour i.e. being able to walk 100m and goal setting outcomes i.e. feeling healthier. Aim: To identify the main reasons why older adults wanted to engage in an app to improve or maintain their PA. Method: A cross sectional, qualitative, online short survey was completed by community dwelling older adults (60+), who were able to engage in physical activity. The participants were asked a single open question to assess their goal setting outcome “Why did they engage in this PA app”. Analysis used descriptive qualitative analysis. Findings: Twenty-four participants enrolled. Responses given were categorised into five themes/ subthemes: General health (i.e. staying healthy), physical health (i.e. maintaining fitness), functional health (i.e. improving balance), mental health (i.e. relieving stress) and prevention of disease and ageing deterioration (i.e. preventing falls). Framing of the reasons using the Self Determination Framework revealed that participants’ motivation for engaging in a PA app was mainly internally driven and of personal importance. Conclusions and implications: To successfully engage older adults in PA, it must be meaningful for them. Incorporation of PA outcomes in any intervention will assist older adults and their health team to develop strategies that are more targeted, personalised, help make physical activity a habit.

###### Tal Lifshits: Physical activity level among adults –How does it change across life span and differ between men and women?

Authors: Tal Lifshits^1^, Deborah Alperovitch-Najenson^1^

^1^ Tel Aviv University, Department of Physical Therapy, Israel

**Background:** Among adults there is a strong correlation between both physical inactivity and sedentary behavior (SB) with risk factors such as mortality, chronic illness and cognitive aspects. Our aim was to review current knowledge of physical activity (PA) level among adults, and the differences between age groups and sexes. **Methods:** Databases were searched for studies of any methodological quality, published in English from 2010 until September 2020. **Results:** This review includes studies from 13 countries, taken place between the years 2000-2015. The population includes 37,687 adults aged 18-102 years old. A clear decline was found in moderate to vigorous PA (MVPA) level as age increased, in all studied populations, among men and women. In all the studies women performed less MVPA than men. Lower vigorous PA (VPA) level was found among women compared to men at the same age. Yet, in almost all the studies, women performed more low PA (LPA). A decreasing trend of LPA levels could be seen in both sexes, starting from 65-age group. SB time significantly increased above the age of 65 and was higher among men compared to women. WHO PA guidelines define the amount of weekly PA need to be performed. In the reviewed studies, adults aged 70 and above were found as not performing VPA at all. **Conclusions:** Cut points used to measure PA level are too general, and PA guidelines refer to young adults. Therefore, we suggest building specific cut-points and PA guidelines for different age groups and sexes, for more accurate research.

###### Kristīne Šneidere: Relationship between cognitive reserve, physical activity, hippocampal volume and working memory in older adults

Authors: Kristīne Šneidere^1^, Anete Šneidere-Šustiņa^1^, Ainārs Stepens^1^

^1^ Rīga Stradiņš University, Latvia

With age being the most significant predictor of mild cognitive impairment and Alzheimer’s disease, the increase in average life-span adds significant concerns regarding the on-going social and economic burden of cognitive impairment later in life. In this study, we aimed to investigate the relationship between long-term physical activity, cognitive reserve, working memory and hippocampal volume. Materials and methods 46 participants, aged 65 to 85 (M = 71.41, SD = 5.12, 23.9% male), with no self-reported on-going neurological, oncological or psychiatric diseases, were involved in the study. Long-term physical activity was assessed using Social Determinants of Health Behaviour questionnaire (FINBALT, 2008), for working memory measures, the Numbers Reversed task was used (Woodcock et al., 2001), cognitive reserve (CR) was determined using Cognitive Reserve Index questionnaire (Nucci et al., 2012) and MRI data were obtained with Siemens 1.5T. Results Hierarchical multiple regression analysis was conducted. The addition of physical activity and cognitive reserve to the prediction of working memory did not lead to a statistically significant increase in R2 and only hippocampal volume significantly predicted working memory scores (R2 = .17, F(44) = 9.02, p = .004), explaining 17% of the variance. Conclusions Greater hippocampal volume predicts better working memory functioning, with such life-style factors as physical activity and cognitive reserve showing no significant impact; however, the limitation of sample size and large age range could have an impact on the current results.

###### Saar Frank Herschkovitz: The effect of a single bout of aerobic exercise on limb motor inhibition as compared to cognitive inhibition in middle age adults

Author: Saar Frank Herschkovitz^1^, Gal Ziv ^1^, Oron Levin^2^, Yael Netz^1^

^1^ The Academic College at Wingate, Israel

2 Katholic University Leuven, Belgium

Background: Motor inhibition is known for its importance for adults in their daily living. It enables performance of selective movements as a result of a given state. Aim: To examine the effect of a single bout of exercise on motor inhibition as compared to its effect on cognitive inhibition among middle-aged adults. The motor inhibition included upper and lower limbs coordination. Methods: 36 healthy subjects (21 women, 15 men) aged 40-60 years old were randomly assigned by gender to either experimental or control group. Each subject was examined in two sessions. The first session included baseline tests of motor inhibition (all limbs coordination) and cognitive inhibition (verbal Stroop test). The second session included a shorter version of the motor inhibition test (three limbs coordination) and the cognitive inhibition test. Subjects in the experimental group performed 26 minutes of moderate intensity aerobic exercise (60% Heart Rate Reserve) on a treadmill whereas the control group, remained sedentary in a chair (resting heart rate). Immediately after, all subjects performed the two tests again (post-tests). Results: Two-way (groups X time) ANOVAs with repeated measures are applied to examine differences between baseline and pre-test scores, and between pre and post test scores on each of the cognitive and motor inhibition measurements. Pearson correlation analysis is used to examine the relation between both forms of inhibition, at baseline tests, pre and post-tests. The preliminary results show no effect of exercise. Further analyses are currently conducted to examine whether there are differences between responders and non-responders.

###### Laimute Samsoniene^:^ The diversity of successful aging: The empowerment of sports veterans

Author: Laimute Samsoniene^1^

^1^ Vilnius University, Institute of Health Sciences, Faculty of Medicine, Lithuania

The International Association of Sports Veterans organises the World Sports Veterans Games (WMG) every four years, integrating more than 30 activities or games. WMG's operating philosophy: to pass on the general and sports culture and experience of different countries to future generations; monitor trends in healthy ageing and contribute to a healthier society. Lithuanian Sports Veterans Association "Penki žiedai" has been participating in WMG activities since 1994, and since 2008. and the European Veterans Games. Methodological provision: the phenomenon of empowerment can be revealed through two interrelated groups of components: the group of general empowerment components and the group of personal empowerment components. Organisation of the study: Components of collective empowerment. A systematic analysis of Lithuanian normative documents, selection criteria: functioning normative documents, search words: sports, health, physical activity, sports veterans; sports. Components of personal empowerment. Research methods: interview method; data is summarised by analysing qualitative content. Fifteen sports veterans were randomly selected: 10 men and 5 women. Sampling criteria: sports experience of at least 30 years, active sports at least 2 times a week and participation in national and / or international competitions. Results: The representative sport of veterans is to be protected as a value that is fundamental, a living collective memory of sport. The modern Lithuanian sports system, equating the sports culture of veterans only with physical activity, hinders its development and damages its’ establishment in the general sports culture. Individual uncoordinated recreational programs of physical activity cannot safely and competitively prepare sports veterans for the International Sports Veterans Games. The findings of the study will contribute to diversity in addressing the challenges and problems of a successfully aging population, and, according to the study, sports experts and politicians will need to provide new guidelines for the interpretation of sports veterans at the societal and political levels. Keywords: empowerment, sports veterans, sports, physical activity.

###### Kathleen Kang: Plasticity of sequential decision making in old age

Authors: Kathleen Kang^1^ & Shu-Chen Li^1,2^

^1^ Lifespan Developmental Neuroscience, Faculty of Psychology; Technische Universität Dresden, Dresden, Germany

^2^ Centre for Tactile Internet with Human in the Loop, Technische Universität Dresden, Dresden, Germany

Older adults face difficulties in decision-making, especially in complex situations where choices need to be made based on delayed outcomes. This difficulty has been attributed to aging-related under-recruitment of the fronto-parietal networks (cf. Eppinger et al. 2015). Since decision-making is a crucial cognitive function, it is imperative to improve decision-making abilities and prevent any further decline through cognitive interventions. However, cognitive interventions to improve sequential decision-making in older adults are still scarce. Therefore, the current study aims to investigate the effectiveness of a home-based sequential decision-making intervention in older adults. Twenty-eight participants (15 females, mean age = 67.89, SD = 1.46, range = 65-70 years old) participated in this single-arm interventional trial where they were asked to do the Markov three-stage decision-making task over a period of 6 weeks (18 sessions in total). The adherence rate was 79%, with 22 participants completing all 18 sessions. In this task, participants had to learn the optimal sequence of action which will provide them with the most reward in the long run. We analysed the data using linear mixed effects model with training sessions (within-subjects) and performance group (between-subjects) as fixed effects, and subject as random effects. Generally, participants made significantly more optimal actions at the end of training (T18) compared to the start of training (T1), *F*(17, 433) = 20.87, *p*<0.001. High improvers performed significantly better than low improvers from as early as T6. While high improvers demonstrated significantly better performance at T18 as compared to T1, low improvers did not demonstrate this, *F*(17, 433) = 6.59, *p*<0.001. Therefore, the findings of this study reiterate the suggestion that cognitive interventions have the potential to induce plasticity in cognitive functioning during aging.

#### Poster Session 2

###### Małgorzata Bagińska: Nutritional behaviour and selected lifestyle aspects in patients of a physiotherapy clinic in Ustrzyki Dolne, Poland, in the context of osteoporosis prevention

Authors: Małgorzata Morawska-Tota^1^, Łukas Tota ^2^, Katarzyna Cetnar^3^

^1^University of Physical Education in Kraków, Department of Sports Medicine and Human Nutrition, Poland

^2^ University of Physical Education in Kraków, Department of Physiology and Biochemistry, Poland

^3^ University of Physical Education in Krakow, Department of Physical Education and Sport, Poland

Introduction: Osteoporosis is a major health, social, and economic problem of the contemporary world. In 2018, the disease was diagnosed in 5.5% of individuals aged over 50 years (including 4.4% of women) in Poland. Purpose: The aim of the study was to assess the nutritional habits and selected lifestyle aspects in patients of a physiotherapy clinic in Ustrzyki Dolne, Poland, in the context of osteoporosis prevention. Material and Methods: The study involved 80 patients (40 women and 40 men) of the Physiotherapy Clinic of the Independent Public Health Care Centre in Ustrzyki Dolne, Poland, aged 46–80 years. The general health of the participants was good, and the vast majority had not been diagnosed with a chronic disease or fracture within the previous 5 years. Overweight or obesity was observed in 50% of women and 70% of men. Dietary habits and selected aspects of lifestyle were assessed by using an authors’ own questionnaire. Results: The participants implemented the principles of proper nutrition to a limited extent. The qualitative evaluation of the respondents’ diet revealed numerous inadequacies, including over-consumption of meat and meat products, as well as under-consumption of fruit, vegetables, and dairy products. Supplementation was applied by about half of the participants, who mostly chose calcium (65%), fish oil and omega-3 acids (57%), and vitamin D (56%). Only 31% of the respondents declared undertaking recreational physical activity at least several times a week (minimum 3 hours per week), and 56% did not smoke. Conclusions: The participants’ dietary errors, sedentary lifestyle, and smoking may translate into an increased risk of osteoporosis in the future. In order to prevent diet-related diseases, it seems justified to provide nutritional education to seniors.

###### Zbigniew Ossowski: Association between sarcopenia related parameters and cognitive functions in postmenopausal women

Authors: Zbigniew Ossowski^1^, Mirosław Smaruj^1^, Vida Česnaitienė^2^, Wojciech Skrobot^1^, Piotr Aschenbrenner^1^

^1^ Gdansk University of Physical Education and Sport, Poland

^2^ Lithuanian Sports University, Kaunas, Lithuania

Background: Age-related loss of skeletal muscle mass and function can lead to sarcopenia. Sarcopenia is particularly common among elderly women and is associated with significant morbidity and mortality. Data on the relationship between sarcopenia and cognitive function is very limited. Therefore, the aim of the study was to determine the relationship between the Skeletal Muscle Index (SMI), strength, functional performance and cognitive functions in postmenopausal women. Methods: The study involved 84 women between 61 and 81 years of age. The Handgrip Muscle Strength (HS), Gait Speed (GS) and SMI were used to measure sarcopenia. Moreover, cognitive function was assessed using Trail Making Test – B version (TMT-B). Results: The SMI and GS was positively correlated with the cognitive function in women. However, no significant relationship was found between HS and TMT-B in the subjects. Conclusions: The results suggest that the level of cognitive functions in postmenopausal women may have a positive relationship with their skeletal muscle mass and functional performance. However, further studies are needed to identify recommendations for programming physical activity aimed at improving cognitive function in women at risk of sarcopenia.

###### Jessica Koschate: Covid-19 pandemic – Time course of physical activity and functional abilities in active older people associated with the lockdown periods in Germany

Authors: Jessica Koschate^1^, Sandra Lau^1^, Michael Hackbarth^1^, Tania Zieschang^1^

^1^ Carl von Ossietzky University Oldenburg, Department Geriatrics, Germany

Introduction: The purpose of the study was to analyse physical activity of older adults during the COVID-19 lockdown periods (LDP1, LDP2) and monitor functional performance. Methods: 35 older adults (20 females, 71±6 yrs, 24±8 kg·m-2), exercising in a chip-controlled fitness circuit were interviewed about their daily physical activity habits (modified Minnesota Leisure Time PAQ), functional ability (LUCAS robust score [RS]) before, during and after the LDP1 as well as before and during LDP2. Data on individual leg strength were extracted from the chip-controlled circuits before and after LDP1. Results: Energy expenditure (EEXP) of daily activities during LDP1 were lower compared with post LDP1 (4230±2775 kcal/wk; p=0.001) and pre LDP2 (5230±2822 kcal/wk, p<0.001), EEXP before LDP1 was lower (3117±1955 kcal/wk) compared with pre LDP2 (p<0.001), EEXP at pre LDP2 was higher than during LDP2 (4113±2469 kcal/wk, p<0.001).The RS decreased during the two LDPs (pre LDP1: 5±1, LDP1: 4±1, post LDP1: 5±1, LDP2: 4±1, p<0.001). Especially exhausting activities were reduced during the LDPs (pre LDP1: 91.4%, LDP1: 20%, post LDP1: 80%, LDP2: 8.6%). The RS during LDP1 correlated significantly (rSP=-0.350; p=0.029) with the decrease in the leg score. Discussion: Physical activity of active older people was reduced during LDP2. Factors of robustness decreased during the LDPs, which was mainly caused by less intense exercise. Lower RS during LDP1 were associated with decreases in leg strength after the LDP. Apparently, intensive exercise was not compensated when access to the fitness circuits was lacking. Adequate home-based alternatives should be promoted during future lockdowns.

###### Katarzyna Kucia: The effects of 3-month aqua-fitness program on the functional fitness in elderly women

Authors: Katarzyna Kucia^1^, Joanna Gradek^1^, Grażyna Kosiba^1^, Magdalena Majer^1^, Maria Gacek^1^

^1^ University of Physical Education in Krakow, Poland

Purpose: The main aim of the study was to assess the functional efficiency fitness impact of the motor control Fullerton Senior Fitness Test, before and after the application 3 – months Aqua Fitness program among women over 60. Participants: The study involved 28 seniors qualified for the Academy of Physical Education in Krakow "Healthy and Active Senior" program in the academic year 2019/2020. The project involved selecting participants for the Aqua fitness double-blind randomized trial (n=28, 68±9 years, body mass index BMI 27.7±4. Methods:
Selected group followed a supervised training routine 2 times/week for 3 months. The variables assessed at baseline and after 3 months were: body composition (BIA), anthropometric indexes i.e. Body Mass Index and Senior Fitness Test (FFST) which evaluated functional fitness. Measured fitness parameters was: strength and muscular endurance, mobility, dexterity, speed, body balance, motor coordination, reaction time, flexibility Results: After 3-months the greatest changes were found in both endurance, agility and strength of the upper and lower limbs. Also were noticed optimistic changes in increases leg strength, and flexibility lower and upper body part. After 3 months Body Mass Index and Fat Mass decreased significantly (P<0.05), Basal Metabolic Rate, Fat Free Mas and Total Body Water definitely increased (P<0.001) Conclusion: Our findings indicated that Aqua Fitness can give some appropriate health parameters better and quicker than other disciplines. It is probably related to the aquatic environment specificity (buoyancy, density, resistance force or wave) and the physical effort in the water.

###### Aileen Lynch: A pilot study to assess the effectiveness of a brief intervention on community dwelling older adults’ physical activity

Authors: Gabrielle McKee^1^, Fintan Sheerin^1^, Sean Kilroy^1^, Mary Harkin^2^, Ciarán Mckinney^2^, Aileen Lynch^1^

^1^ School of Nursing and Midwifery, Trinity College Dublin, Ireland
^2^ Age and Opportunity, Ireland

Background: Physical activity (PA) is one of the target risk factors when assisting people make healthier lifestyle options. As only 30% of older adults meet the recommended PA guidelines, effective interventions are required to improve their engagement in PA. Aim: To evaluate the effectiveness of a brief intervention in improving PA in older adults. Method: In this quasi experimental study, community dwelling older adults (age 50+) attending age-related public fairs were invited to partake in a brief intervention (5-15min) to assist them in improving or maintaining their PA. The brief intervention, presented by trained individuals, addressed the following: participants’ facilitators and barriers to PA engagement; their preferred activities; their PA goals and specific planning around achieving these goals (what activity, when, with whom, an alternative plan if required). Participants were health screened for their suitability to partake in PA (PAR-Q) and their PA was assessed at baseline and at 3 months using the IPAQ. Ethical approval was granted for this study. Results: At baseline, 27 participants enrolled in the study. Of the participants who partook in the follow-up PA assessment, their PA levels increased from 57% at baseline, to 86%. Conclusions and implications: Continued engagement in PA in older age will help delay the onset of chronic disease and prolong independence. The intervention, specifically designed for older adults, was effective in improving older adults’ PA and used behavioural change principles and motivational interviewing; strategies that could be easily incorporated into many health care providers’ interactions with this cohort.

###### Gina Krause: Physical activity and exercising benefits in dementia care

Authors: Gina Krause^1^, Yael Netz^2^, Michael Brach^1^,

^1^ Institute of Sport and Exercise Sciences, University of Münster, Germany
^2^ The Academic College at Wingate, Israel

Actimentia is an Erasmus+ project aiming to foster physical activity in the everyday life of people with mild cognitive impairment or dementia and their caregivers. Dementia is a major cause of disability and dependency among elderly worldwide, and often underlies social exclusion. It has a significant impact not only on individuals but also on their caregivers, families, and communities. Dementia is a disease that gradually develops. However, an appropriate exercise program can improve the condition of dementia patients and may also improve the patients' quality of life and delay dementia progression. That’s where the Actimentia Project steps in! The Actimentia online training platform is designed to develop basic skills on physical exercising and activity for formal and informal caregivers, so that they can use them regularly during their care giving tasks for maintaining the wellbeing of the dementia patients, as well as for themselves. In addition to general background and information, the platform includes three main parts: dancing, exergames and physical activity lessons. These three areas offer various exercises at different levels of difficulty, which can be combined and selected according to preference. The online training platform is available at: [www.actimentia.org](http://www.actimentia.org). The project website is available at: [www.actimentia.eu](http://www.actimentia.eu).

###### Ana Maria Rizescu: Means of injury prevention and the influence of the correlation between the type of temperament and the severity of injuries in the game of football-tennis

Authors: Ana Maria Rizescu^1^, Mariana Cordun^1^

^1^ National University of Physical Education and Sports from Bucharest

The game of football-tennis appeared internationally, in 1922 in the Czech Republic, which was then called football over the rope, and in 1940 the first written rules appeared. Football-tennis is a relatively young sport, which grew in the ‘80s. In a football-tennis match, the technique of hitting the ball differs from that applied in football, as well as that used in tennis. The name of the football-tennis discipline together with its theoretical and practical substantiation is independent of the tennis and football sports branches. Safety in sports and physical activity are important prerequisites for continuous participation in sports, as well as for maintaining a healthy lifestyle. For this reason, the prevention and control of sports injuries are important goals for society as a whole. This article provides an overview of the correlation between the type of temperament of athletes and the severity of trauma. The temperament that influences the injury mechanisms is not limited to athletes. In addition, the components of temperaments, extroverted and introverted often act together. Some components can directly affect the risk of injury and are by definition a risk factor. Other components of extroverted and introverted temperaments may influence the risk of injury indirectly. The means of preventing sports injuries are part of what is called the “prevention sequence”. The extent of the problem of sports injuries is often described by the incidence of injuries and indicators of the severity of injuries. The main means of preventing trauma is movement in various active and / or passive forms, which by repetition becomes physical exercise. Exercise has an important role in influencing intellectual and emotional functions and contributes to the development of character and personality. In this paper we want to report that through the game of football-tennis we can contribute in terms of improving physical condition, but also the prevention of injuries through stretching exercises, mobility, elasticity and dynamic exercises, adapted to the level of the work group, from the youngest to the oldest. Keywords: football-tennis, injuries, stretching, performance, exercises.

###### Daniele Magistro: Effectiveness of a lifestyle intervention in promoting the mobility functioning in institutionalised older adults

Authors: Daniele Magistro^1^, Fabio Carlevaro^2^, Francesca Magno^2^, Martina Simon^2^, Nicola Camp^1^, Giovanni Musella^2^

1 Nottingham Trent University, United Kingdom

2 University of Torino, Italy

Institutionalised older people are at higher risk of physical functioning decline. Lifestyle interventions offer potential for reducing such negative outcomes. The present study aims at investigating the effect of a lifestyle-based physical activity intervention on mobility functioning in institutionalized older adults. A randomized controlled trial was conducted comparing a physical activity intervention and a no-treatment control condition over a 6-month experimental phase. Eighty-one individuals living in residential care facilities were recruited, with a mean age of 78 (SD = 13,2) in the intervention group and 81.9 (SD = 10.3) in the control group. Baseline and 6-months measurements for the Time Up and Go test, 30-Second Chair Stand test and Tinetti test were collected. The findings showed that the lifestyle intervention had a positive effect on physical functions. There was a statistically significant change between the means of the two groups over time for the Time Up and Go test (p < 0.05) and the Tinetti Test (p < 0.05). In general, the intervention groups showed a stable condition with respect to overall mobility function, balance, and gait while the control group showed decreased performance. These results underline that even in critical conditions, relatively simple lifestyle-based physical activity intervention may promote a more positive adjustment to old age.

###### Paulina Łukaszek: Perception of physical self-attractiveness in terms of pro-health behavior including physical activity and the use of beauty treatments in women over 60

Authors: Monika Wilk-Głodzik^1^, Paulina Łukaszek^1^, Jacek Głodzik^1^

^1^ University of Physical Education in Krakow, Poland

Many scientific reports show the relationship between pro-health behavior, especially physical exercise, and maintaining health, high physical fitness and a positive perception of one's own body. Physical activity has been indicated as the direction of health promotion in the elderly in order to maintain functional fitness and to prevent and combat various diseases. Social changes and the development of technology in the field of cosmetology and aesthetic medicine in recent years have also resulted in an increased interest in outer appearance. This phenomenon also applies to older women. It seems that the body image related to the above-mentioned factors is necessary to maintain life satisfaction at a high level. The aim of the study was to examine the perception of physical self-attractiveness in terms of various health behaviors, taking into account physical activity and the use of beauty treatments in 143 women over 60 from the urban environment. A group of women who regularly attended beauty salons was selected among the respondents. The restrictions on physical activity and the beauty industry related to the Coronavirus pandemic were also taken into account. To achieve the objectives of the study, an anonymous questionnaire was carried out, taking into account the authors’ questions and IPAQ. The use of methods to improve the appearance of the body is a complex problem and depends on many factors, including the prohibitions related to the Coronavirus pandemic. The surveyed women take care to improve their appearance to a different extent and use various methods of improving it.

###### Charlotte Sylvie Le Mouel: Postural adjustments in anticipation of predictable perturbations allow elderly fallers to achieve a balance recovery performance equivalent to elderly non-fallers

Authors: Charlotte Sylvie Le Mouel^1^, Romain Tisserland^2^, Thomas Robert^3^, Romain Brette^4^

^1^ Institute of Sport and Exercise Sciences, University of Münster, Germany
^2^ Université de Poitiers, France
^3^ Université de Lyon, France
^4^ Sorbonne Université, France

Background: In numerous laboratory-based perturbation experiments, differences in the balance recovery performance of elderly fallers and non-fallers are moderate or absent. This performance may be affected by the subjects adjusting their initial posture in anticipation of the perturbation. Research questions: Do elderly fallers and non-fallers adjust their posture in anticipation of externally-imposed perturbations in a laboratory setting? How does this impact their balance recovery performance? Methods: 21 elderly non-fallers, 18 age-matched elderly fallers and 11 young adults performed both a forward waist-pull perturbation task and a Choice Stepping Reaction Time (CSRT) task. Whole-body kinematics and ground reaction forces were recorded. For each group, we evaluated the balance recovery performance in the perturbation task, change in initial center of mass (CoM) position between the CSRT and the perturbation task, and the influence of initial CoM position on task performance. Results: The balance recovery performance of elderly fallers was equivalent to elderly non-fallers (p > 0.5 Kolmogorov-Smirnov test). All subject groups anticipated forward perturbations by shifting their CoM backward compared to the CSRT task (young: 2.1% of lower limb length, elderly non-fallers: 2.7%, elderly fallers: 2.2%, Hodges-Lehmann estimator, p < 0.001 Mann-Whitney U). This backward shift increases the probability of resisting the traction without taking a step. Significance: The ability to anticipate perturbations is preserved in elderly fallers and may explain their preserved balance recovery performance in laboratory-based perturbation tasks. Therefore, future fall risk prediction studies should carefully control for this postural strategy, by interleaving perturbations of different directions for example.

### PHYSICAL ACTIVITY PROJECTS FOR OLDER ADULTS

###### Sylwia Tałach-Kubas: Using the Actimentia training platform to build active and healthy habits among elderly at a risk of dementia – good practice during pandemia

Author: Sylwia Tałach-Kubas^1^

^1^Actimentia

The objective of this presentation is to share results of application of training based on the ACTIMENTIA platform as a tool for promoting physical activity & exercise among persons aged 60+ from Malopolska region (Poland). ACTIMENTIA is an interactive training platform developed in frame of an Erasmus + international project managed by the University of Muenster. The platform is based on video instructions and the structure includes three components: lessons – guided trainings, dances and exergames, offering wide possibilities of composing professional trainings for elderly –a group at risk of dementia. Although ACTIMENTIA was developed mostly with the idea of bringing physical activity and exercise into the daily routine of caregivers of dementia patients and care receivers’, the programme may have wider influence on promoting physical activity and regular training among seniors, especially in time of Covid 19 restrictions. PLinEU Association has organised 2 pilot trainings of ACTIMENTIA programme. For 10 weeks we have organised guided on-line trainings using the platform. The group of beneficiaries consisted of 10 persons aged 65 – 80, most of them with some mild cognitive impairments (MCI). The presentation will show some examples of trainings based on the ACTIMENTIA programme and present the benefits from the pilot intervention. Undoubtedly, this pilot training has improved both physical and psychological condition of all participants influencing their quality of life during pandemic restrictions. It may be recommended to other Senior Clubs and NGOs working with elderly, but also to individual 60+ persons as a measure of prevention and retardation of dementia and MCI.

###### Ismaël Brunot: The Actimentia Motion Capture System

Author: Ismaël Brunot^1^, Gina Krause^2^, Sina Hinternesch^3^, Michael Brach^2^

^1^ Interactive 4 D, Nice, France

^2^ Institute of Sport and Exercise Sciences, University of Münster, Germany
^3^ Institute for Health, Nutrition and Sport Sciences, Europa-University Flensburg, Germany

The video explains and demonstrates how motion capture is used to create virtual dance teachers. First of all, dances are created, which are specifically adapted to older adults with cognitive impairments or dementia. The movements are captured using cameras. Markers are positioned on the joints of the dancers. Computers collect the images of the markers captured by the cameras and then convert them into three dimensional movements. An animation file is created and then exported. Along with the motion capture, 3D characters (avatars) are created. A skeleton is integrated into the characters. Once the characters have a skeleton it is possible to animate them. The characters are now animated using the motion capture files. The 3D characters video is synchronized with music via editing software.

###### Joanna Gradek: Active and Healthy Senior, a project tailored to the times

Authors: Joanna Gradek^1^, Magdalena Majer^1^, Grażyna Kosiba^1^

^1^ University of Physical Education in Krakow, Poland

Aim: The aim of the project is to strengthen the pro-health and social competences of elderly people. The project is based on their participation in recreational activities with health training features and lectures on aging as a natural process, on which the individual has a real influence by modifying his lifestyle. Taking part in the project allowed seniors to systematically participate in organised physical activity and learn about a healthy way of life. Material and methods: The project consists of 2 editions, each lasting 9 months. The activity classes are divided into 3 three-month modules consisting of various types of physical activities: Nordic walking, smovey® exercises, aqua fitness and psychophysical exercises (including body art, pilates, yoga). Each participant was subjected to tests, which included anthropometric measurements, a physical fitness test and a survey regarding the lifestyle of the project participants. 136 people joined the first edition and they were divided into 8 exercise groups, participating in 1.5 hours of exercise twice a week for 3 months (48 hours in total) and in 9 lectures (27 hours in total). The type of physical activity changed after each module. Results: After 3 months of exercise, positive changes in the functional fitness of the participants were noticed, including balance, cardio-respiratory endurance, flexibility and muscle strength. Summary: An additional effect of the project is to create a Center for Health Education and Physical Activity of Seniors at the Academy of Physical Education in Krakow for the continuation of regular educational and integration activities, as well as improving the functional abilities of seniors.

Register of Persons

Abeln	60

Adamiak	63

Ahmed	9, 61, 73

Alperovitch-Najenson	79

Alves	32

Ammar	10, 43, 73

Argov	68

Arnon	57, 68

Aschenbrenner	85

Auerswald	8, 36

Baghdiguian	9, 38, 40

Bagińska	10, 84

Ballesteros	2, 7, 58

Benmoha	68

Bentlage	2, 12, 61, 73

Biolo	44

Bouça-Machado	61

Brach	2, 9, 12, 61, 62, 73, 89, 96

Branco	32

Brette	93

Brunot	6, 96

Bukowska	2, 7

Cachola	27

Camp	91

Carlevaro	91

Česnaitienė	76, 85

Cetnar	84

Chastin	9, 38, 39

Chtourou	73

Clauß	35

Cordes	6, 24, 37

Cordun	90

de Bruin	4, 6, 18

Degens	45

Dohle	69

Dransmann	62

Drozdova-Statkeviciene	76

Fernandez del Olmo	59

Frank Herschkovitz	8, 81

Fricke	6, 21, 25

Fröhlich	8, 34, 35

Gacek	65, 87

Gallardo	77

Gallardo	77

Gallardo	8, 77

Gaspar de Matos	28, 32

Gautier	9, 38, 41

Giannouli	11, 49, 50, 51

Gimunova	71

Gleizniene	76

Głodzik	92

Gomes	29, 32

Gouveia	28, 31

Gradek	2, 6, 27, 65, 87, 97

Grässler	10, 47

Guillet–Descas	38, 41

Gulde	72

Hackbarth	86

Halfpaap	10, 48

Harkin	88

Hermsdörfer	72

Hinrichs	2, 10, 11, 12, 49, 50, 53

Hinternesch	96

Hökelmann	2, 9, 10, 42, 43, 47

Hortobágyi	59

How	61, 73

Hübert	62

Ivaničová	56

Jacobs	68

Jansen	11, 49, 52

Judica	61

Kałamacka	59

Kang	8, 83

Katkutė	8, 76

Keskinen	53

Kilroy	78, 88

Knoll	43

Koivunen	10, 64

Koschate	11, 86

Kosiba	65, 87, 97

Kostrzon	63

Kowalska	10, 67

Krause	11, 89, 96

Krčmár	56

Krčmárová	7, 56

Kruse	25

Kucia	9, 11, 87

Kusleikiene	76

Kvedaras	55

Labott	48

Lau	86

Le Mouel	11, 93

Levin	2, 7, 55, 57, 76, 81

Li	83

Lidor	57

Lifshits	8, 79

Loureiro	27, 28, 29, 30, 31, 32

Loureiro	7, 26, 27, 28, 29, 30, 31, 32

Łukaszek	92

Lynch	11, 78, 88

Magistro	11, 91

Magno	91

Mahoney	22

Majer	2, 10, 65, 87, 97

Manet	9, 38, 40

Marchant	9, 38, 41

Marconcin	28, 29, 30, 31

Marin	61

Marques	7, 27, 28, 29, 30, 31, 32

Martins	28, 30, 31

Marušič	6, 10, 22, 46, 70

Masiulis	55, 76

McKee	8, 78, 88

Mckinney	88

Mechling	2

Mętel	2, 9, 63

Meyer	36

Morawska-Tota	84

Muiños	58

Müller	8, 33, 34, 35

Müller	43

Müller	43

Munukka	64

Murawska-Ciałowicz	30

Musella	91

Narici	44

Netz	2, 6, 7, 10, 57, 68, 81, 89

Niederer	23

Nikolic	75

Nottelmann	62

Ossowski	11, 85

Peralta	7, 28, 30, 31

Peskar	12, 70

Pišot	10, 44, 45, 46

Portegijs	11, 49, 53, 64

Pukenas	76

Rantanen	12, 19, 53, 64

Reales	58

Reggiani	44

Rehfeld	43

Rieker	58

Rittweger	45

Rizescu	11, 90

Robert	93

Roffler	50

Rosa	7, 27

Rössler	50

Rudisch	24, 37

Rukun-Halaby	57

Saajanaho	53

Sabino	7, 29

Samsoniene	8, 82

SantosNone	31

Schmidle	12, 72

Sebera	66, 71

Seinsche	11, 49, 51

Serbetar	8, 75

Sheerin	78, 88

Sillanpää	64

Simon	91

Šimunič	10, 44, 45

Skotáková	66, 71

Skrobot	85

Smaruj	85

Šneidere	8, 80

Šneidere-Šustiņa	80

Sobo	69

Stemplewski	2, 12

Stemski	69

Stepens	80

Stračárová	10, 66

Stuckenschneider	9, 60

Svobodová	12, 66, 71

Tałach-Kubas	2, 6, 95

Talar	7, 59

Tan	62

Tchelet	68

Tisserland	93

Tota	84

Trabelsi	73

Tropea	61

Tuomola	53

Valatkeviciene	76

van Haren	59

Verghese	22

Vints	7, 55

Voelcker-Rehage	6, 21, 24, 34, 35, 36, 37

Vogel	6, 23

Vogt	23

Völker	62

von Holdt	36

Wilk-Głodzik	11, 92

Wollesen	8, 24, 25, 33, 37

Wolter	12, 69

Yekutieli	68

Zieschang	86

Zijlstra	2, 7, 51

Ziv	81

Zwingmann	9, 24, 37
