## Additional file 4 for "Planning and Conducting an Online Conference at the time of COVID-19: Lessons Learned from EGREPA 2021"

**CHECKLIST**

- Choose a white, neutral background (please do not use a virtual background).
- Lighting should be consistently bright (e.g. your face should be lit from the front).
- Avoid a window in the background.
- Make sure there is no background noise.
- The camera should be at eye level (for example, put a large book under it).
- Use in-ear-headphones with an integrated microphone for a higher audio quality.
- Choose a comfortable sitting position and adjust your camera image so that your head and shoulders can be seen centrally.

**PRACTICAL TIPS**

- The less you move your upper body during the presentation, the better.
- Speak directly to the microphone and look into the camera as often as possible.
- Do not touch your microphone.
- Speak clearly and slightly slower than usual (presentation pace).
- Don't go past the last slide - instead of ending the presentation, stay on the last slide.
