## Additional file 5 for "Planning and Conducting an Online Conference at the time of COVID-19: Lessons Learned from EGREPA 2021"

**USER-GUIDE (POWER-POINT)**

In order to pre-record your presentation, we strongly recommend to use PowerPoint. How to proceed is explained step-by-step, below.

Step 1

Open the finished PowerPoint presentation from which you want to create the video. Please select, "*Slide Show*" in the menu at the top. Afterwards, click on "*Record slide show*" in the toolbar at the top.

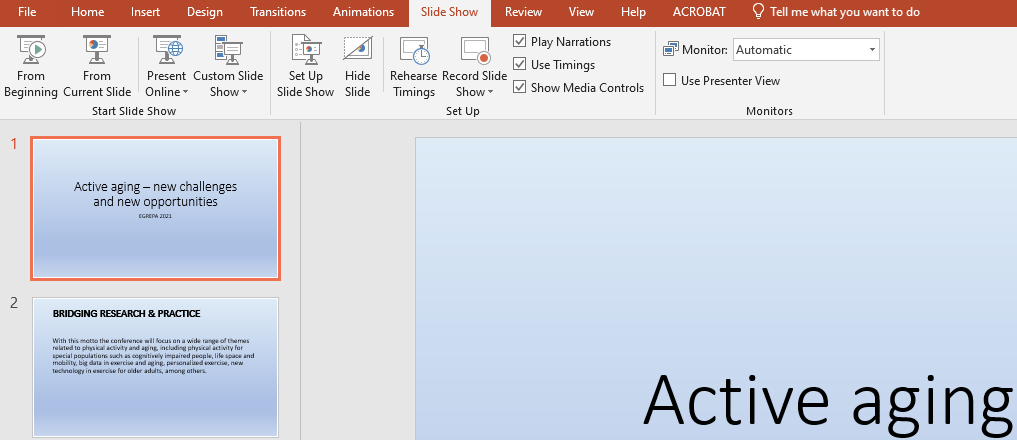

Step 2

Now you can choose to record audio or video for the respective slide by selecting the red "*Record*" button at the top on the left side. If you want to end your recording, simply click on the “*Stop*” button next to it. To view or listen to your recording, select the “*Replay*” button. If you want to discard your previous recording and start a new one, click on "*Clear*” (current slide or all slides) in the bar at the top.

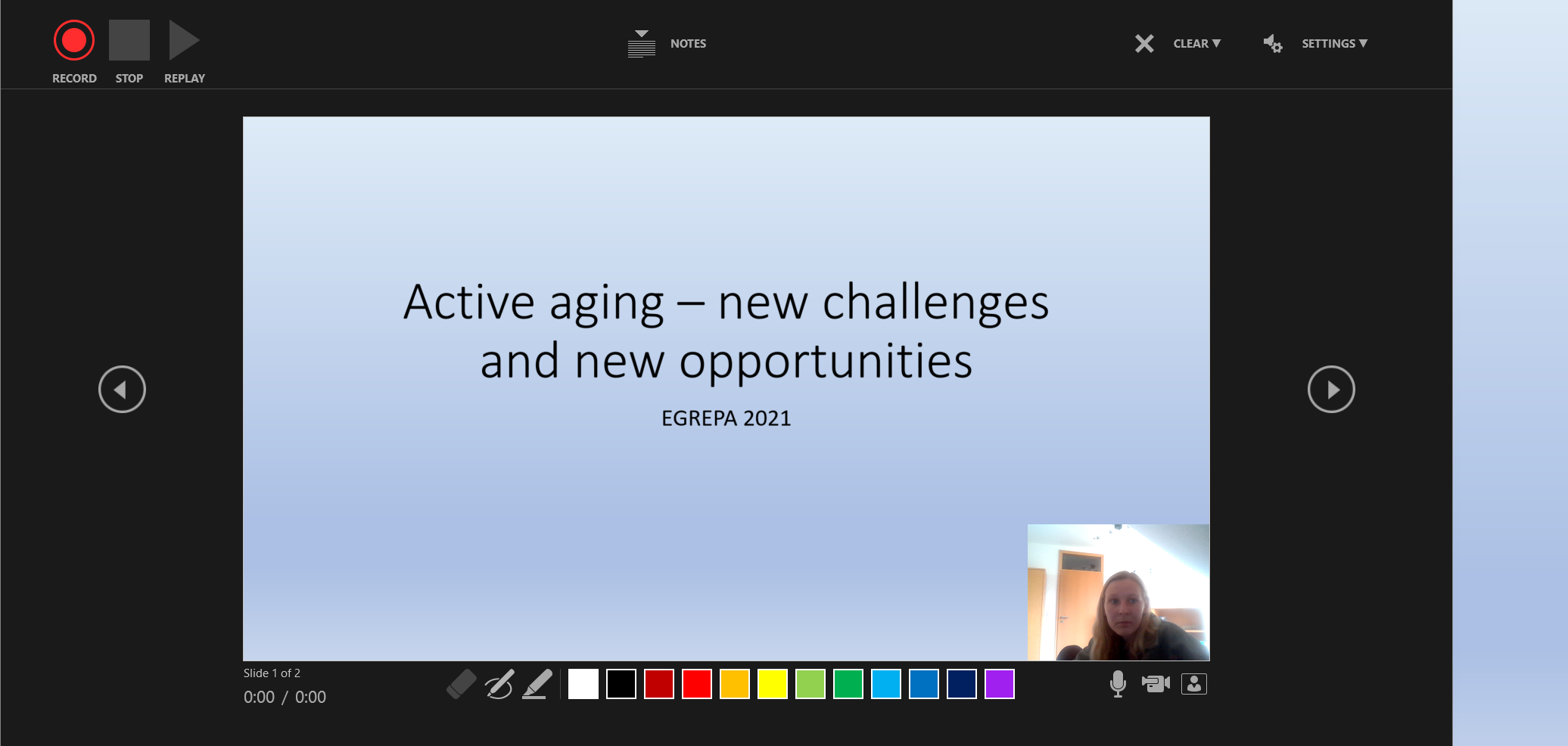

Video

Step 3

Save the presentation in ppt format, just like any other PowerPoint presentation. This allows you to make changes to it at any time.

Step 4

Select in the bar at the top the button “*File*” and then choose “*Export*” and choose beside *“Create a Video*”.

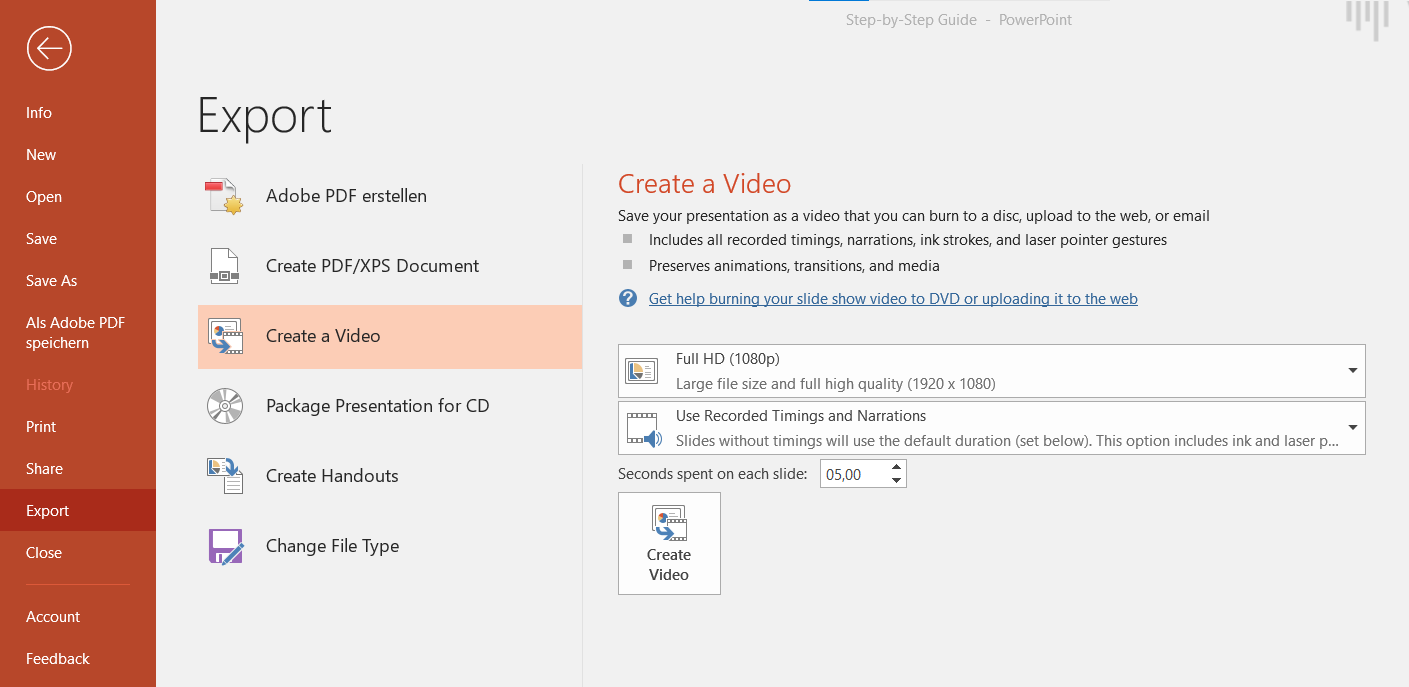

Step 5

Choose, if possible, the “*Full HD*” option and make sure you select “*Use Recorded Timings and Narrations*”. Please finally “*Create Video*”.

Step 6

After you have clicked on "Create video", in the following window you only have to determine in which folders you want to save your video from PowerPoint. You can also select the file type in which you want to save your PowerPoint video. Please choose “*MPEG4 format*”, because it has the advantage that it runs on a larger number of devices and systems.
