## Additional file 6 for "Planning and Conducting an Online Conference at the time of COVID-19: Lessons Learned from EGREPA 2021"

**USER-GUIDE (INDICO)**

In order to upload your pre-recorded presentation video, please follow the two steps below.

Step 1

Please login with your Indico Account. Choose in the left sidebar the button “My Conference”, select “My Contributions” and click on your contribution.

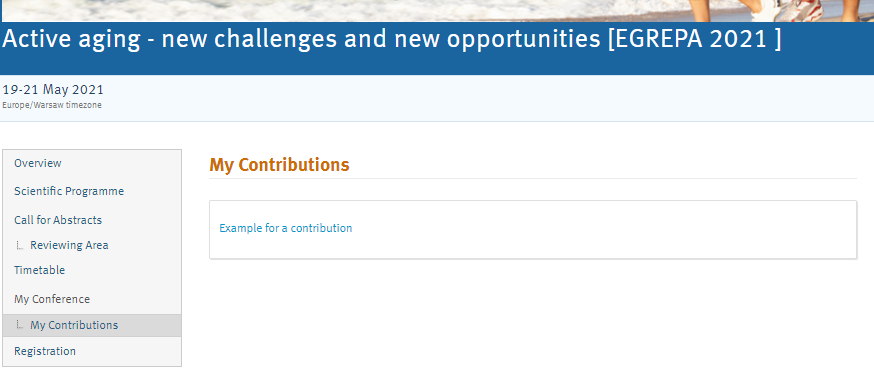

Step 2

Afterwards, you see details to your contribution. Select the “*pen*” in the section “*Presentation materials*”. Finally, please configure the “Protected” mode. Please upload the video with the presentation title and your surname.

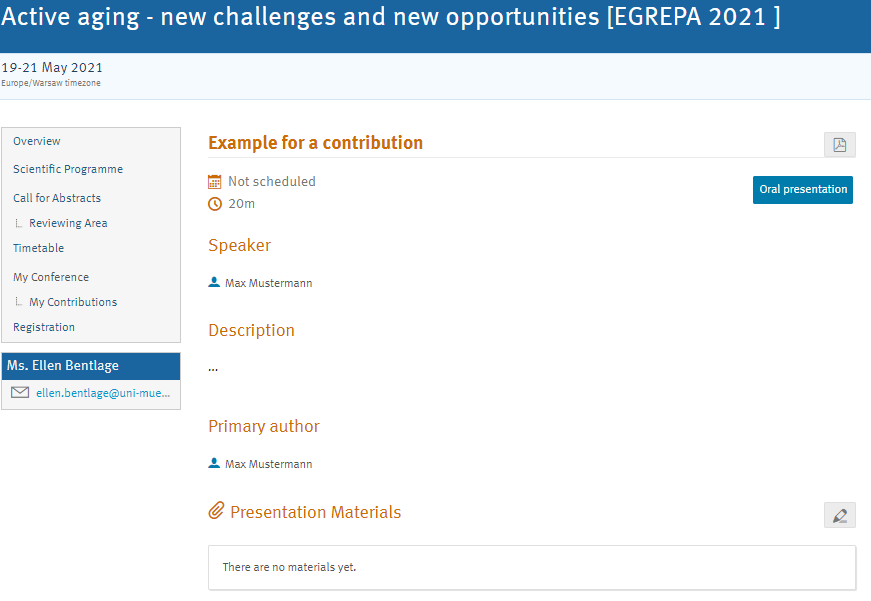
