## Additional file 7 for "Planning and Conducting an Online Conference at the time of COVID-19: Lessons Learned from EGREPA 2021"

**GATHER INTRODUCTION VIDEO**

• Please click on [this link](https://wwuindico.uni-muenster.de/event/162/attachments/54/616/Gather%20Instruction%20Video.mp4) to access the introduction video to Gather.

**GATHER HANDOUT**

What is Gather?

A platform that allows participants to move freely it´s avatar using the arrows of the computer keyboard and interact within a virtual 2D space. When avatars approach one another, participants can talk to each other through a video call, launched automatically. As in real life, participants are free to choose where to go and with whom to speak, exchanging a few words with one, engaging in a longer conversation with another or just deciding to move away if they wish. Interactive objects such as videos can be accessed when the avatar is near the object. We welcome you to explore the space, to meet new colleagues, or plan a collaboration with old ones.

Gather as part of our EGREPA 2021 Online Conference

For the Virtual EGREPA 2021 conference you have the following areas where you can gather from May 19th, 2021 through May 21, 2021:

• The main area/ lobby (open space and private rooms).

• Poster Session Rooms (for poster discussions).

• Lounge with a warm atmosphere (for private conversations).

**(!)** Please note that these rooms are for poster discussions and casual meetings, all other sessions will be held via the YouTube Livestream & Zoom.

What you need

• A desktop/laptop with a mic and camera.

• A web browser - Chrome or Firefox are recommended (mobile phones & tablets cannot be used).

• Check if your browser permitted camera and microphone access.

• We strongly recommend using headphones.

If you have technical difficulties

• Refreshing the page will fix most things.

• If that doesn’t work, try muting and unmuting your mic and camera.

• Check if your browser permitted camera and microphone access.

Start

Before joining the space, you need to enter your name and create your own avatar:

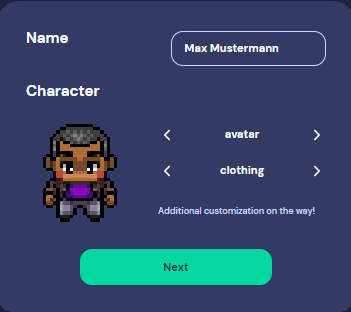

Features

• Open an interactive object by clicking “X” on your keyboard.

• There is a locate feature to find others by clicking their name in the participant panel.

• There is a messaging feature that allows you to message people in four ways:

1. Individually by clicking on their name in the participant panel.
2. Locally to the people you are video chatting with.
3. Room chat with all the people in the current room you are in.
4. Globally to all the people in your map.

• Want to full screen someone else’s video? Just click on their video.

Explanations of the Icons

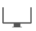
Screen sharing ability

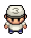

Change your avatar character and clothing

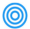

Change your interaction distance

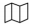

Mini map to preview the space you’re in

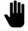
Raise hand feature: good for Q&A in larger discussion groups

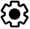
Opens the settings menu
