## Additional file 8 for "Planning and Conducting an Online Conference at the time of COVID-19: Lessons Learned from EGREPA 2021"

**Q 14: What were positive aspects of the online conference?**

| Category | Statement (N=29) |
| --- | --- |
| Scientific | - Constructive questions. - High quality research, as well as the chance to an open discussion through Zoom and Gather. - Discussions after each presentation. - I enjoyed each presentation and discussion. - The conference was interesting and enlightening. - Open and interesting discussions. - Modern, great researchers, high quality posters. - Diverse projects. |
| Technical | - Extremely well run, especially the transition between the different applications (YouTube, Gather, Zoom). - Technically, the program went well. - Successful use of alternative technical solutions to realise the conference / despite it being virtual: still the feeling of a shared conference visit. - Gather was fun and a great alternative, to discuss the conference content with others. |
| Social | - We can safely meet in a large group of participants. Each person could choose the most interesting presentations for her/him and not dedicate time to less interesting. - Safety, meeting researchers from all over the world, who run projects about active aging. - Meeting people and talk with them. - You could get to know a lot of people and watch all the sessions. - Various participants from different countries. - In Gather, I met a lot of new people. - Possibility to talk to a lot of people or chat with them. |
| Organisational | - Digital organisation was great. Easy to follow. - Organisation, physical activity/relaxed-breaks. - I really appreciate pre-record presentation, because of stress limitation. - It was easier to hold the timeline, than in stationary conferences. - To make it possible to conduct the conference, even though the pandemic is still ongoing. - Very good organisation. - The communication and attention provided by the organisers for any questions. - It was well organised and had a clear structure. Besides, it was taken care of the schedule and the information/ instructions given in advanced were very helpful. - Great organisation. - Well filled and structured programme. When I first wanted to write that I missed longer breaks, the structure allowed to skip some parts that you considered less interesting. Also, everything could be re-watched. - Thank you for the well-organised conference. - The communication and attention provided by the organisers for my questions. |

**Q 15: What were negative aspects of the online conference?**

| Category | Statement (N=29) |
| --- | --- |
| Scientific | / |
| Technical | - Negative was sometimes the bad audio. - Limited integration and access to talk longer to presenters. - To switch from YouTube to Zoom and vice versa. - I had problem to get to Gather with link. - At the beginning a bit confusing Zoom/YouTube, why not solely use Zoom. |
| Social | - It would have been great to jest up in person, not online. It was also hard to spend so much time in front of the computer. - Lack of direct contact. - None, but I prefer in person. - Less networking possibilities than a meeting in person, but still great. - In general, I think online conferences cannot compete with personal meetings, especially, discussion-wise online conferences are limited. Normally, breaks after sessions allow for quick exchanges, that cannot be deepened in the evenings or during social events and help to build collaborations and exchange ideas better. - It was impossible to meet directly in Krakow. |
| Organisational | - Too many tabs open. - Just listen to pre-record presentations instead of live presentations. - Need more & longer breaks – very tiring in front of the screen all day. - It was sometimes hard to understand what will take place in which platform. - For me it was a little bit hard to sit next to the computer for long periods of time. - It was a pity that Gather was not used more often. I saw always the same persons on Gather. Most of the time only about 7 participants were online. |

**Q 15: How can we improve the next online conference?**

| Category | Statement (N=19) |
| --- | --- |
| Scientific | / |
| Technical | - Zoom only. - Just keep one platform for next congress or conference. - Motivate the participants to turn on the video. - Keep interaction possible without videos on YouTube. Maybe everything could have been done in one platform like Zoom. - It should be recommended to pay more attention to presentation on YouTube. In many cases there were problems to read the text. |
| Social | - More opportunities for exchange in smaller groups of interest just after presentations. - Maybe more Gather time, but I hope we could attempt and meet physically next year. - I would like to have more time in Gather to discuss poster and presentation. |
| Organisational | - More breaks; Maybe to add one longer break in full day lectures. - Longer breaks and time for gathering after each session. - Live talks. - Maybe reduce sessions, in which the presentations are streamed simultaneously on YouTube to sessions only for discussion and pre-uploading the videos for everyone beforehand. The speakers could have 5 minutes of talking time to reintroduce the presentations to the audience, before discussion starts. - It would have been nice if Gather could have played a bigger role. For example, that you would need to walk into the main conference room to participate in the discussion, e.g. walking into this room would allow you to press x to open Zoom. This way, during breaks everyone listening would be in the same room, which would decrease the threshold to interact with each other during breaks. - It would be nice to have possibility to see oral poster presentations in Gather. |
